# VARIATIONS IN THE ULTRAVIOLET FLORAL PATTERNS AND POLLINATOR PREFERENCE AMONG SELECTED NON-INVASIVE AND INVASIVE PLANTS OF TAMIL NADU, INDIA

**DOI:** 10.1101/2023.06.27.546802

**Authors:** Indhar Saidanyan Ravichandran, Parthiban Balasingam, Mohan Raj Rajasekaran, Karthikeyan Ananthapadmanabhan, Manojkumar Muthuvel, Kottaimuthu Ramalingam, Vigneshwaran M, Kamaladhasan Nalluchamy, Saravanan Soorangkattan, Anbarasan MR, Chandrasekaran Sivagnanam

## Abstract

Fossil evidence shows that pollinator-mediated plant reproduction evolved ∼140 million years ago and bee facilitated pollination evolved ∼70 million years ago. Human vision is limited to the visible color range of 400 to 750 nanometres, whereas most pollinators can perceive the ultraviolet (UV) range in addition to visible colors. Bees have been reported to have highest spectral sensitivity in the ultraviolet spectrum. The main objectives of the study were (1) to assess the prevalence of ultraviolet floral patterns, (2) to analyse floral patterns in relation to plant-pollinator interaction among invasive and non-invasive plants, and (3) to test for intraspecific floral pattern variations among plants with different flower color morphs. A study was conducted on 188 plant species (80 invasive and 108 non-invasive) from parts of Western and Eastern Ghats region of Tamil Nadu, India. The flowers of the studied plant species were imaged in ultraviolet (320-380 nm) and visible spectrums. The mode of pollination for the selected species were documented and confirmed with existing literature. The intraspecific variations in the floral patterns among flower color polymorphic plant species (N=10) were documented in ultraviolet and visible spectrums. Among the studied plant species, around 58% had discernible floral patterns when observed in the UV spectrum, whereas the rest were observed to completely absorb or reflect UV radiation. Whereas 46% of the studied plants exhibited no pattern in the visible spectrum. A significant difference was observed in the pollinator relationship among the ultraviolet floral patterns in invasive plants (*χ2* = 63.98, *df* = 32, *P* < 0.001), whereas no significant variation was evidenced in the pollinator relationship among the ultraviolet floral patterns in non-invasive plants (*χ2* = 19.50, *df* = 24, *P* = 0.724). Analysis of pollinator preference revealed that invasive species were mostly pollinated by bee and butterfly mediated pollination, whereas non-invasive species were mostly pollinated by bees and generalist insects. Intraspecific variations in the floral ultraviolet signal were observed among different morphs in a few flower color polymorphic species, especially in *Lantana camara*. The multispectral analysis of floral patterns revealed that plants utilize both the visible and ultraviolet spectrums to effectively communicate with pollinators. The results from the present study strongly suggest that the variation in the floral ultraviolet signature among invasive species might play a vital role in plant-pollinator interaction and invasion success.

## Introduction

Estimates show that pollinator-mediated plant reproduction might have evolved around 140 million years ago, and bee facilitated pollination in flowering plants might have evolved around 70 million years ago (Wojcik & Wojcik, 2021). The co-evolution of pollinators and plants has led to the evolution and diversification of specific plant reproductive traits to effectively communicate with pollinators to ensure reproductive success (Briscoe & Chittka, 2001; Wojcik & Wojcik, 2021). Plants have been found to employ visual, scent, tactile, acoustic, and gustatory signals to attract pollinators (Raguso & Willis, 2002). The visual signal provides contextual information on the location and availability of the flower even from a longer distance, while the rest of the signals are more localized (Milet-Pinheiro et al., 2015; Raguso & Willis, 2002). Visual cues play a vital role in guiding pollinators to flowers (Briscoe & Chittka, 2001; Kelber et al., 2003; Young et al., 1991).

Humans perceive wavelengths in the range of 400 to 750 nanometers as specific hues of colors such as blue, green, and red (Boynton, 1990). The majority of pollinators, including bees, butterflies, birds, and other insects, are capable of perceiving the ultraviolet range of the spectrum (Cuthill et al., 2000; Kelber et al., 2003; Shrestha et al, 2013a; van der Kooi et al., 2021), and bees have been reported to have the highest sensitivity in the ultraviolet spectrum (Chen et al., 2020; Davies, 2017; van der Kooi et al., 2021). With reference to the human perceptible spectrum, studies on plant-pollinator relationships have yielded only a partial understanding of their relationship. Until very recent developments in digital photography, only a very few methods were available to visualize and understand ultraviolet floral cues-based communication with pollinators (Garcia et al., 2014; Napoleone et al., 2022). The recent understanding of the plant traits expressed in the ultraviolet spectrum has paved the way for a deeper understanding of plant-pollinator communication and the environment (Chen et al., 2020; Dyer et al., 2012; Koski, 2020; Ödeen & Håstad, 2010).

Most of the UV radiation, especially all the germicidal UV-C radiation in the wavelength range of 100–280 nm and most of the UV-B (280–315 nm), is blocked by the naturally occurring ozone layer (McKenzie et al., 2003). The UVA radiation that penetrates through the atmosphere is biologically active (Godar et al., 2009). Pollinator visual cues are color, markings, shapes, and guides on flowers that visually guide and attract pollinators to flowers (Lucas-Barbosa et al., 2016; Young et al., 1991). Plants have been documented to exhibit different floral patterns through pigments that either absorb or reflect ultraviolet radiation like the visible color patterns (Chittka & Menzel, 1992; Gronquist et al., 2001; Kevan et al., 1996; Koski & Ashman, 2014; Lunau, 1992; Papiorek et al., 2016; Primack, 1982; Utech & Kawano, 1975). A recent study on flowering plants in the Neotropical Savanna documented the different patterns exhibited in ultraviolet and visible wavelengths for pollinator communication (Tunes et al., 2021). Visual pollinator cues can be produced by differential color patterns or by altering the wavelength of light with physical structures on the surface of the flower (Moyroud & Glover, 2017). Visual flower colors and cues help in flower identification by providing striking contrast from the green vegetation (Kevan et al., 1996). Nectar guides and pollen marking are some of the specialized pollination cues that guide the visiting pollinator to nectar and pollen, thereby conserving pollinator energy and increasing pollination efficiency and constancy (Hansen et al., 2012; Leonard & Papaj, 2011).

Multi-seasonal flowering and flower color polymorphism have been reported to be prevalent among invasive plant species when compared to native or exotic species (Saidanyan et al., 2016). Invasive species have not only been more efficient at attracting pollinators but also have been documented to affect the existing relationship between native plants and their pollinators at the community level at invaded sites (King & Sargent, 2012; Parra-Tabla & Arceo-Gómez, 2021). Invasive plants were found to attract various suites of pollinators and longer floral availability, which translates to better reproductive success (Keller et al., 2011), thereby gradually displacing native plant populations. Invasive plant species have been found to have higher floral constancy in comparison to the native plant species in the invaded areas (Albrecht et al., 2016). Invasive species, in particular, have been observed to be more efficient at attracting pollinators and better at integrating into an introduced ecosystem (Anu et al., 2022; Vilà et al., 2009).

The association of flower color with their exhibited pollinator cues in invasive species may aid pattern recognition in pollinators, thereby increasing the odds of successful pollination (Albrecht et al., 2016). The floral spectral reflectance of invasive species has been found to be more prominent in the ultraviolet region in comparison to non-invasive species (Sooraj et al., 2019). Researchers have addressed the invasive plant-pollinator interaction with reference to the visible spectrum (Beans & Roach, 2015; Gibson et al., 2012; Goodell & Parker, 2017; Narbona et al., 2021; Saidanyan et al., 2016; Trunschke et al., 2021; Yan et al., 2016); whereas there has been a lacuna in understanding the impact of ultraviolet patterns and pollinator association among invasive species. Consequently, this study was conducted to examine the prevalence of ultraviolet patterns in invasive and non-invasive species of flowering plants, to comprehend their relationship with pollinators, and to evaluate the effect of the ultraviolet signature on invasive success.

## Methodology

### Study sites

The research was conducted in the areas surrounding the Western Ghats region, which includes the Agasthiyarmalai Hills, Andipatti Hills, and Palani Hills, and the parts of the Eastern Ghats region, which include the Kararandhamalai Hills and Sirumalai Hills (Figure 1). The Western Ghats, one of the world’s biodiversity hotspots, is known for its species richness and diversity (Karanth et al., 2016). Previous reports have confirmed the pervasiveness of invasive species in the Western Ghats region of Tamil Nadu (Aravindhan & Rajendran, 2014; Chandrasekaran & Swamy, 1995, 2002, 2010; Swamy et al., 2000), suppressing the endemic plant species population in the region (Balaguru et al., 2016). In the study sites, the flower color characteristics of invasive and non-invasive terrestrial angiosperm species, as well as their pollinator associations, were photographed and recorded. Plant sampling at each study site was done as described by Tunes et al., 2021; field observations and data collection were done over a period of one year (July 2021–July 2022). About four 25-m^2^ quadrats were randomly placed at each study site, and all the plants with open flowers during the time of observation were selected for the study. The study sites were geographically located using a handheld GPS recorder (Garmin eTrex^®^ H).

**Figure 1:**
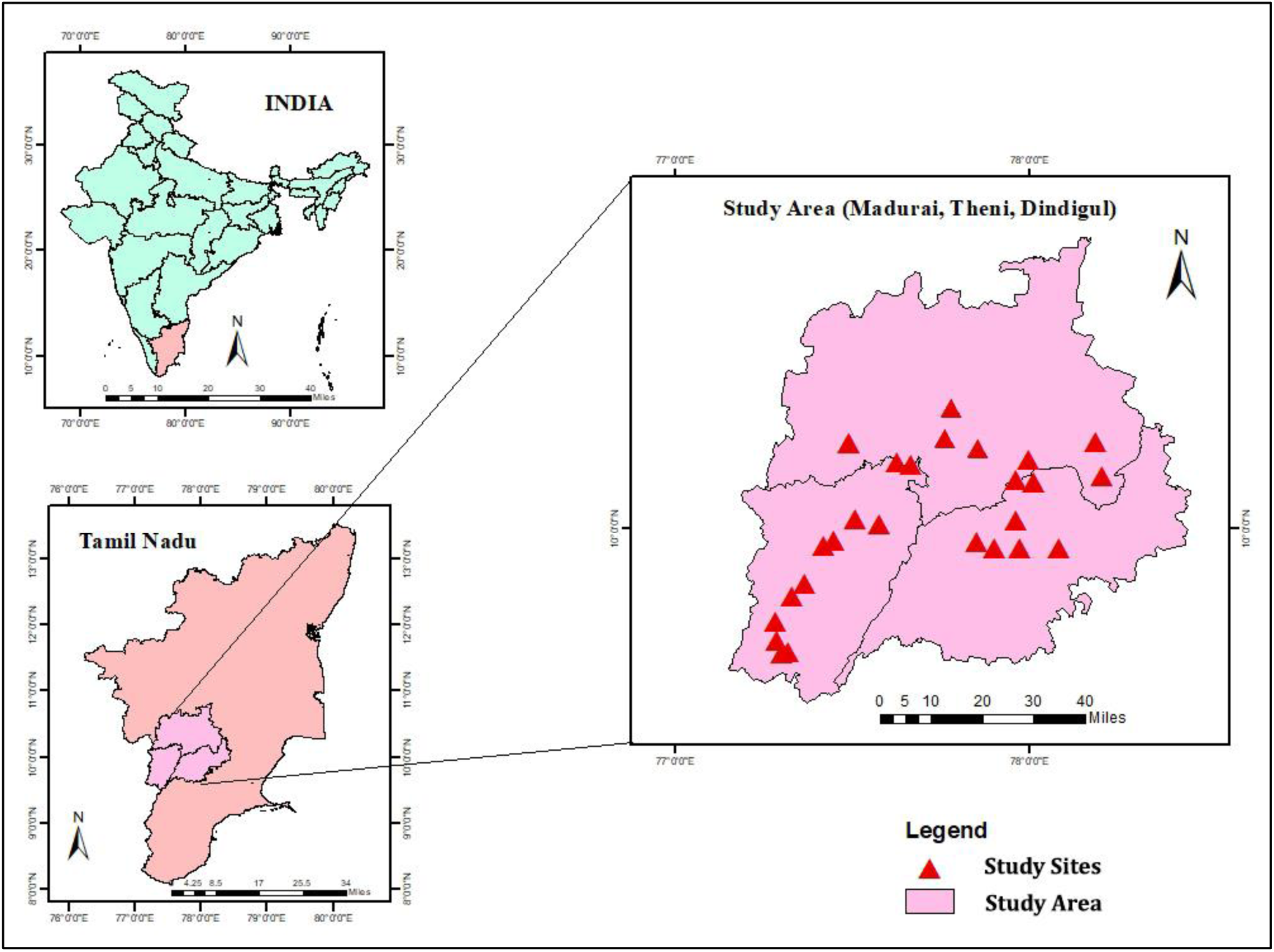
Map of the study site areas in the Western Ghats (Agasthiyarmalai Hills, Andipatti Hills and Palani Hills) and Eastern Ghats (Kararandhamalai Hills and Sirumalai Hills) regions of Tamilnadu, south India.

### Multi-spectral imaging

To determine the various floral patterns exhibited in the visible spectrum (400–750 nm) and ultraviolet A spectrum (315–400 nm) by the studied non-invasive and invasive species of terrestrial flowering plants, multispectral imaging was carried out at the field sites. Standardized techniques were adapted for plant imaging in the visible and ultraviolet spectrums (Koski & Ashman, 2016; Napoleone et al., 2022; Schulte et al., 2019). Visible-spectrum imaging was done with a Sony A6000 mirrorless camera equipped with a 16–50 mm F3.5–5.6 variable lens. The color profile for the visible color images was set to “neutral” to accurately represent the floral colors. The Godox TT520 II flash was used to supplement the visible image’s uniform illumination. The highest possible aperture (F3.5) was preferred with the given lens, provided there was adequate illumination from the flower and sunlight conditions. The visible camera was programmed in aperture priority mode, and the shutter speed and ISO were set to “Auto.” The image color profile was calibrated for color accuracy using a Pantone® X-Rite Color Checker Passport Photo 2. White balancing was done using virgin PTFE tape as a white reflectance standard (99%). Modern digital camera sensors and lenses are treated with a special coating that specifically reflects ultraviolet (UV) radiation (Napoleone et al., 2022). Similarly, a specialized lens that is transparent to UV light along with a bandpass filter that only allows UV wavelengths of light was required. A full spectrum converted Sony A6000 mirrorless camera equipped with a Novoflex Noflexar 35 mm F3.5 prime lens attached to a M42 to NEX/M mount was used. The 2” Baader U-Venus filter with a transmission peak at 350 nm and a 30 nm bandpass was used to specifically capture UV images in the UV-A spectral range of 320–380 nm. Artificial UV illumination was required for imaging as most of the ultraviolet radiation from the sun is blocked by the Earth’s ozone layer or refracted by the atmosphere. A full-spectrum converted Nikon SB–700 flashlight was used to provide adequate UV illumination for the floral subject. The converted camera, along with the specialized lens and UV filter, forms the UV imaging setup (Figure 2) required to capture floral images. White balancing was performed with virgin PTFE tape (a Lambertian reflectance surface) with a specific UV reflection of 99%. The UV camera was set to aperture priority, and the shutter speed and ISO were set to “Auto.” Both visible and ultraviolet imaging was done in natural field conditions and supplemented by appropriate lighting for the different spectra of observations.

**Figure 2:**
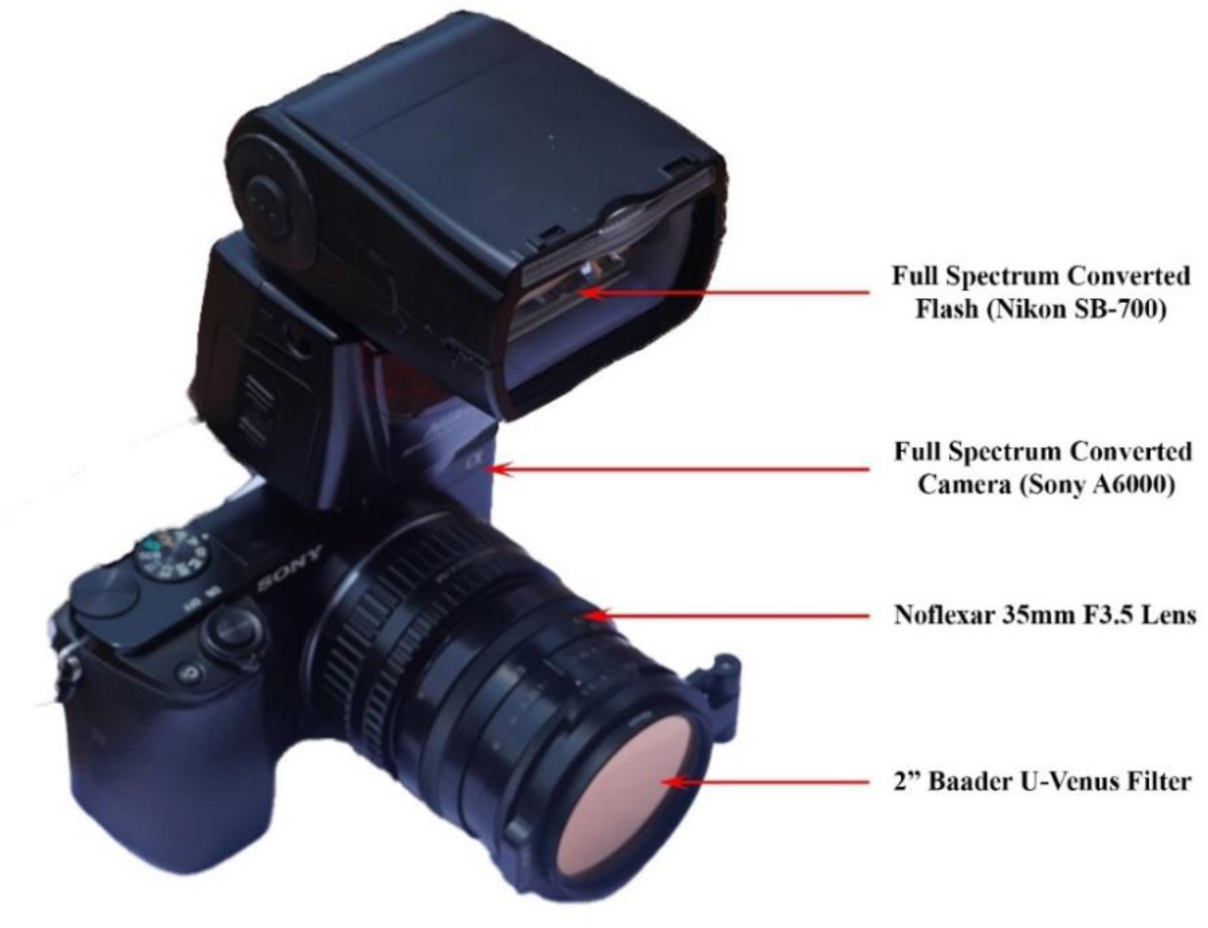
Ultraviolet imaging setup with full spectrum converted Sony A6000 camera equipped with Noflexar 35mm lens along with screw mounted 49mm Baader-U filter.

### Image analysis

The methodology for plant floral image interpretation was adapted from recent literature (Tunes et al., 2021). The floral patterns were categorized into four different categories for the visible spectrum observations and five different categories for the ultraviolet spectrum observations. The visible floral color patterns were categorized as follows: i) Patternless (VIS-P), ii) Bullseye (VIS-BE), iii) Contrasting Corolla (VIS-CC), and iv) Contrasting Reproductive Structures (VIS-CR) (Figure 3). Similarly, classification of floral color patterns in the ultraviolet spectrum is as follows: i) Patternless (UV-P), ii) Absorbing (UV-A), iii) Bullseye (UV-BE), iv) Contrasting Corolla (UV-CC), and v) Contrasting Reproductive Structures (UV-CR) (Figure 4). The patternless flowers in both the visible and ultraviolet spectrums (VIS-P and UV-P) were perceived to be bright, reflecting, and uniform without any discerning patterns in their respective spectrums. The ultraviolet absorbing (UV-A) pattern was perceived to be a uniformly dark flower without any discernible pattern or differentiation from the background in the UV spectrum. The floral bullseye pattern was perceived as having a dark central region (comprised by the central corolla region and reproductive structures) with a brighter peripheral region in both spectrums (VIS-BE and UV-BE) of observation. The contrasting corolla pattern in both spectrums (VIS-CC and UV-CC) was perceived as contrasting regions in the corolla that differ in brightness or hue (visible) and most frequently serve as a nectar or pollen guide to pollinators. The contrasting reproductive structure pattern under both spectrums (VIS-CR and UV-CR) of observation was perceived to have contrasting reproductive structures (darker or lighter) with reference to the corolla. In post-processing, the UV images were split into red, green, and blue channels, in post-processing, and only the “red” channel data was considered for the study as it held the most information in the UV spectrum (Koski & Ashman, 2016; Tunes et al., 2021).

**Figure 3:**
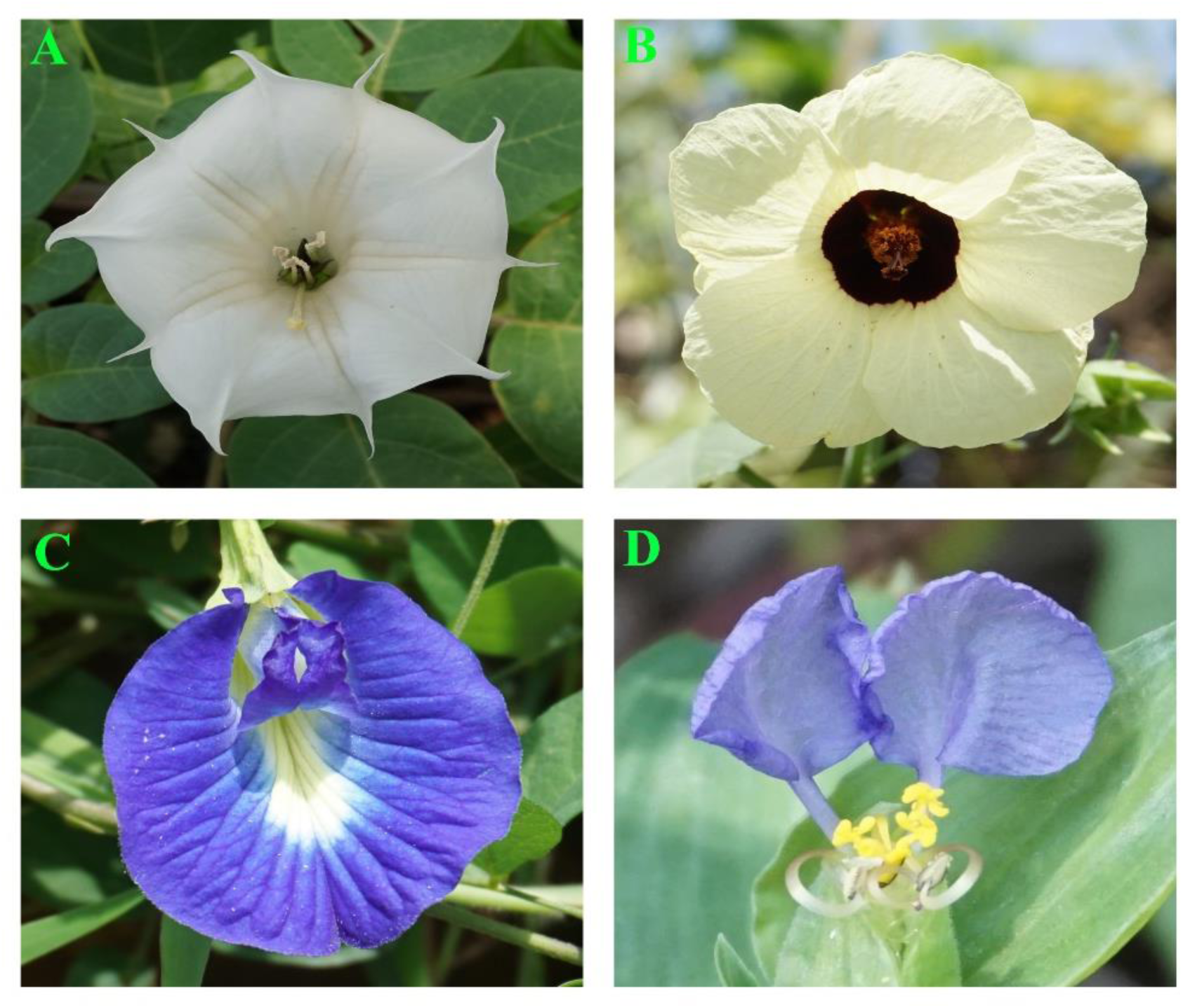
Representative images of different visible color floral patterns observed in the study sites. The different patterns observed are as follows: (A) Patternless (*Datura innoxia* Mill.); (B) Bullseye (*Hibiscus vitifolius* L.); (C) Contrasting Corolla (*Clitoria ternatea* L.) and (D) Contrasting Reproductive Structures (*Commelina benghalensis* L.).

**Figure 4:**
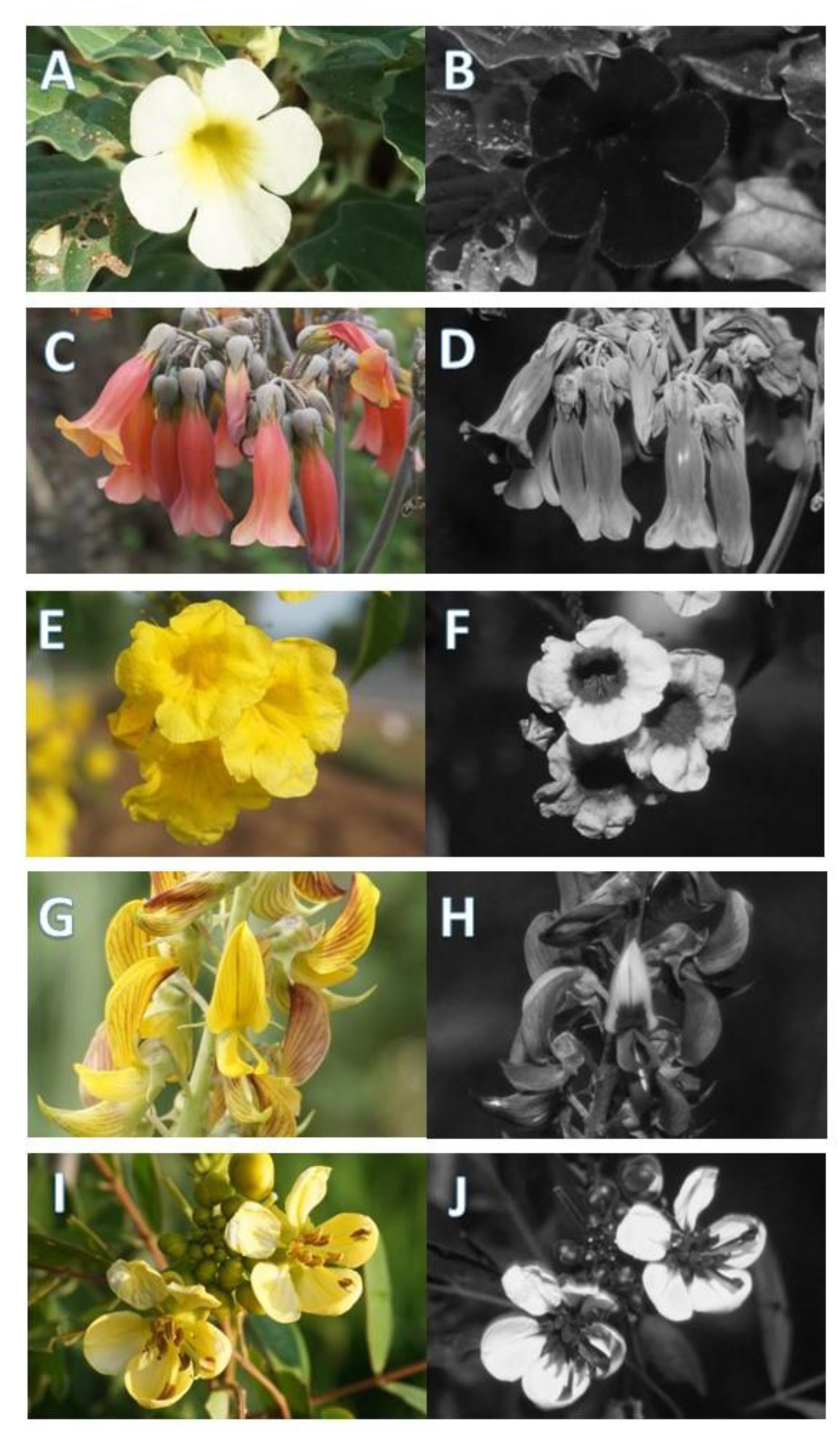
Representative images of different ultraviolet patterns observed in the study sites. The different patterns observed are as follows: (A&B) UV Absorbing (*Pedalium murex* L.); (C&D) UV Patternless (*Kalanchoe delagoensis* Eckl. & Zeyh.); (E&F) UV Bullseye (*Tecoma stans* (L.) Juss. ex Kunth); (G&H) UV Contrasting Corolla (*Crotalaria pallida* Aiton), and (I&J) UV Contrasting Reproductive Structures (*Senna siamea* (Lam.) H.S.Irwin & Barneby).

### Floral trait dataset

The plants studied in the field sites were taxonomically identified with the help of standard taxonomic keys (Matthew, 1981), and any ambiguity that arose with identification was clarified with plant taxonomists at the Botanical Survey of India. The flower colors displayed by plants were interpreted from visual observations and from captured images. The perceived flower colors were then grouped into three categories based on the spectral sensitivity of human eyes (Schnapf et al., 1987), such as: i) the VIB group; ii) the White group; and iii) the YOR group. The VIB flower color group includes flower colors in the violet, indigo, and blue ranges, which correspond to the peak sensitivity (420–440 nm) of human cone cells at the short wavelength. The YOR flower color group consists of flower colors in the yellow, orange, and red range, which corresponds to the peak sensitivity (560–580 nm) of human cone cells in the long wavelength. The White flower color group comprises white, pale white, greenish-white, yellowish-white, and pinkish-white colors. The pollinators associated with each species were determined based on the most frequent pollinator observed during the study. Plants with missing pollinator information were substantiated with data from published literature, databases (Flowers of India, 2022; KFRI - KFSTAT Database, 2022; TRY Plant Trait Database, 2022), and books (Matthew, 1981). The primary pollinator associations in the studied plant species were categorized as follows: i) Bee-mediated pollination (Melittophily), ii) Non-specific insect-mediated pollination (Generalist), iii) Both bee and butterfly-mediated pollination (Melittophily & Psychophily); iv) Bird-mediated pollination (Ornithophily), v) Butterfly-mediated pollination (Psychophily), vi) Self-pollinating plants and self-fertile flowers (Autogamy), vii) Hawkmoth-mediated pollination (Sphingophily), viii) Wind pollination (Anemophily), ix) Moth-mediated pollination (Phalaenophily), and x) Bat-mediated pollination (Chiropterophily). The invasive status of the studied plant species was confirmed with reference to the existing invasive plant databases and compendiums (CABI Compendium Invasive Species, 2022; GISD, 2019). All the native and exotic plant species in the study have been classified together as non-invasive plants (Sooraj et al., 2019). A detailed list of all the studied plant species along with their flower color, invasion status, ultraviolet pattern, visible pattern, flower color grouping, and associated pollinator is given in Supplementary Table 1.

### Phylogenetic analysis of UV patterns and invasive status

A phylogenetic tree was constructed for the studied species using the V.PhyloMaker (Jin & Qian, 2019) package developed for the R programming language. The phylogenetic signal of each species was calculated using the ultraviolet categorical variables (Borges et al., 2019). The open-source package uses data from published phylogenetic mega-trees, holds data on 74,533 vascular plants across 10,587 genera. The phylogenetic tree is constructed based on the existing nuclear sequences such as internal transcribed spacer (ITS), external transcribed spacer (ETS), and chloroplast (trnL-F) available for each of the studied species. The specific UV floral pattern along with invasive status for each species were mapped in the phylogenetic tree construct.

### Statistical analysis

All statistical analyses were carried out using OriginPro (Origin Pro, 2021) and the R programming language (R Core Team, 2022). The plant trait data collected for the studied species were grouped into two or more variables, thereby Pearson’s chi-squared test (χ ^2^) was found to be ideal for comparison.

#### Pearson’s Chi-square

The Pearson’s chi-square test was employed to test if any observed differences between the plant trait categorical variables arose by chance. All the plant trait data categories were tested for its independence with their reported invasiveness of the plant species.

#### Binomial logistic regression

The plant trait data were subjected to logistic regression (binomial) to predict the odds of invasion in relation to the categorical variables such as flower color group, pollinator association, visible pattern, and UV pattern. The invasiveness of the studied plant species was considered the dependent variable, whereas the individual plant traits were considered independent variables for the analysis.

## Results

The floral color patterns exhibited by flowers were imaged in the visible (VIS) and ultraviolet (UV) spectrums among 188 flowering plant species belonging to 46 families. The floral patterns were categorized based on their appearance in the visible and ultraviolet spectrums. The VIS and UV images of all the 188 studied flowering plants are displayed in Supplementary Figure 1.

### Floral pattern distribution in the visible and ultraviolet spectrum

Observations on the distribution of perceived floral patterns in the VIS spectrum showed that patternless flowers (VIS-P) were the most abundant (46%), followed by contrasting reproductive structures (VIS-CR; 24%), contrasting corolla (VIS-CC; 20%), and bullseye (VIS-BE; 10%) among the studied species. Analysis of the distribution of perceived floral patterns in the UV spectrum showed that absorbing (UV-A) was most abundant (37%), followed by bullseye (UV-BE; 26%), contrasting reproductive structures (UV-CR; 20%), contrasting corolla (UV-CC; 12%), and patternless (UV-P; 5%) among the studied species. Patternless flowers were higher in the visible (VIS) spectrum than the ultraviolet (UV) spectrum. There were significant variations in bullseye, contrasting corolla, and contrasting reproductive structure patterns among the visible (VIS) and ultraviolet (UV) spectrums in the studied plants (Table 1).

**Table 1:** Distribution (%) of visible (VIS) and ultraviolet (UV) floral patterns in the studied plant species (N = 188).

| **Patterns** | **Visible (VIS)**<br>**Spectrum (%)** | **Ultraviolet**<br>**(UV) Spectrum**<br>**(%)** | $\chi^2$ | *df* | *P* |
| --- | --- | --- | --- | --- | --- |
| Patternless (P) | 46 | 5 | 107.61 | 12 | < 0.001 |
| Absorbing (UV-A) | NA | 37 |  |  |  |
| Bullseye (BE) | 10 | 26 |  |  |  |
| Contrasting Corolla<br>(CC) | 20 | 12 |  |  |  |
| Contrasting<br>Reproductive<br>Structures (CR) | 24 | 20 |  |  |  |

NA – Not applicable

### Distribution of visible floral patterns among ultraviolet floral patterns

The distribution of VIS floral patterns with respect to their UV floral patterns in the studied plant was represented as a chord diagram (Figure 5). There was significant variation observed in the distribution of VIS patterns with respect to UV patterns in the studied plants (*χ ^2^*= 107.61; *df*=12; *P* < 0.001). In visible patternless (VIS-P) flowers (N = 87), 42% displayed an absorbing pattern (UV-A), followed by a bullseye pattern (33%; UV-BE) in the ultraviolet spectra. In visible bullseye (VIS-BE) patterned flowers (N = 18), 72% displayed a bullseye pattern (UV-BE), followed by an absorbing pattern (22%; UV-A) in the ultraviolet spectra. In visible contrasting reproductive structure (VIS-CR) patterned flowers (N = 45), 53% displayed contrasting reproductive structures (UV-CR), followed by an absorbing (36%; UV-A) pattern in the ultraviolet spectra. In VIS contrasting corolla (VIS-CC) patterned flowers (N = 38), 44% displayed a contrasting corolla pattern (UV-CC), followed by an absorbing (UV-A) in the ultraviolet spectrum (Figure 5).

**Figure 5:**
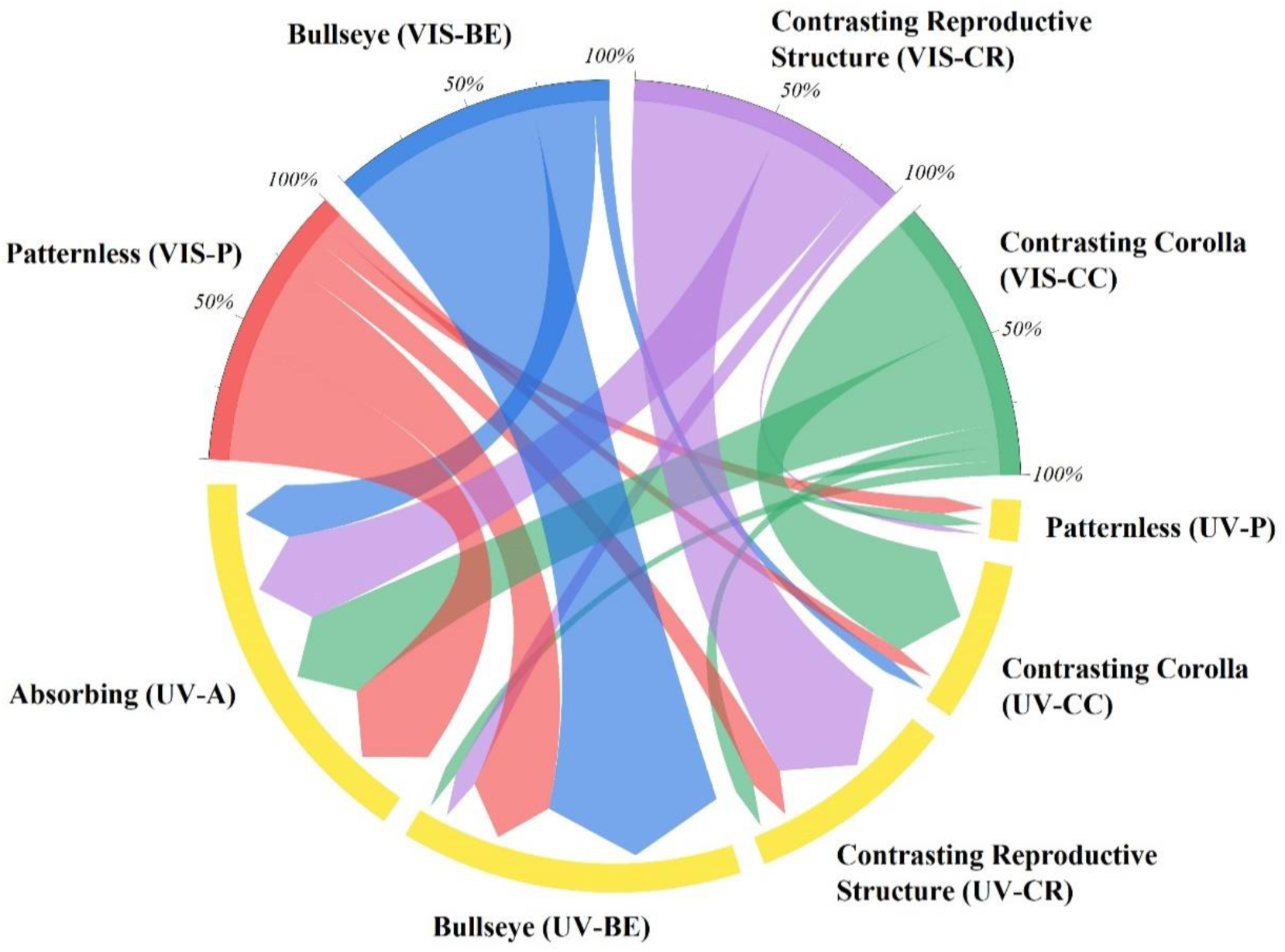
Chord diagram showing the distribution of visible (VIS) floral patterns with respect to ultraviolet (UV) floral patterns in studied plants (N = 188). The chords are unidirectional. Chord thickness corresponds to the percentage of VIS floral pattern related to a given UV floral pattern (χ ^2^= 107.61; df=12; < 0.001).

### Pollinator relationship among visible floral patterns

There was no significant variation observed in pollinators among the different visible floral pattern categories in the studied plants (*χ^2^*= 34.73, *df* = 27, *P* = 0.14). Analysis of the relationship between visible floral patterns and pollinators reveals that VIS patternless flowers (VIS-P) were found to be pollinated by melittophily (37%), followed by melittophily & psychophily (25%), and generalist pollinator (20%) categories. Visible bullseye (VIS-BE) pattern flowers were observed to be pollinated by melittophily (28%), melittophily & psychophily (28%), and generalist pollinator (28%) categories. Visible contrasting corolla (VIS-CC) patterned flowers were pollinated by generalists (32%), followed by melittophily & psychophily (24%). Flowers exhibiting a visible contrasting reproductive structure (VIS-CR) pattern were pollinated by melittophily (42%), followed by melittophily & psychophily (31%) categories (Figure 6). Almost all pollinators were found to be recorded on plants with visible patternless (VIS-P) flowers (Table 2).

**Figure 6:**
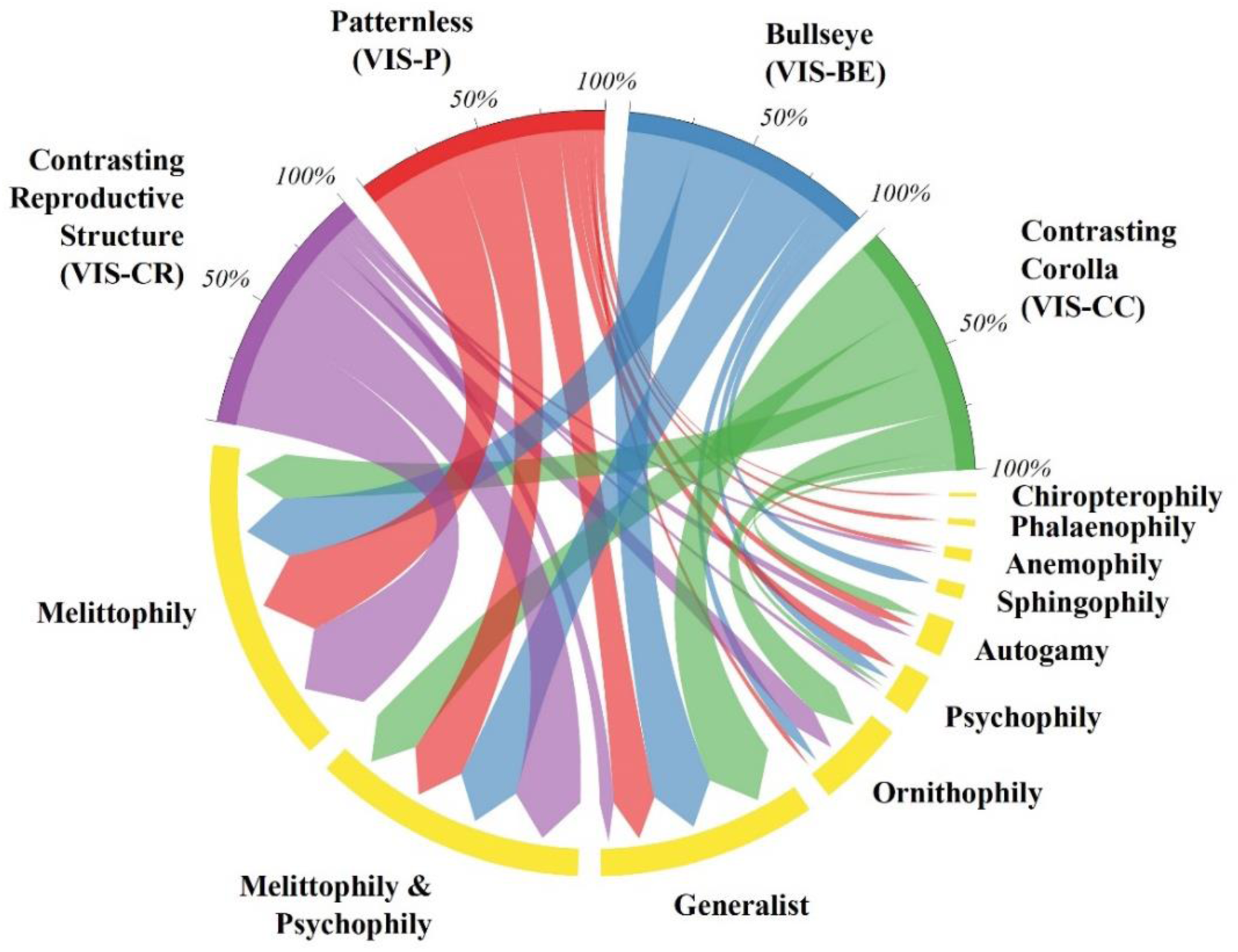
Chord diagram showing the relationship of pollinators with respect to visible (VIS) floral patterns in studied plants (N = 188). The chords are unidirectional. Chord thickness corresponds to the percentage of VIS floral pattern related to a given pollinator (*χ2* = 34.73, *df* = 27, *P* = 0.14).

**Table 2:** Number of plants with visible (VIS) floral patterns with respect to pollinators in the studied plant species.

| Pollinators | Visible Pattern Categories (N = 188) | | | | $\chi^2$ | df | P |
| --- | --- | --- | --- | --- | --- | --- | --- |
|  | VIS-P | VIS-BE | VIS-CC | VIS-CR |  |  |  |
| Anemophily | 2 | - | - | 1 | 1.29 | 3 | 0.73 |
| Autogamy | 4 | - | 2 | 2 | 0.92 | 3 | 0.81 |
| Generalist | 17 | 5 | 12 | 3 | 8.97 | 3 | 0.03 |
| Chiropterophily | 1 | - | - | - | 1.16 | 3 | 0.76 |
| Melittophily | 32 | 5 | 8 | 19 | 4.80 | 3 | 0.18 |
| Ornithophily | 2 | 1 | 6 | 5 | 8.15 | 3 | 0.04 |
| Psychophily | 5 | 1 | 1 | 1 | 1.25 | 3 | 0.74 |
| Phalaenophily | 2 | - | - | - | 2.34 | 3 | 0.50 |
| Sphingophily | - | 1 | - | - | 9.49 | 3 | 0.02 |
| Melittophily &<br>Psychophily | 22 | 5 | 9 | 14 | 0.72 | 3 | 0.86 |

**Visible Pattern Categories** - Patternless (**VIS-P**), Bullseye (**VIS-BE**), Contrasting Corolla (**VIS-CC**), Contrasting Reproductive Structures (**VIS-CR**).

### Pollinator relationship among visible floral patterns with respect to invasive status

A significant difference was observed in the pollinator relationship among the visible floral patterns in non-invasive plants (*χ2* = 35.13, *df* = 18, *P* = 0.009), whereas no significant variation was evidenced in the pollinator relationship among the visible floral patterns in invasive plants (*χ2* = 19.691, *df* = 24, *P* = 0.714). Analysis of the relationship between visible floral patterns and pollinators among invasive plant species revealed that plants with VIS patternless (VIS-P) and VIS bullseye (VIS-BE) patterned flowers were mostly pollinated by melittophily & psychophily (34% and 50%, respectively), whereas invasive plants with VIS contrasting corolla (VIS-CC) and VIS contrasting reproductive structure (VIS-CR) patterned flowers were mostly pollinated by melittophily (36% and 40%, respectively). Analysis of the relationship between visible floral patterns and pollinators among non-invasive plant species revealed that plants with VIS patternless (VIS-P) and VIS contrasting reproductive structure (VIS-CR) patterned flowers were mostly pollinated by melittophily (45% and 44%, respectively), whereas non-invasive plants with VIS bullseye (VIS-BE) and VIS contrasting corolla (VIS-CC) patterned flowers were mostly pollinated by generalist pollinators (Table 3).

**Table 3:** Variations in pollinators among visible pattern categories with respect to invasive status in the studied plant species (*χ^2^* = 34.73;

| Invasive Status | Visible Pattern Categories | Pollinators (%) | | | | | | | | | | $\chi^2$ | $df$ | $P$ |
| --- | --- | --- | --- | --- | --- | --- | --- | --- | --- | --- | --- | --- | --- | --- |
|  |  | Anemophily | Autogamy | Chiropterophily | Generalist | Melittophily | Melittophily & Psychophily | Ornithophily | Phalaenophily | Psychophily | Sphingophily |  |  |  |
| Invasive | *VIS-P* | 5 | 11 | 3 | 5 | 26 | 34 | 3 | 5 | 8 | - | 19.69 | 24 | 0.714 |
|  | *VIS-BE* | - | - | - | 12 | 38 | 50 | - | - | - | - |  |  |  |
|  | *VIS-CC* | - | - | - | 14 | 36 | 29 | 21 | - | - | - |  |  |  |
|  | *VIS-CR* | 5 | 10 | - | 5 | 40 | 35 | 5 | - | - | - |  |  |  |
| Non-invasive | *VIS-P* | - | - | - | 31 | 45 | 18 | 2 | - | 4 | - | 35.13 | 18 | 0.009 |
|  | *VIS-BE* | - | - | - | 40 | 20 | 10 | 10 | - | 10 | 10 |  |  |  |
|  | *VIS-CC* | - | 8 | - | 42 | 13 | 21 | 12 | - | 4 | - |  |  |  |
|  | *VIS-CR* | - | - | - | 8 | 44 | 28 | 16 | - | 4 | - |  |  |  |

**Visible Pattern Categories** - Patternless (*VIS-P*), Bullseye (*VIS-BE*), Contrasting Corolla (*VIS-CC*), Contrasting Reproductive Structures (*VIS-CR*).

### Pollinator relationship among ultraviolet floral patterns

There was significant variation in pollinators among the different ultraviolet floral patterns in the studied plants (*χ^2^* = 61.77, *df* = 36, *P* < 0.004). The ultraviolet absorbing (UV-A) patterned flowers were pollinated by melittophily & psychophily (33%), generalist pollinators (24%), and melittophily (21%), according to the analysis of the relationship between ultraviolet floral patterns and pollinators. Plants that had ultraviolet bullseye patterned (UV-BE) flowers were pollinated by melittophily (38%), followed by melittophily & psychophily (27%), and generalists (19%) categories. Ultraviolet contrasting corolla (UV-CC) patterned flowers were pollinated by melittophily (35%), followed by generalists (26%). Plants that had ultraviolet contrasting reproductive structure (UV-CR) patterned flowers were mostly pollinated by melittophily (55%), followed by melittophily & psychophily (21%) categories. Ultraviolet patternless (UV-P) flowers were pollinated by melittophily (22%) and ornithophily (22%) categories (Figure 7). Almost all pollinators were found to be recorded in plants with ultraviolet absorbing (UV-A) and ultraviolet bullseye (UV-BE) patterned flowers (Table 4).

**Figure 7:**
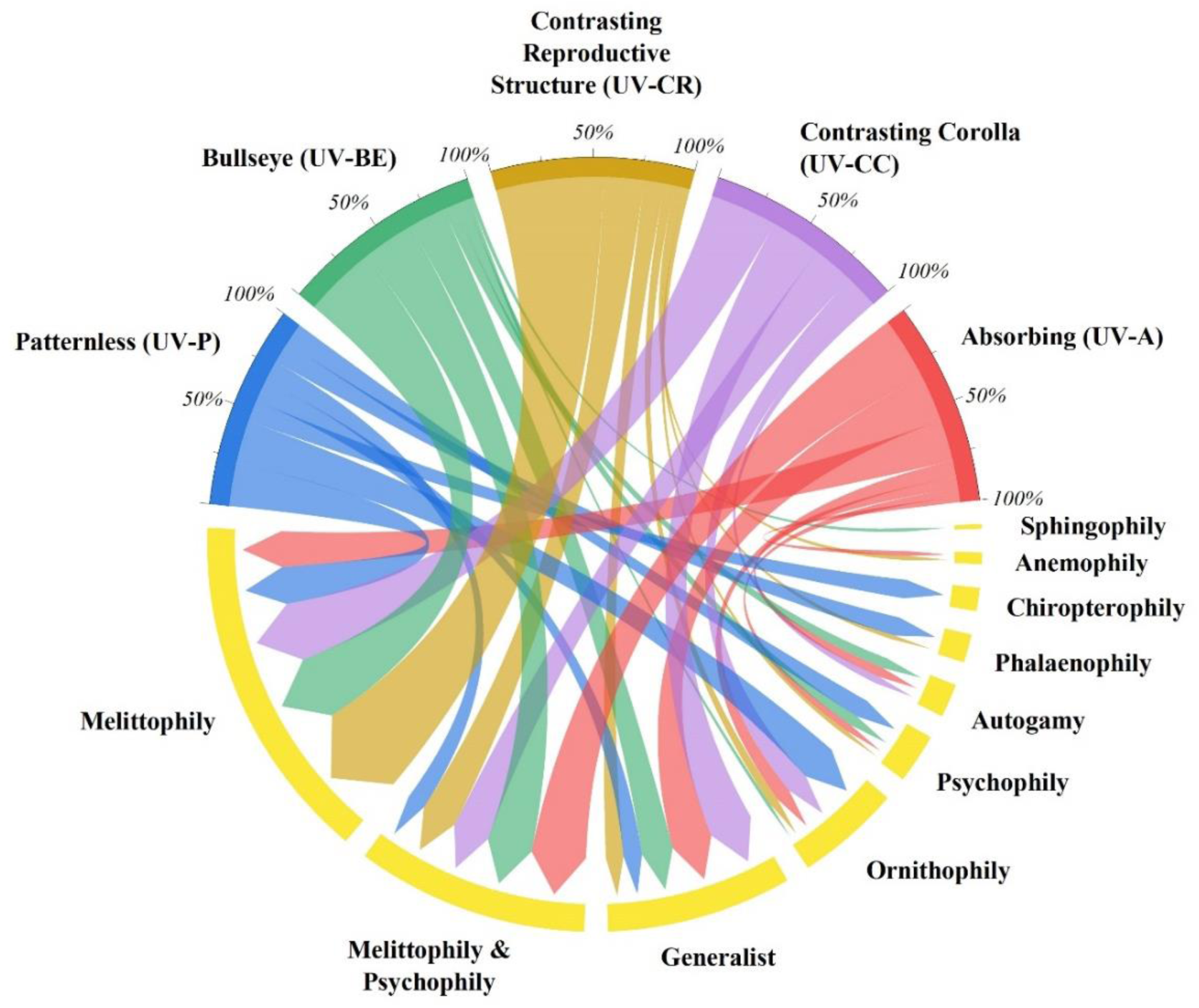
Chord diagram showing the relationship of pollinators with respect to ultraviolet (UV) floral patterns in studied plants (N = 188). The chords are unidirectional. Chord thickness corresponds to the percentage of UV floral pattern related to a given pollinator (*χ2* = 61.77, *df* = 36, *P* = 0.04).

**Table 4:** Distribution of number of plants with ultraviolet (UV) floral patterns with respect to pollinators in the studied plant species.

| Pollinators | Ultraviolet Pattern Categories<br>(N = 188) | | | | | $\chi^2$ | df | P |
| --- | --- | --- | --- | --- | --- | --- | --- | --- |
|  | UV-A | UV-P | UV-BE | UV-CC | UV-CR |  |  |  |
| Anemophily | 2 | - | - | - | 1 | 2.26 | 4 | 0.68 |
| Autogamy | 4 | - | 3 | 1 | - | 2.92 | 4 | 0.57 |
| Generalist | 17 | 1 | 9 | 6 | 4 | 3.99 | 4 | 0.40 |
| Chiropterophily | - | 1 | - | - | - | 19.99 | 4 | < 0.01 |
| Melittophily | 15 | 2 | 18 | 8 | 21 | 13.40 | 4 | < 0.01 |
| Ornithophily | 6 | 2 | 1 | 3 | 2 | 6.29 | 4 | 0.17 |
| Psychophily | 3 | 1 | 3 | - | 1 | 2.77 | 4 | 0.59 |
| Phalaenophily | - | 1 | - | - | 1 | 11.03 | 4 | 0.02 |
| Sphingophily | - | - | 1 | - | - | 2.93 | 4 | 0.56 |
| Melittophily &<br>Psychophily | 23 | 1 | 13 | 5 | 8 | 3.39 | 4 | 0.49 |

**Ultraviolet Pattern Categories** - Ultraviolet Absorbing (**UV-A**), Ultraviolet Patternless (**UV-P**), Ultraviolet Bullseye (**UV-BE**), Ultraviolet Contrasting Corolla (**UV-CC**), Ultraviolet Contrasting Reproductive Structures (**UV-CR**).

### Pollinator relationship among ultraviolet floral patterns with respect to invasive status

A significant difference was observed in the pollinator relationship among the ultraviolet floral patterns in invasive plants (*χ2* = 63.98, *df* = 32, *P* < 0.001), whereas no significant variation was evidenced in the pollinator relationship among the ultraviolet floral patterns in non-invasive plants (*χ2* = 19.50, *df* = 24, *P* = 0.724). Analysis of the relationship between ultraviolet floral patterns and pollinators among invasive plant species revealed that plants with UV absorbing (UV-A) and UV bullseye (UV-BE) patterned flowers were mostly pollinated by melittophily & psychophily (46% and 42%, respectively), whereas invasive plants with UV contrasting reproductive structure (UV-CR) patterned flowers were predominantly (65%) pollinated by melittophily. Invasive plants with UV contrasting corolla (UV-CC) patterned flowers were equally pollinated by melittophily, melittophily & psychophily, and ornithophily (29% each). Analysis of the relationship between ultraviolet floral patterns and pollinators among non-invasive plant species revealed that plants with UV bullseye (UV-BE), UV contrasting corolla (UV-CC), and UV contrasting reproductive structure (UV-CR) patterned flowers were predominantly pollinated by melittophily (40%, 38%, and 48%, respectively), whereas non-invasive plants with UV absorbing (UV-A) patterned flowers were mostly pollinated by generalists (34%) (Table 5).

**Table 5:** Variations in pollinators among ultraviolet pattern categories with respect to invasive status in the studied plant species (*χ^2^* = 61.77; *df* = 36; *P* = 0.004).

| Invasive Status | Ultraviolet Pattern Categories | Pollinators (%) | | | | | | | | | | $\chi^2$ | $df$ | $P$ |
| --- | --- | --- | --- | --- | --- | --- | --- | --- | --- | --- | --- | --- | --- | --- |
|  |  | Anemophily | Autogamy | Chiropterophily | Generalist | Melittophily | Melittophily & Psychophily | Ornithophily | Phalaenophily | Psychophily | Sphingophily |  |  |  |
| Invasive | *UV-P* | - | - | 25 | 25 | - | - | 25 | 25 | - | - | 63.98 | 32 | < 0.001 |
|  | *UV-A* | 8 | 11 | - | 8 | 15 | 46 | 8 | - | 4 | - |  |  |  |
|  | *UV-BE* | - | 12 | - | 4 | 35 | 42 | - | - | 7 | - |  |  |  |
|  | *UV-CC* | - | - | - | 13 | 29 | 29 | 29 | - | - | - |  |  |  |
|  | *UV-CR* | 6 | - | - | 6 | 65 | 17 | - | 6 | - | - |  |  |  |
| Non-invasive | *UV-P* | - | - | - | - | 40 | 20 | 20 | - | 20 | - | 19.50 | 24 | 0.724 |
|  | *UV-A* | - | 2 | - | 34 | 25 | 25 | 9 | - | 5 | - |  |  |  |
|  | *UV-BE* | - | - | - | 36 | 40 | 9 | 5 | - | 5 | 5 |  |  |  |
|  | *UV-CC* | - | 6 | - | 31 | 38 | 19 | 6 | - | - | - |  |  |  |
|  | *UV-CR* | - | - | - | 14 | 48 | 24 | 10 | - | 4 | - |  |  |  |

**Ultraviolet Pattern Categories** - Ultraviolet Absorbing (*UV-A*), Ultraviolet Patternless (*UV-P*), Ultraviolet Bullseye (*UV-BE*), Ultraviolet Contrasting Corolla (*UV-CC*), Ultraviolet Contrasting Reproductive Structures (*UV-CR*).

### Distribution of visible floral patterns among flower color groups

There was no significant variation in visible floral pattern categories among the flower color groups in the studied plants (*χ^2^* = 11.43, *df* = 6, *P* = 0.07). The results show that overall, 38% of the plant species had flower color in the YOR spectrum, followed by White (37%) and VIB (25%) color spectrum categories (Figure 8). The distribution of visible (VIS) flower patterns among flower color groups showed that the visible patternless (VIS-P) category was found to be the highest (57%) in plants belonging to the YOR flower color group, whereas the visible bullseye (VIS-BE) pattern category was the least (2%) among the plants belonging to the YOR flower color group. In plants belonging to the White color group, the visible patternless (VIS-P) category was the highest (43%) and the visible bullseye (VIS-BE) pattern category was the least (15%). In plants belonging to the VIB flower color group, the visible patternless (VIS-P) category was found to be the highest (34%), and the visible bullseye (VIS-BE) pattern category was the least (13%) (Table 6).

**Figure 8:**
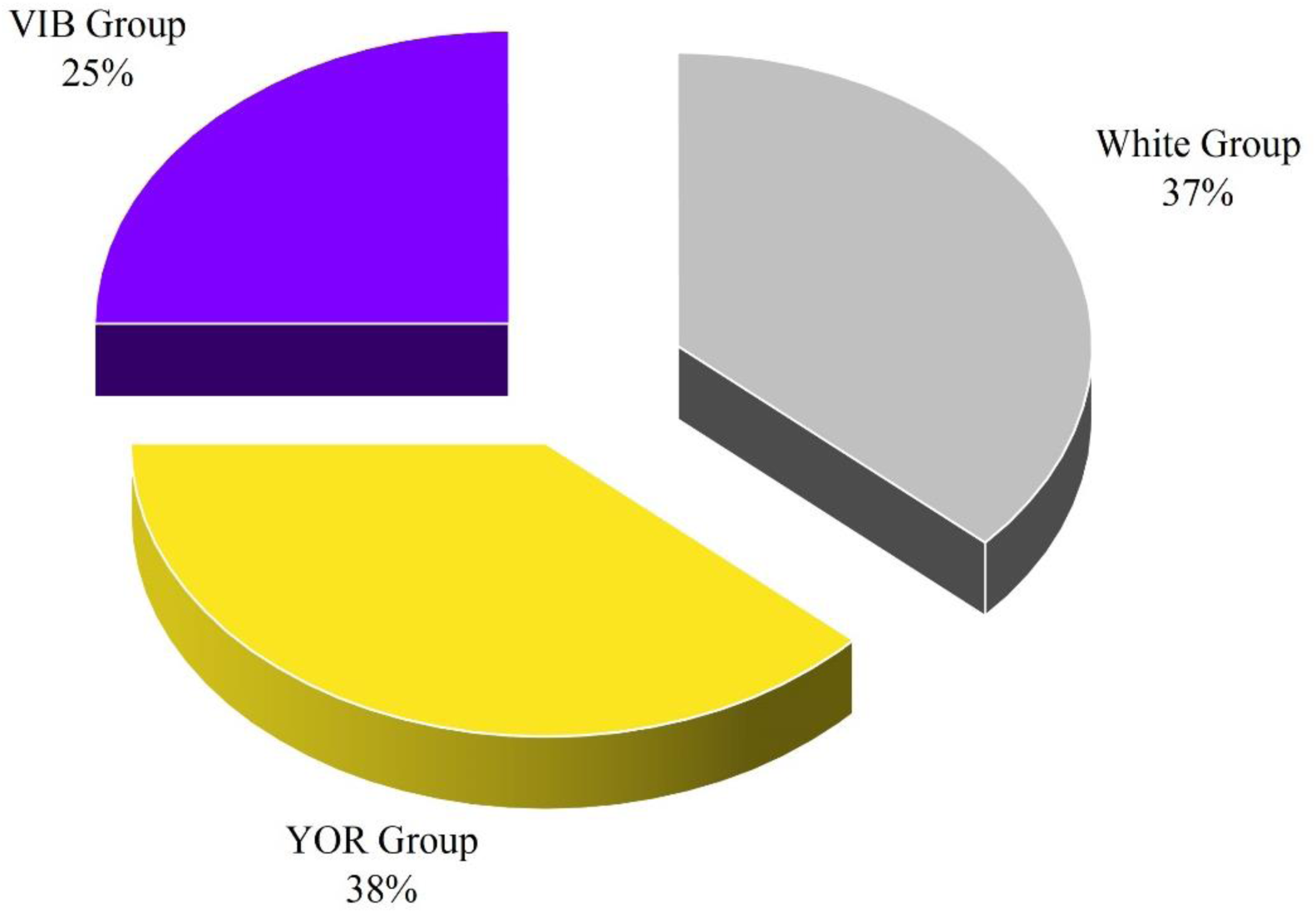
Distribution of the studied plant species (N = 188) with respect to flower color group classification. VIB Group – Flower colors belonging to violet, indigo, blue color range of the visible spectrum, White Group – Flower colors in the range of white, pale white, yellowish-white, pinkish white, and YOR Group – Flower colors belonging to yellow, orange, red range of the visible spectrum.

**Table 6:** Distribution of visible (VIS) floral pattern categories with respect to flower color groups in the studied plant species.

| **Flower Color Group** | **Visible Pattern Categories (%) [N = 188]** |  |  |  |
| --- | --- | --- | --- | --- |
|  | Patternless (VIS-P) | Bullseye (VIS-BE) | Contrasting Corolla (VIS-CC) | Contrasting Reproductive Structures (VIS-CR) |
| *White*<sup>#</sup> | 43 | 15 | 16 | 26 |
| *VIB* * | 34 | 13 | 25 | 28 |
| *YOR*<sup>^</sup> | 57 | 2 | 21 | 20 |

\**VIB Group* – Flower colors belonging to violet, indigo, blue color range of the visible spectrum
<sup>#</sup>*White Group* – Flower colors in the range of white, pale white, yellowish-white, pinkish white
<sup>^</sup>*YOR Group* – Flower colors belonging to yellow, orange, red range of the visible spectrum

### Distribution of ultraviolet floral patterns among flower color groups

There was significant variation in the ultraviolet floral pattern categories among the flower color groups in the studied plants (*χ2* = 20.26, *df* = 8, *P* = 0.01). The distribution of ultraviolet (UV) floral patterns among the flower color groups showed that the ultraviolet absorbing (UV-A) pattern category was found to be highest (52%) in plants belonging to White flower color group and the ultraviolet contrasting corolla (UV-CC) pattern category was least (7%) in plants belonging to the White flower color group. In plants belonging to the VIB flower color group, the ultraviolet bullseye (UV-BE) pattern category was found to be the highest (32%), and the ultraviolet patternless (UV-P) category was the least (2%). In plants belonging to the YOR flower color group, the ultraviolet absorbing (UV-A) pattern category was found to be the highest (29%), while the ultraviolet patternless (UV-P) category was found to be the least (3%) (Table 7).

**Table 7:** Distribution of ultraviolet (UV) floral pattern categories with respect to flower color groups in the studied plant species.

| **Flower Color Group** | **Ultraviolet Pattern Categories (in %) [N = 188]** |  |  |  |  |
| --- | --- | --- | --- | --- | --- |
|  | Absorbing (UV-A) | Patternless (UV-P) | Bullseye (UV-BE) | Contrasting Corolla (UV-CC) | Contrasting Reproductive Structures (UV-CR) |
| *White*<sup>#</sup> | 52 | 9 | 22 | 7 | 10 |
| *VIB* * | 28 | 2 | 32 | 11 | 27 |
| *YOR*<sup>^</sup> | 29 | 3 | 25 | 18 | 25 |

\**VIB Group* – Flower colors belonging to violet, indigo, blue color range of the visible spectrum
<sup>#</sup>*White Group* – Flower colors in the range of white, pale white, yellowish-white, pinkish white
<sup>^</sup>*YOR Group* – Flower colors belonging to yellow, orange, red range of the visible spectrum

### Pollinator relationship among flower color groups

There was significant variation in pollinators among the flower color groups in the studied plants (*χ^2^* = 31.77, *df* = 18, *P* = 0.02). The plants belonging to White color group were pollinated by melittophily & psychophily (30%), followed by generalists (29%), and melittophily (26%) categories. Plants belonging to VIB flower color groups were mostly pollinated by melittophily (40%), followed by melittophily & psychophily (19%) categories. YOR flower color group plants were found to be pollinated by melittophily (38%), followed by melittophily & psychophily (28%) categories (Figure 9).

**Figure 9:**
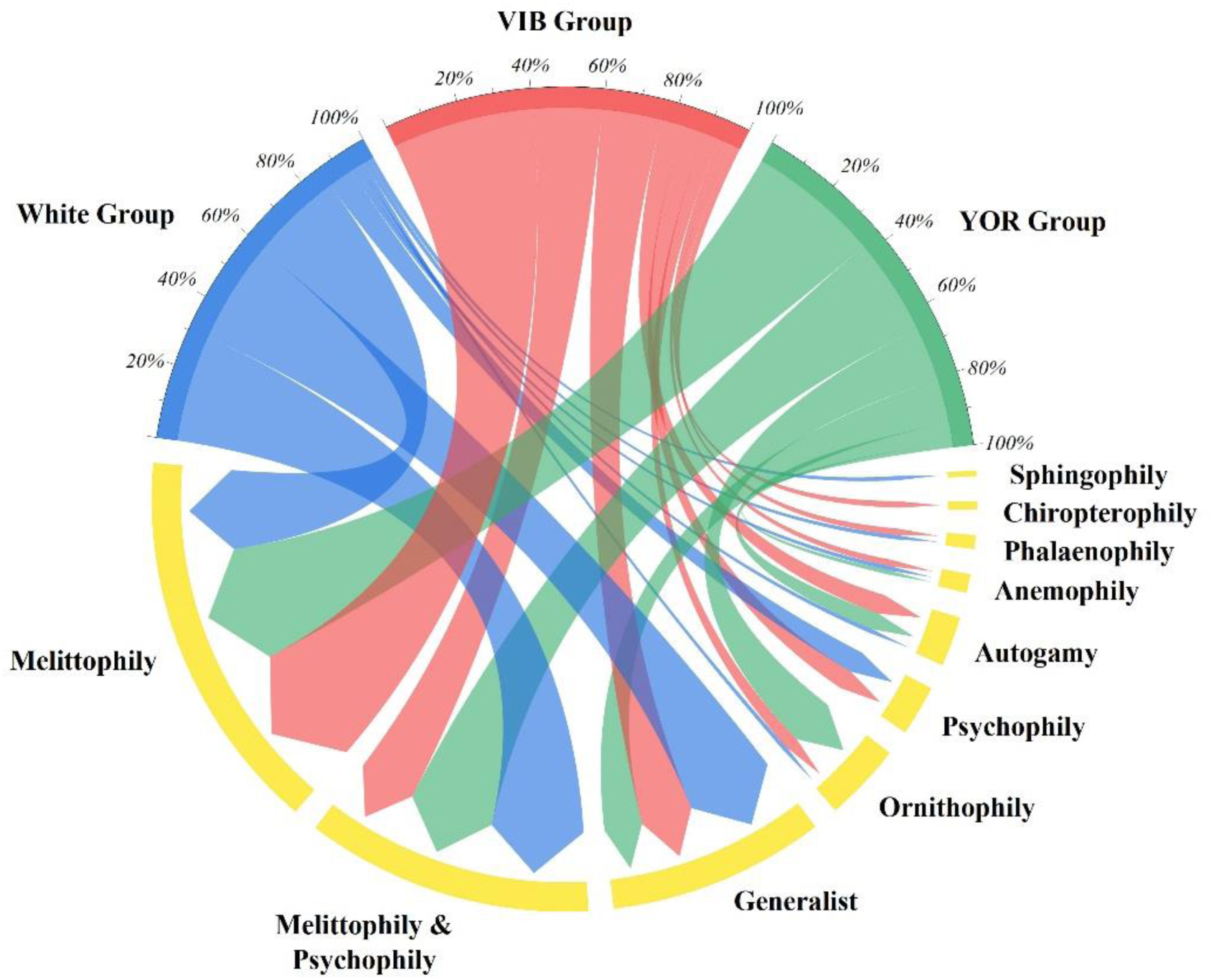
Chord diagram showing the relationship of pollinators with respect to flower color groups in studied plants (N = 188). The chords are unidirectional. Chord thickness corresponds to the percentage of flower color group to a given pollinator association (*χ2* = 31.77, *df* = 18, *P* = 0.02).

### Flower color group distribution with respect to invasive status

There was significant variation in the distribution of flower color groups with respect to the invasive status of the studied plants (*χ2* = 6.618, *df* = 2, *P* = 0.03). The distribution of flower color groups among invasive plant species revealed that YOR and VIB flower color groups were predominant (44% and 30%, respectively) in invasive plant species. Similarly, in the case of non-invasive species, observations found that the White color group was predominant (44%), followed by the YOR color group (34%) (Figure 10).

**Figure 10:**
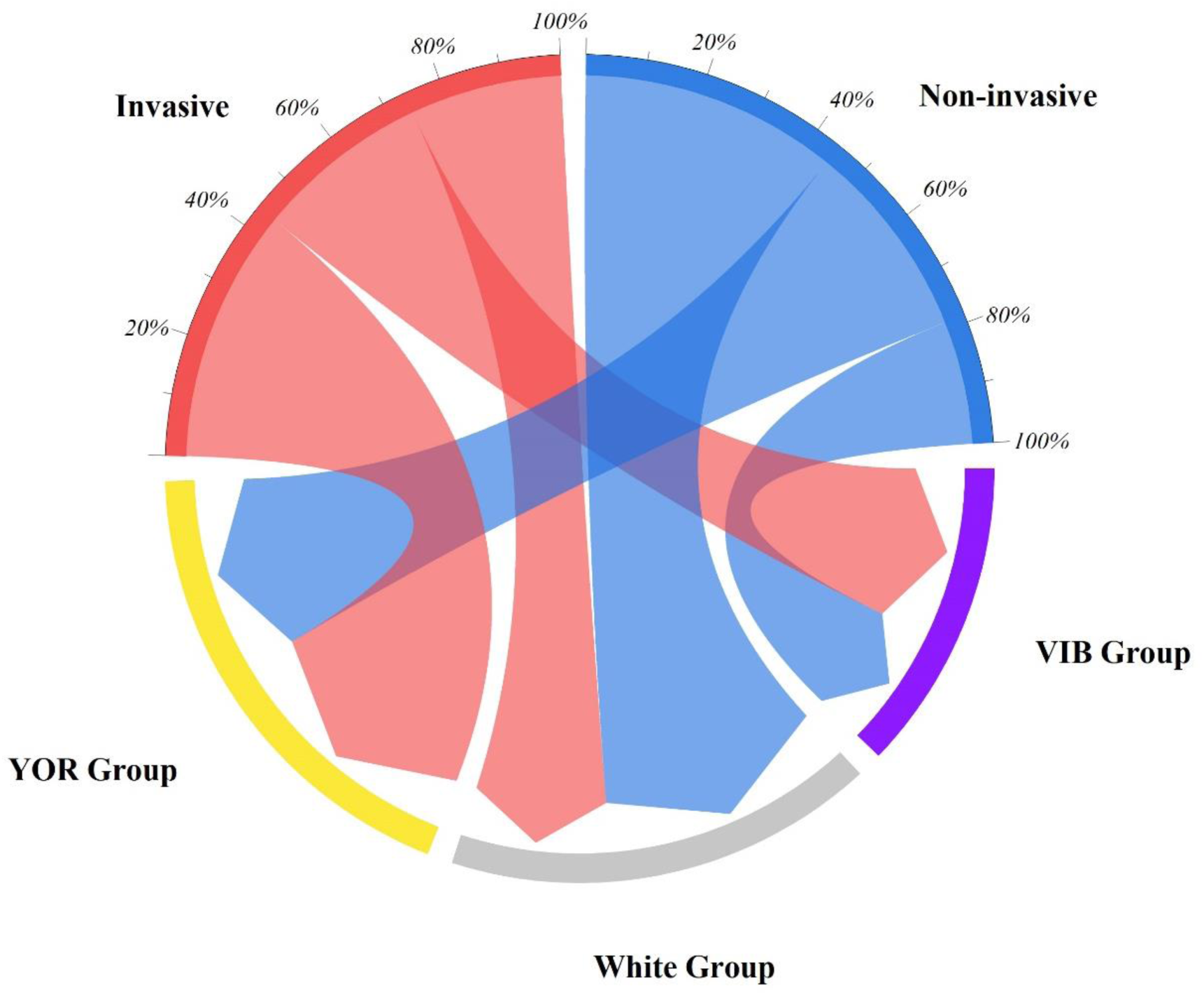
Chord diagram showing the distribution of flower color groups with respect to invasive status in studied plants (N = 188). The chords are unidirectional. Chord thickness corresponds to the percentage of invasive status related to a given flower color group (*χ2* = 6.618, *df* = 2, *P* = 0.03). VIB Group – Flower colors belonging to violet, indigo, blue color range of the visible spectrum, White Group – Flower colors in the range of white, pale white, yellowish-white, pinkish white, and YOR Group – Flower colors belonging to yellow, orange, red range of the visible spectrum.

### Pollinator relationship with respect to invasive status

There was significant variation in pollinators with respect to the invasive status of the studied plants (*χ^2^* = 26.931, *df* = 9, *P* = 0.001). In the studied plants, invasive species were pollinated by more types of pollinator categories than non-invasive species. Invasive plants were found to be pollinated by melittophily & psychophily (35%), followed by melittophily (33%) categories. Non-invasive plants, on the other hand, were found to be pollinated by melittophily (35%), followed by generalist (29%) categories (Figure 11).

**Figure 11:**
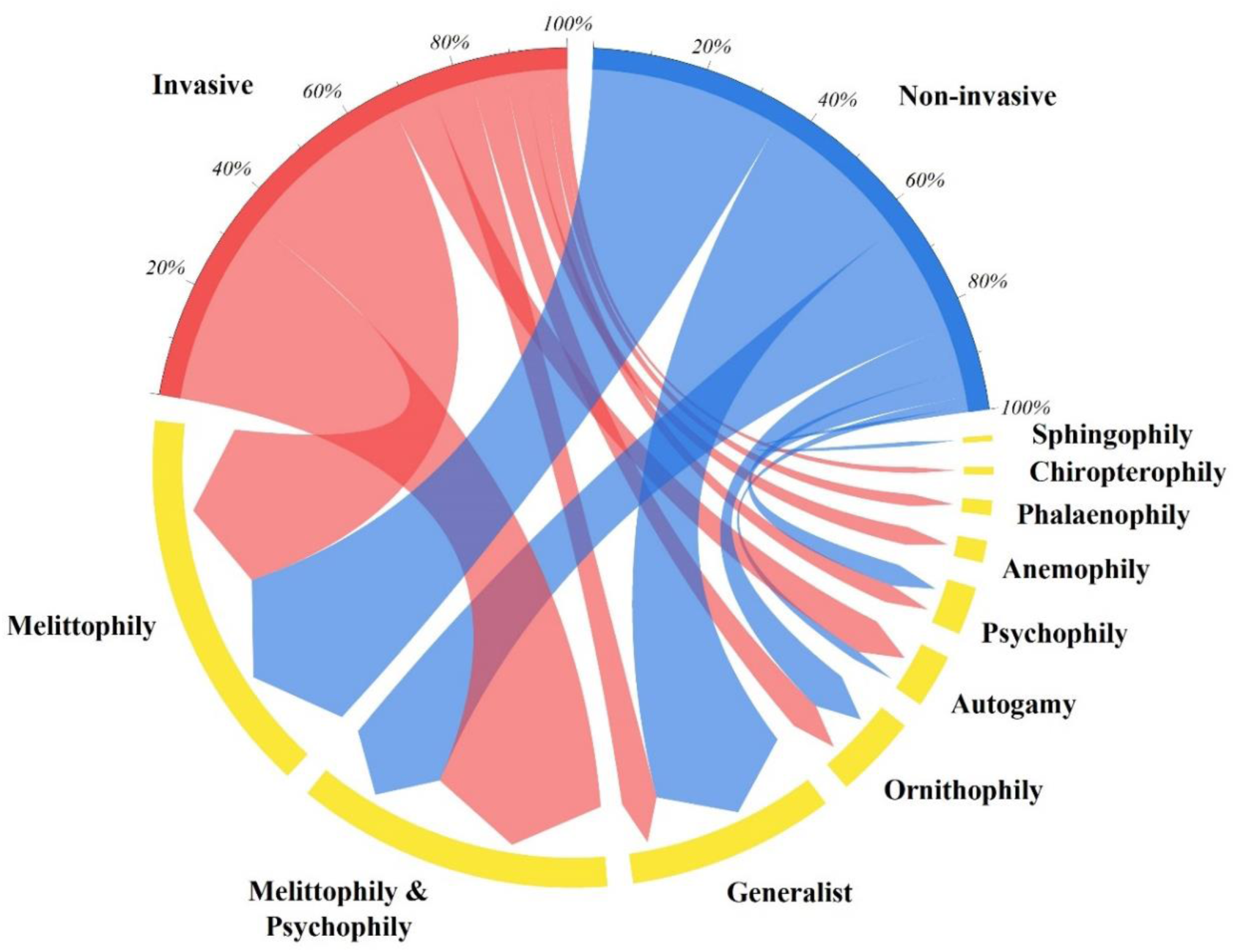
Chord diagram showing the relationship of pollinators with respect to invasive status in studied plants (N = 188). The chords are unidirectional. Chord thickness corresponds to the percentage of invasive status related to a given pollinator (*χ2* = 26.931, *df* = 9, *P* = 0.001).

### Visible floral pattern distribution with respect to invasive status

There was no significant variation in the distribution of visible floral patterns with respect to the invasive status of the studied plants (*χ^2^*= 0.644, *df* = 3, *P* = 0.88). The invasive and non-invasive plants had similar distributions toward each category of visible floral patterns. About 48% of invasive plants were found to have visible patternless (VIS-P) flowers, followed by visible contrasting reproductive structure (VIS-CR; 25%) floral pattern, visible contrasting corolla (VIS-CC; 17%) floral pattern, and visible bullseye (VIS-BE; 10%) floral pattern. Similarly, among non-invasive species, about 45% were found to have visible patternless (VIS-P) flowers, followed by visible contrasting reproductive structures (VIS-CR; 23%) floral pattern, visible contrasting corolla (VIS-CC; 22%) floral pattern, and visible bullseye (VIS-BE; 10%) floral pattern (Figure 12).

**Figure 12:**
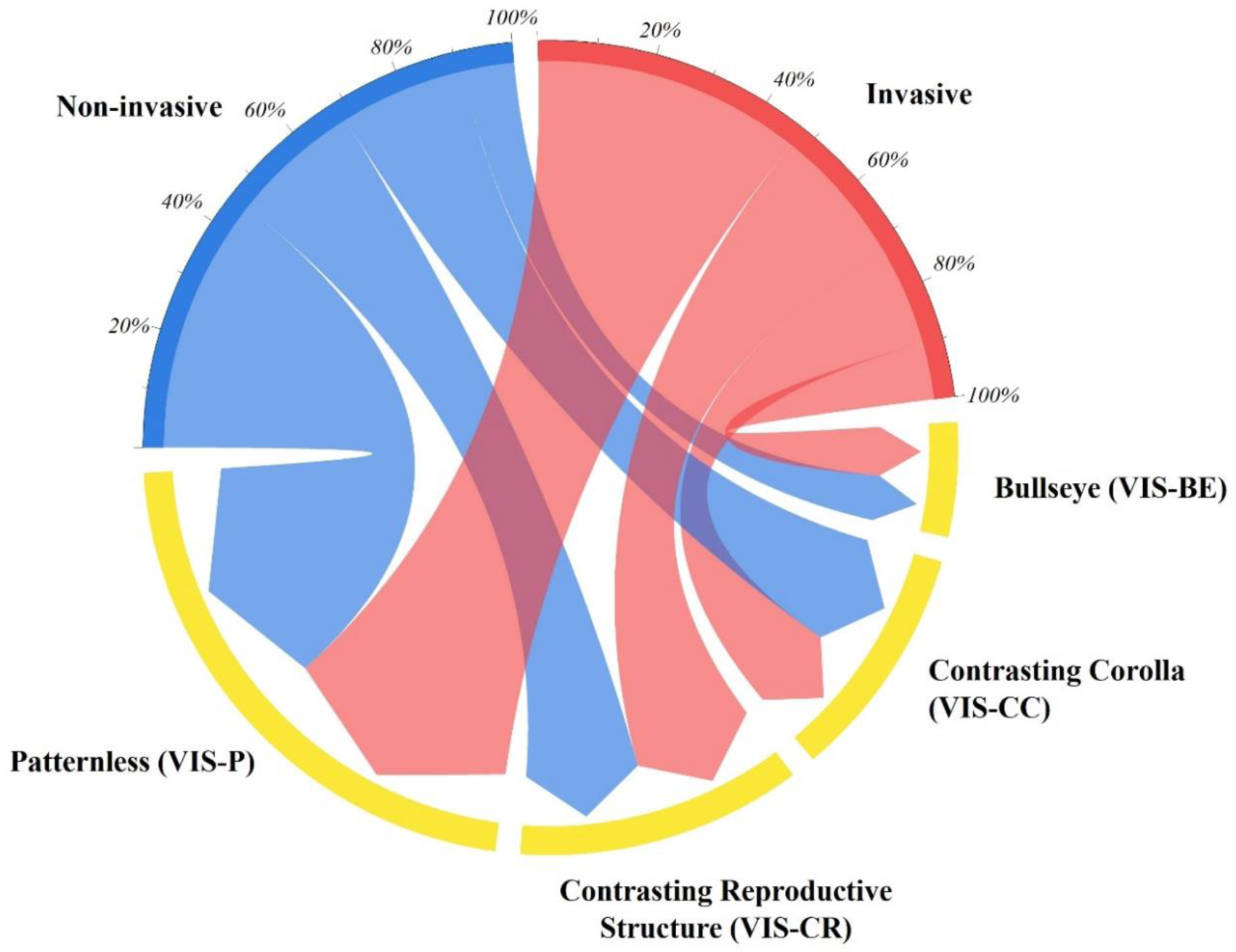
Chord diagram showing the distribution of visible (VIS) floral patterns with respect to invasive status in studied plants (N = 188). The chords are unidirectional. Chord thickness corresponds to the percentage of invasive status related to a given visible floral pattern category (*χ2* = 0.644, *df* = 3, *P* = 0.88).

### Ultraviolet floral pattern distribution with respect to invasive status

There was no significant variation in the distribution of ultraviolet floral patterns with respect to the invasive status of the studied plants (*χ^2^* = 4.955, *df* = 4, *P* = 0.29). The invasive and non-invasive plants had almost similar distributions toward each ultraviolet floral pattern category. About 33% of invasive plants were found to have ultraviolet absorbing (UV-A) patterned flowers, followed by ultraviolet bullseye (UV-BE; 32%) floral pattern, ultraviolet contrasting reproductive structures (UV-CR; 21%) floral pattern, ultraviolet contrasting corolla (UV-CC; 9%) floral pattern, and ultraviolet patternless (UV-P; 5%) flowers. Similarly, in non-invasive plants, about 41% of invasive plants were found to have ultraviolet absorbing (UV-A) patterned flowers, followed by ultraviolet bullseye (UV-BE; 20%) floral pattern, ultraviolet contrasting reproductive structures (UV-CR; 19%) floral pattern, ultraviolet contrasting corolla (UV-CC; 15%) floral pattern, and ultraviolet patternless (UV-P; 5%) flowers (Figure 13).

**Figure 13:**
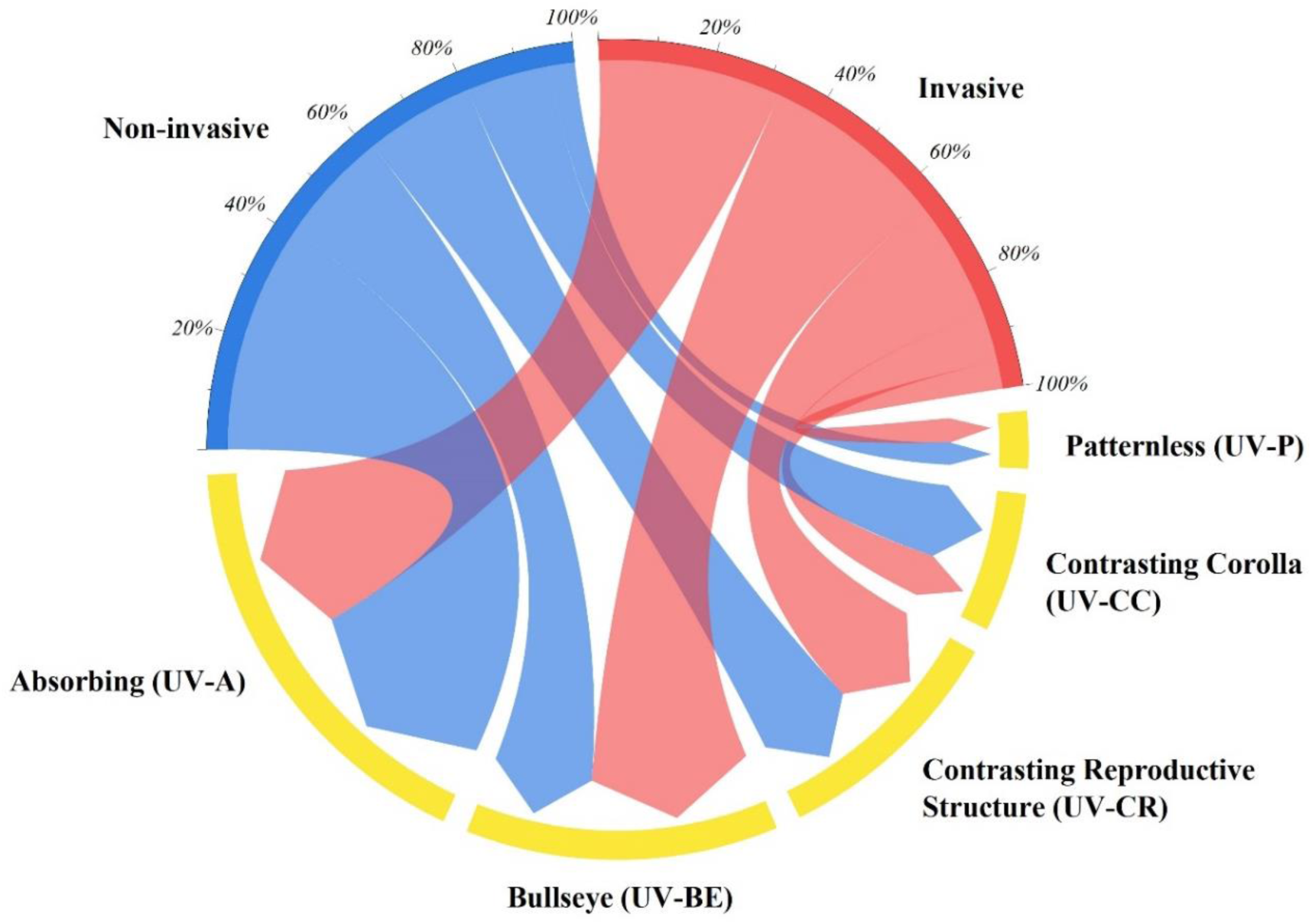
Chord diagram showing the distribution of ultraviolet (UV) floral patterns with respect to invasive status in studied plants (N = 188). The chords are unidirectional. Chord thickness corresponds to the percentage of invasive status related to a given ultraviolet floral pattern category (*χ2* = 4.955, *df* = 4, *P* = 0.29).

### Analysis of invasive status and ultraviolet patterns with respect to plant phylogeny

The ultraviolet floral patterns and invasive status were analyzed in relation to the phylogeny among the studied species (Figure 14). A significant degree of phylogenetic signal was observed between ultraviolet floral patterns and plant species phylogeny (δ = 2.758, *P* < 0.001). The Leguminosae (N = 28) family was the most prevalent in the studied species of plants, followed by the Acanthaceae (N = 16) and Lamiaceae (N = 14) plant families. Among the invasive plants (N = 80) in the study, 26 plants were found to display an ultraviolet absorbing (UV-A) pattern, another 26 plants displayed an ultraviolet bullseye (UV-BE) pattern, and 17 plants were found to display an ultraviolet contrasting reproductive structure (UV-CR) pattern. Among the invasive plants that display an ultraviolet absorbing (UV-A) pattern (N = 26), the contributions of the Compositae (N = 7) and Leguminosae (N = 6) families were greater in comparison to the rest of the families. Among the invasive plants that display an ultraviolet bullseye (UV-BE) pattern (N = 26), the contributions of the Apocynaceae (N = 3), Compositae (N = 3), Convolvulaceae (N = 3), and Verbenaceae (N = 3) families were equally high compared to the rest of the families. Among the invasive plants with an ultraviolet contrasting reproductive structure (UV-CR) pattern (N = 17), the contribution of Leguminosae (N = 5) was greater in comparison to the rest of the families (Table 8).

**Figure 14:**
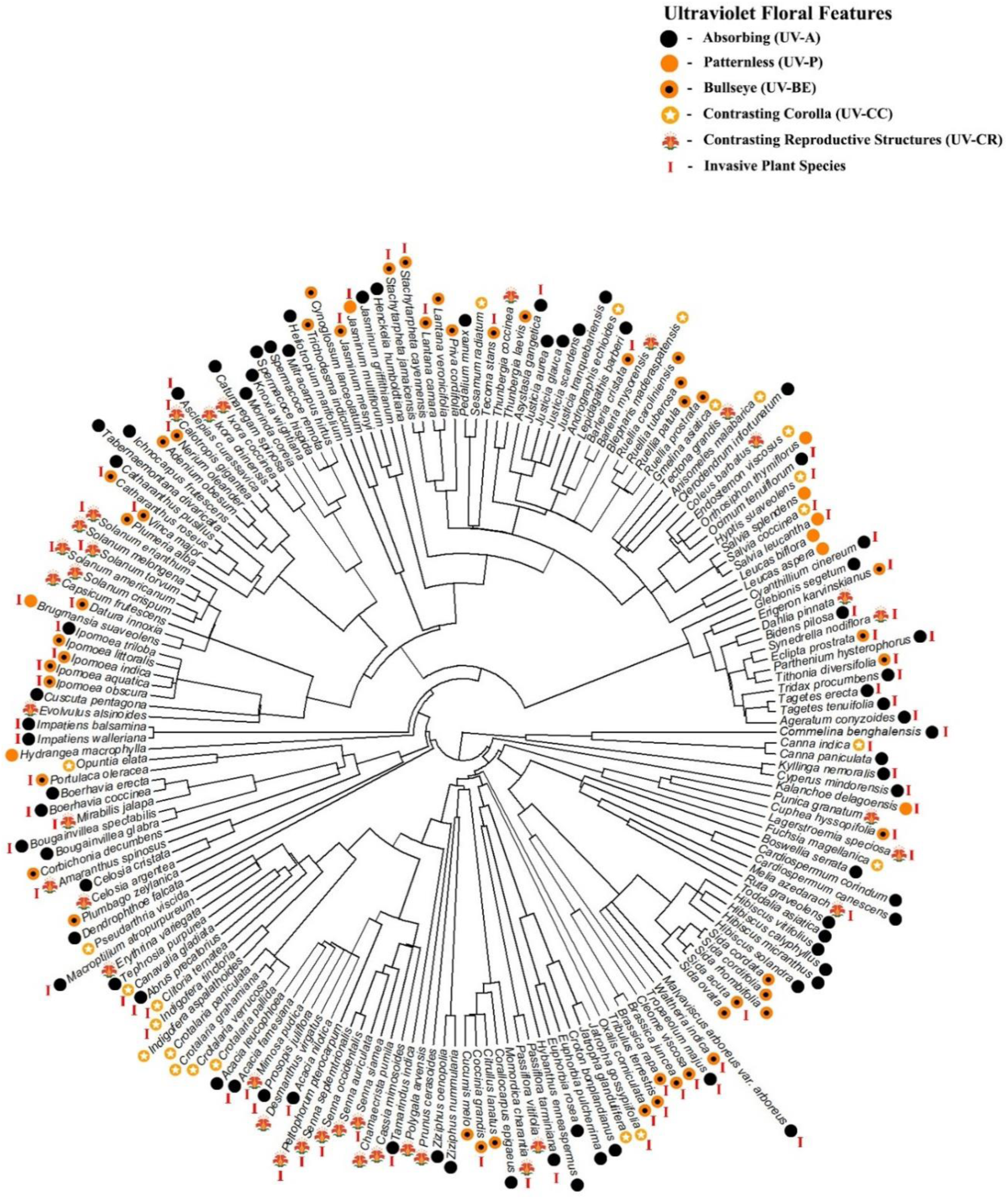
Phylogenetic tree representing studied flowering plant species (N = 188) along with their respective ultraviolet (UV) floral pattern and invasive status. Ultraviolet Absorbing (UV-A), Ultraviolet Patternless (UV-P), Ultraviolet Bullseye (UV-BE), Ultraviolet Contrasting Corolla (UV-CC), Ultraviolet Contrasting Reproductive Structures (UV-CR).

**Table 8:** Distribution of number of plant species among plant families along with their respective visible and ultraviolet floral patterns in the studied plant species (N = 188).

| S. No. | Plant Family | No. of Plant Species |  | Ultraviolet Patterns (N = 188) |  |  |  |  |  |  |  |  |  |
| --- | --- | --- | --- | --- | --- | --- | --- | --- | --- | --- | --- | --- | --- |
|  |  |  |  | Absorbing (UV-A) |  | Patternless (UV-P) |  | Bullseye (UV-BE) |  | Contrasting Corolla (UV-CC) |  | Contrasting Reproductive Structures (UV-CR) |  |
|  |  | Non-invasive | Invasive | Non-invasive | Invasive | Non-invasive | Invasive | Non-invasive | Invasive | Non-invasive | Invasive | Non-invasive | Invasive |
| 1. | Acanthaceae | 14 | 2 | 5 | 1 | - | - | 5 | 1 | 2 | - | 2 | - |
| 2. | Amaranthaceae | 2 | 1 | 1 | - | - | - | - | - | - | - | 1 | 1 |
| 3. | Apocynaceae | 5 | 5 | 3 | 1 | - | - | 2 | 3 | - | - | - | 1 |
| 4. | Balsaminaceae | 0 | 2 | - | 2 | - | - | - | - | - | - | - | - |
| 5. | Bignoniaceae | 0 | 1 | - | - | - | - | - | 1 | - | - | - | - |
| 6. | Boraginaceae | 3 | 0 | 1 | - | - | - | 2 | - | - | - | - | - |
| 7. | Brassicaceae | 0 | 2 | - | - | - | - | - | 2 | - | - | - | - |
| 8. | Burseraceae | 1 | 0 | 1 | - | - | - | - | - | - | - | - | - |
| 9. | Cactaceae | 1 | 0 | - | - | - | - | - | - | 1 | - | - | - |
| 10. | Cannaceae | 1 | 1 | 1 | - | - | - | - | - | - | 1 | - | - |
| 11. | Cleomaceae | 0 | 1 | - | - | - | - | - | 1 | - | - | - | - |
| 12. | Commelinaceae | 0 | 1 | 1 | 1 | - | - | - | - | - | - | - | - |
| 13. | Compositae | 3 | 11 | 1 | 7 | - | - | 1 | 3 | - | - | 1 | 1 |
| 14. | Convolvulaceae | 3 | 4 | 1 | 1 | - | - | 1 | 3 | - | - | 1 | - |
|  |  |  |  | Absorbing (UV-A) |  | Patternless (UV-P) |  | Bullseye (UV-BE) |  | Contrasting Corolla (UV-CC) |  | Contrasting Reproductive Structures (UV-CR) |  |
|  |  | Non-invasive | Non-invasive | Non-invasive | Invasive | Non-invasive | Invasive | Non-invasive | Invasive | Non-invasive | Invasive | Non-invasive | Invasive |
| 15. | Crassulaceae | 0 | 1 | - | - | - | 1 | - | - | - | - | - | - |
| 16. | Cucurbitaceae | 3 | 2 | 1 | - | - | - | 2 | 1 | - | - | - | 1 |
| 17. | Cyperaceae | 0 | 2 | - | 2 | - | - | - | - | - | - | - | - |
| 18. | Euphorbiaceae | 4 | 1 | 3 | - | - | - | - | - | 1 | 1 | - | - |
| 19. | Gesneriaceae | 1 | 0 | 1 | - | - | - | - | - | - | - | - | - |
| 20. | Hydrangeaceae | 1 | 0 | - | - | 1 | - | - | - | - | - | - | - |
| 21. | Lamiaceae | 9 | 5 | 1 | 1 | 4 | 1 | - | - | 3 | 2 | 1 | 1 |
| 22. | Leguminosae | 14 | 14 | 2 | 6 | - | - | - | - | 7 | 3 | 5 | 5 |
| 23. | Lophiocarpaceae | 1 | 0 | - | - | - | - | 1 | - | - | - | - | - |
| 24. | Loranthaceae | 1 | 0 | 1 | - | - | - | - | - | - | - | - | - |
| 25. | Lythraceae | 1 | 2 | - | - | - | - | - | 1 | - | - | 1 | 1 |
| 26. | Malvaceae | 9 | 3 | 4 | 1 | - | - | 5 | 2 | - | - | - | - |
| 27. | Meliaceae | 0 | 1 | - | - | - | - | - | - | - | - | - | 1 |
| 28. | Nyctaginaceae | 2 | 3 | 2 | 2 | - | - | - | - | - | - | - | 1 |
| 29. | Oleaceae | 1 | 2 | 1 | - | - | 1 | - | 1 | - | - | - | - |
| 30. | Onagraceae | 1 | 0 | - | - | - | - | - | - | 1 | - | - | - |
|  |  |  |  | Absorbing (UV-A) |  | Patternless (UV-P) |  | Bullseye (UV-BE) |  | Contrasting Corolla (UV-CC) |  | Contrasting Reproductive Structures (UV-CR) |  |
|  |  | Non-invasive | Non-invasive | Non-invasive | Invasive | Non-invasive | Invasive | Non-invasive | Invasive | Non-invasive | Invasive | Non-invasive | Invasive |
| 31. | Oxalidaceae | 0 | 1 | - | - | - | - | - | 1 | - | - | - | - |
| 32. | Passifloraceae | 1 | 1 | - | 1 | - | - | - | - | - | - | 1 | - |
| 33. | Pedaliaceae | 2 | 0 | 1 | - | - | - | - | - | 1 | - | - | - |
| 34. | Plumbaginaceae | 1 | 0 | - | - | - | - | 1 | - | - | - | - | - |
| 35. | Polygalaceae | 1 | 0 | - | - | - | - | - | - | - | - | 1 | - |
| 36. | Portulacaceae | 0 | 1 | - | - | - | - | - | 1 | - | - | - | - |
| 37. | Rhamnaceae | 2 | 0 | 2 | - | - | - | - | - | - | - | - | - |
| 38. | Rosaceae | 1 | 0 | - | - | - | - | - | - | - | - | 1 | - |
| 39. | Rubiaceae | 8 | 0 | 6 | - | - | - | - | - | - | - | 2 | - |
| 40. | Rutaceae | 2 | 0 | 2 | - | - | - | - | - | - | - | - | - |
| 41. | Sapindaceae | 2 | 0 | 2 | - | - | - | - | - | - | - | - | - |
| 42. | Solanaceae | 4 | 5 | - | - | - | 1 | - | 1 | - | - | 4 | 3 |
| 43. | Tropaeolaceae | 0 | 1 | - | - | - | - | - | - | - | - | - | 1 |
| 44. | Verbenaceae | 2 | 3 | - | - | - | - | 2 | 3 | - | - | - | - |
| 45. | Violaceae | 1 | 0 | 1 | - | - | - | - | - | - | - | - | - |
| 46. | Zygophyllaceae | 0 | 1 | - | - | - | - | - | 1 | - | - | - | - |

Leguminosae was the largest family among the studied species, comprising 28 species, of which 14 were invasive and another 14 were non-invasive. Among the plants in the Leguminosae family, UV contrasting reproductive structures (UV-CR) and UV contrasting corolla (UV-CC) patterns were predominant. The plant species, such as *Mimosa pudica, Peltophorum pterocarpum, Senna septemtrionalis, Senna occidentalis,* and *Senna siamea,* were all found to be invasive and have UV contrasting reproductive structure (UV-CR) patterns. Following this, plant species such as *Canavalia gladiata, Clitoria ternatea,* and *Indigofera tinctoria* were all found to be invasive with a contrasting corolla (UV-CC) pattern. *Abrus precatorius, Acacia farnesiana, Acacia leucophloea,* and *Macroptilium atropurpureum* falling within the family were observed to be invasive and have a UV absorbing (UV-A) pattern. Within the same family, plant species such as *Crotalaria grahamiana, Crotalaria pallida, Crotalaria paniculata, Crotalaria verrucose,* and *Indigofera aspalathoides* were found to be non-invasive while having a UV contrasting corolla pattern (UV-CC).

In the second-most prevalent family in the study, Acanthaceae (N = 16), the most predominant patterns were UV absorbing (UV-A) and UV bullseye (UV-BE). The non-invasive plants belonging to the genus Justicia (*Justicia aurea, Justicia glauca, Justicia scandens,* and *Justicia tranquebariensis*) were found to have an absorbing pattern (UV-A). The non-invasive plants, such as *Ruellia caroliniensis, Ruellia patula, Ruellia prostrata, Ruellia tuberosa,* and *Thunbergia laevis*, were found to have a bullseye (UV-BE) pattern. *Asystasia gangetica* (UV-A) and *Barleria cristata* (UV-BE) were the only two invasive species in the Acanthaceae family.

Plant observations in the Compositae family (N = 14) were mostly invasive species in the present study. Among the family, an absorbing (UV-A) pattern was most prevalent. Plant species such as *Ageratum conyzoides, Cyanthillium cinereum, Tagetes tenuifolia, Tagetes erecta, Tridax procumbens, Parthenium hysterophorus,* and *Bidens pilosa* were reported to be invasive and have UV absorbing pattern (UV-A). Plants such as *Eclipta prostrata, Erigeron karvinskianus,* and *Tithonia diversifolia* in the Compositae family were invasive with the UV bullseye pattern (UV-BE).

The Lamiaceae plant family (N = 14) was mostly UV patternless (UV-P) and UV contrasting corolla (UV-CC) patterned. *Leucas aspera, Leucas biflora, Salvia leucantha, Salvia splendens,* and *Orthosiphon infortunatum* had the UV-P pattern, where only *Salvia leucantha* was invasive. Plants such as *Anisomeles malabarica, Endostemon viscosus, Gmelina asiatica, Hyptis suaveolens,* and *Salvia coccinea* were UV-CC patterned, where only the last two of the above species were invasive (Table 8).

Plants belonging to the Malvaceae family (N = 12) were either bullseye patterned (UV-BE) or absorbing (UV-A) in nature. All plants of the *Sida* genus were bullseye (UV-BE) patterned, while plants of the genus Hibiscus were all UV absorbing (UV-A) patterned among the Malvaceae family. Plant species such as *Malvaviscus arboreus var. arboreus* (UV-A), *Sida acuta* (UV-BE), and *Waltheria indica* (UV-BE) were invasive.

Among the plants in the Solanaceae family (N = 9), UV contrasting reproductive structure (UV-CR) pattern was found to be most abundant. Plant species such as *Solanum americanum, Solanum erianthum,* and *Solanum torvum* were observed to be invasive and have a contrasting reproductive structure (UV-CR) pattern.

The Nyctaginaceae family (N = 5) of plants were all UV absorbing (UV-A) patterned except for *Mirabilis jalapa* (UV-CR). *Boerhavia coccinea, Bougainvillea spectabilis,* and *Mirabilis jalapa* were observed to be invasive among the Nyctaginaceae family.

The plants in the Verbenaceae family (N = 5) were found to display the bullseye pattern (UV-BE) exclusively where plants such as *Lantana camara*, *Stachytarpheta cayennensis,* and *Stachytarpheta jamaicensis* were invasive among them. The Convolvulaceae family of plants, such as *Ipomoea aquatica, Ipomoea indica,* and *Ipomoea obscura,* were reported to be invasive and have a UV bullseye pattern (UV-BE). Plants of the Balsaminaceae family, *Impatiens balsamina* and *Impatiens walleriana,* were both found to be invasive as well as ultraviolet absorbing (UV-A) patterned.

A group of closely related plants from different families such as *Boswellia serrata* (Burseraceae), *Cardiospermum canescens* (Sapindaceae)*, Cardiospermum corindum* (Sapindaceae), *Ruta graveolens* (Rutaceae), *Toddalia asiatica* (Rutaceae), *Hibiscus calyphyllus* (Malvaceae), *Hibiscus micranthus* (Malvaceae), *Hibiscus vitifolius* (Malvaceae), and *Hibiscus solandra* (Malvaceae) were found to be absorbing (UV-A) and non-invasive. Similarly, another group of plant species from different families, such as *Oxalis corniculata* (Oxalidaceae), *Tribulus terrestris* (Zygophyllaceae), *Brassica rapa* (Brassicaceae)*, Brassica juncea* (Brassicaceae), *Cleome viscosa* (Cleomaceae), and *Waltheria indica* (Malvaceae), were found to have bullseye patterns (UV-BE) and were invasive (Table 8).

### Ultraviolet floral pattern variations within morphs of flower color polymorphic plants

Among the 188 studied species of plants, around 10 species were observed to have flower color polymorphism. Plant species such as *Barleria cristata, Catharanthus roseus, Lantana camara, Mirabilis jalapa, Nerium oleander,* and *Portulaca oleracea* were invasive, and their flowers were flower color polymorphic. Species such as *Bougainvillea glabra, Dahlia pinnata, Hibiscus rosa-sinensis,* and *Tagetes tenuifolia* were non-invasive and had polymorphic flower colors. Among the flower color polymorphic species, *Lantana camara* was found to exhibit different flower colors such as dark-pink, white-pink, orange-red, and red. Flower colors in *L. camara*, such as dark-pink and white-pink, were found to have an ultraviolet bullseye (UV-BE) pattern and fell under the VIB flower color group category. In contrast, orange-red and red flower colors fell under the YOR flower color group category and were found to have an ultraviolet absorbing (UV-A) pattern. The variation in ultraviolet floral pattern among the different flower color morphs in *L. camara* might play a significant role in attracting diverse pollinators (Table 9).

**Table 9:**
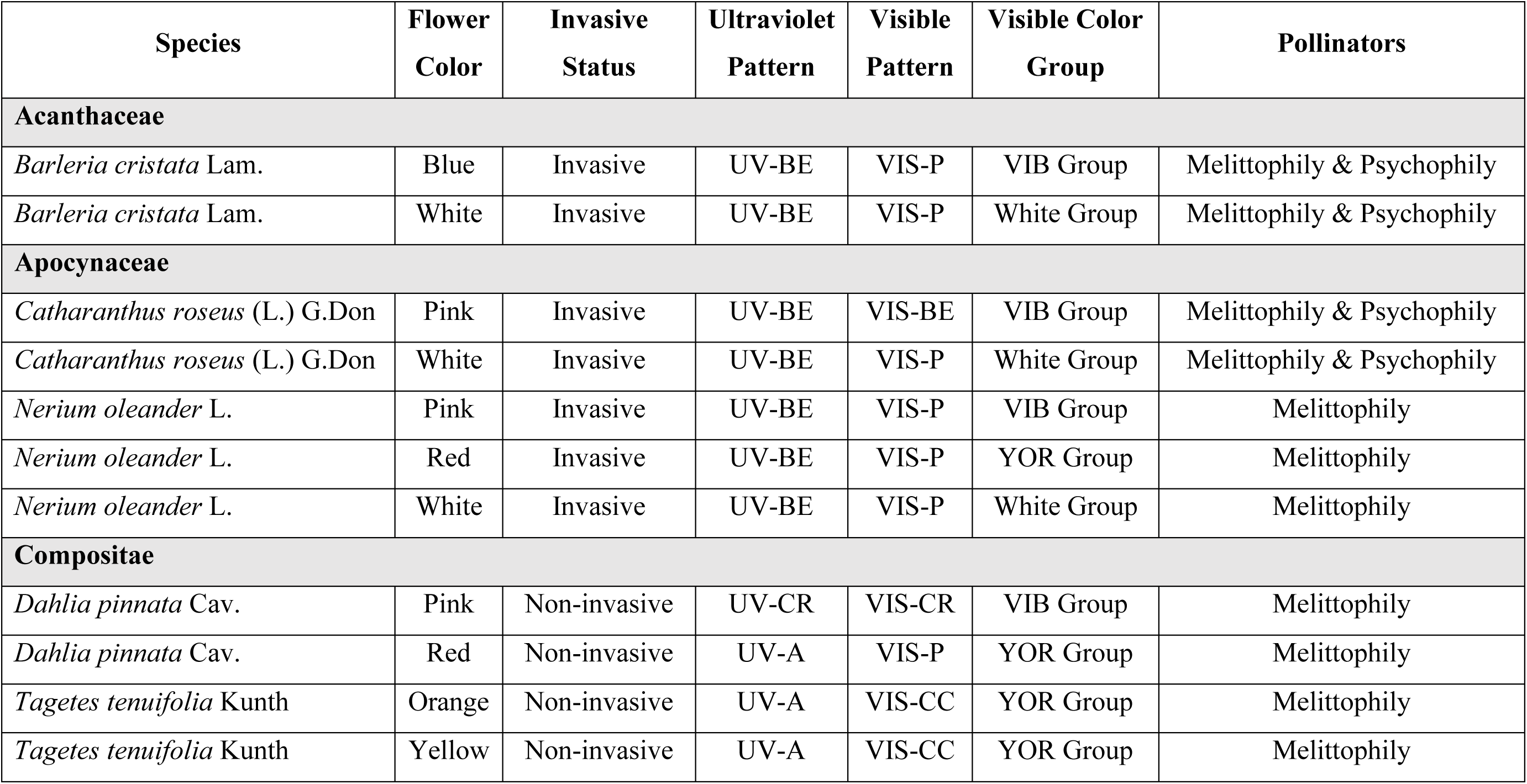
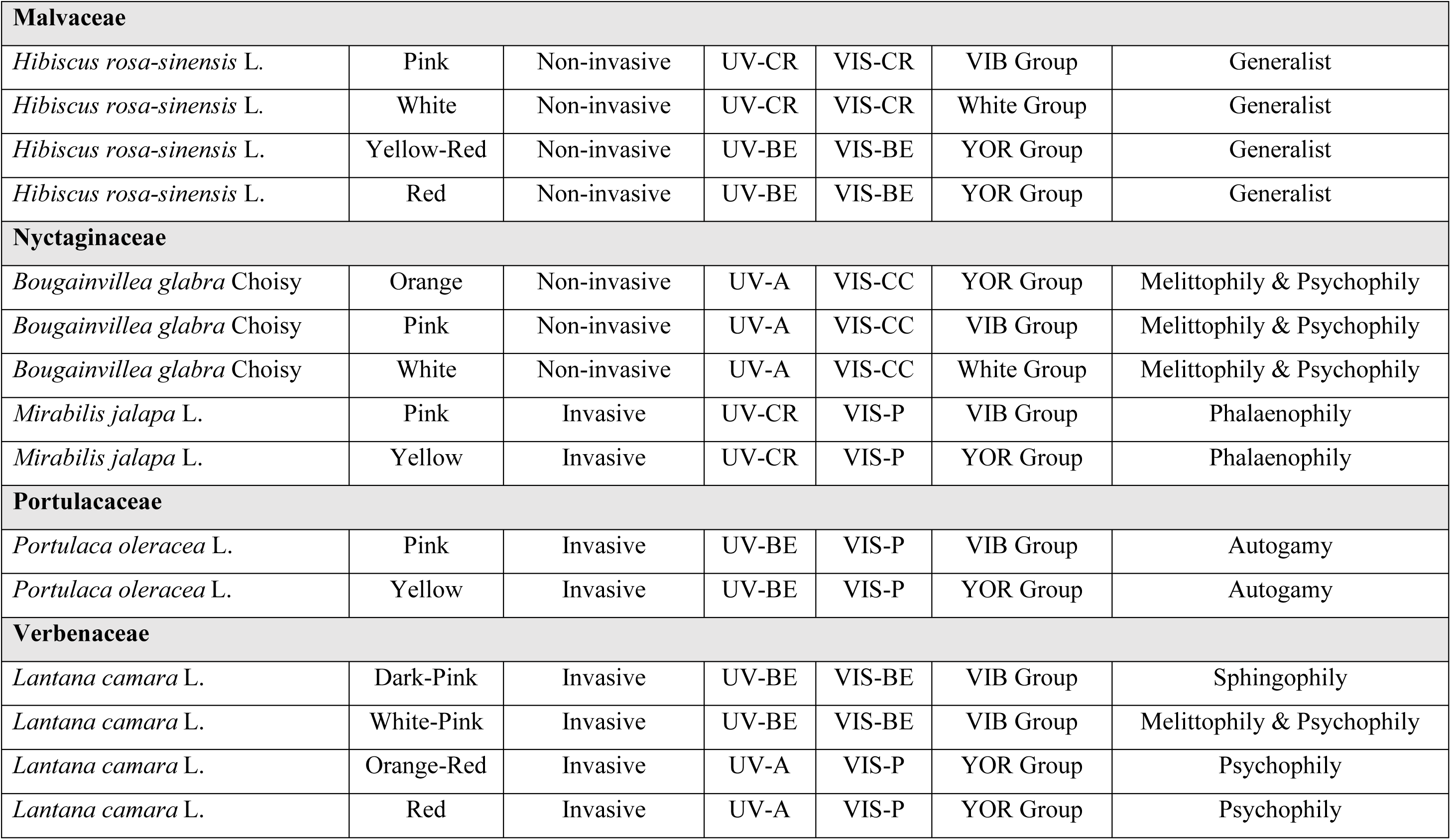
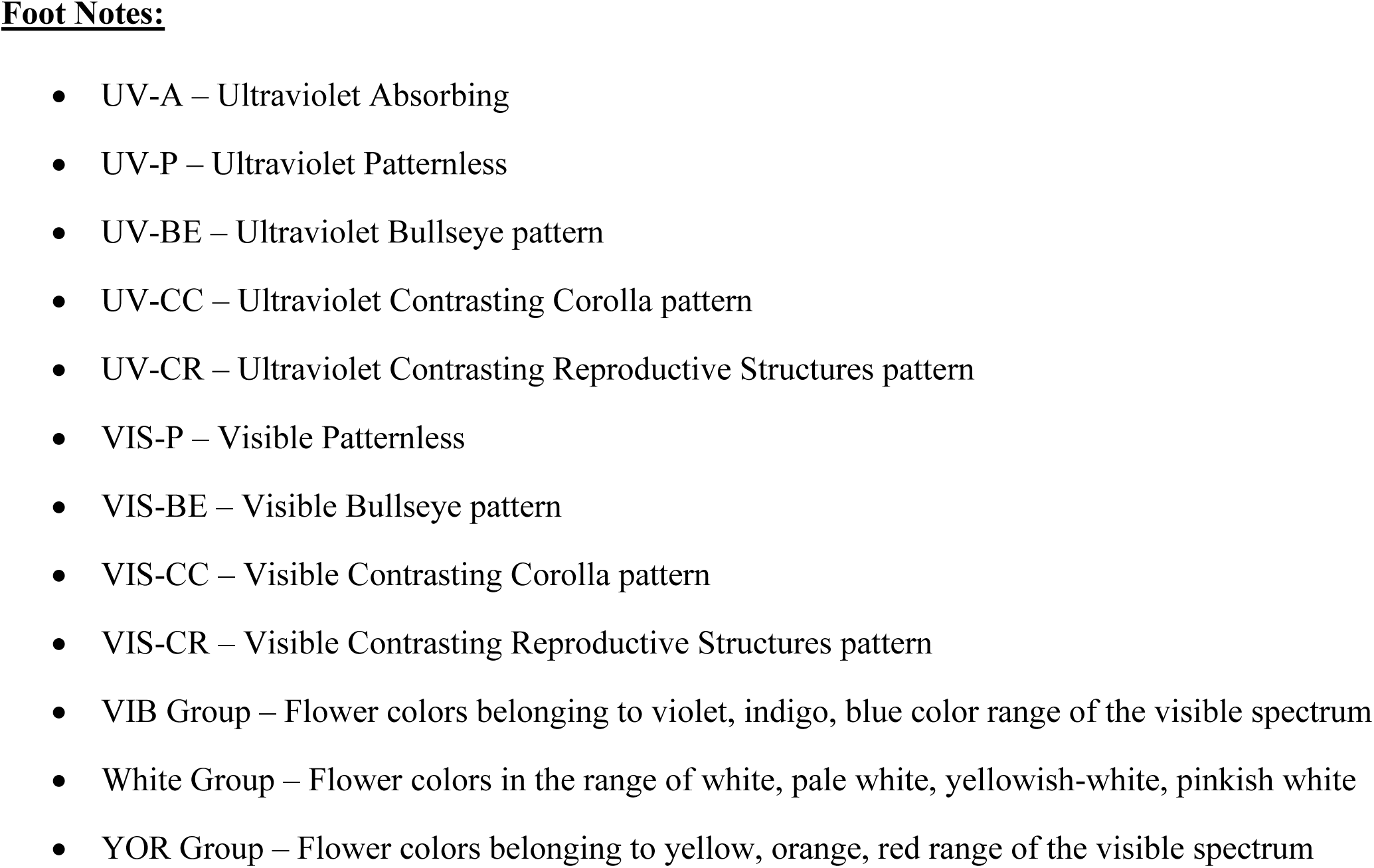
List of studied plant species that exhibit flower color polymorphism (N = 10) along with their respective flower color morph, invasive status, ultraviolet floral pattern, visible floral pattern, and pollinators.

## Discussion

The results from the previous chapter clearly demonstrate that floral reproductive traits might play a vital role in the invasion success of plants. In extension, to test the association between floral patterns (in the visible and ultraviolet spectrums) and pollinators in relation to invasion, this study was conducted among 188 flowering plant species (belonging to 46 families) from parts of the Western Ghats and Eastern Ghats regions of Tamil Nadu. Among the studied plants, 37% of the plant species showed an ultraviolet absorbing (UV-A) pattern, *i.e.,* they were found to absorb most of the ultraviolet radiation between the wavelength ranges of 320–380 nm (Baader-U), thereby being indistinguishable from the background. Another 5% of the plant species were observed to be reflective without any patterns in the ultraviolet spectrum (UV-P). Plants with the UV-P pattern were among the brightest in the UV spectrum due to their almost complete reflection of ultraviolet wavelengths of light. A study in the Neotropical Savanna of Brazil, however, reported that around 66.2% of the studied plants (N = 80) had ultraviolet absorbing (UV-A) and reflecting (UV-P) patterns, and only 33.8% had any distinguishable patterns on their flowers (Tunes et al., 2021). The observations from the present study showed that a majority (58%) of the studied plant species exhibited ultraviolet patterns (Bullseye, Contrasting Corolla, Contrasting Reproductive Structures). The lower relative abundance of various ultraviolet floral patterns in the Neotropical Savanna could be attributed to the latitudinal variations in the intensity of ultraviolet radiation reaching the earth’s surface (Rajpurohit & Schmidt, 2019). The Western Ghats region of Tamil Nadu, a biodiversity hotspot situated in the tropical region, receives abundant sunlight and ultraviolet radiation throughout the year (Harinarayan et al., 2013). A significant variation was observed in the distribution of VIS patterns among UV patterns in the studied plants. Even though 46% of the studied plants had no discernible pattern (VIS-P) in the visible color spectrum, around 50% of the plants in the VIS patternless category were observed to display discernible patterns (UV bullseye, UV contrasting corolla, and UV contrasting reproductive structure) in the ultraviolet spectrum. The present result suggests that a flower may not necessarily display similar patterns in both the ultraviolet and visible spectrums and may utilize different patterns to effectively attract pollinators (Chittka et al., 1994; Koski & Ashman, 2014). The visible spectral floral patterns such as bullseye (VIS-BE), contrasting corolla (VIS-CC), and contrasting reproductive structures (VIS-CR) were found to have the same pattern in the ultraviolet spectrum in the majority of cases (>45%) of studied plant species. Among the different visible patterns, visible patternless (VIS-P) flowers were most frequently found to display other patterns than their similar ones (UV-P) in the ultraviolet spectrum. Floral UV patterns are created by the same floral pigments that are responsible for their visible colors (Thompson et al., 1972). The variations in the ultraviolet patterns among flowers have been reported to be due to the UV absorbing tendency of flavonoids, chalcones, and anthocyanins (Harborne, 1981; Kevan, 1979), whereas carotenoids and anthocyanins have been reported to reflect UV radiation (Kevan et al., 1996; Schlangen et al., 2009).

The relationship between floral patterns in the visible spectrum and pollinators showed no significant association, whereas a significant association was observed in the relationship between ultraviolet floral patterns and different pollinators. The results from the present study suggest that ultraviolet floral patterns play a deterministic role in specific pollinator associations. Plants evolved visual and olfactory cues as a means of communication with pollinators. Visual cues in particular play a vital role in flower identification by pollinators (Briscoe & Chittka, 2001; Kelber et al., 2003). Humans have trichromatic vision, able to perceive light in the red, green, and blue wavelengths with the help of ocular cone cells in the spectrum range of 380 to 750 nanometers; similarly, bees also have trichromatic vision, but in the electromagnetic spectral range of 300 to 600 nanometers, able to perceive ultraviolet, blue, and green colors (Shrestha et al., 2013a). Butterfly vision has been documented to be polychromatic and to be able to perceive many more colors, ranging from ultraviolet to deep reds (van der Kooi et al., 2021). Similarly, birds have also been documented to have tetrachromatic vision and to be able to perceive the ultraviolet spectrum along with the complete visible spectrum (Cuthill et al., 2000; Shrestha et al., 2013b). Other pollinators, such as moths, beetles, and flies, can perceive more than three different colors, including ultraviolet (Kelber et al., 2003; van der Kooi et al., 2021). All these pollinators, specifically bees that play a major role in plant pollination, are highly sensitive in the ultraviolet range (300–400 nm) of the electromagnetic spectrum when compared to the visible colors (Chen et al., 2020; Davies, 2017; van der Kooi et al., 2021). Bee-pollinated plants mostly had contrasting reproductive structures (UV-CR) or bullseye pattern (UV-BE) patterns in the ultraviolet spectrum. Flowers that were pollinated by melittophily & psychophily, or generalist pollinators, were predominantly ultraviolet absorbing (UV-A) in pattern. Similar observations in UV pattern specific pollinator interaction were reported in Hemerocallis flowers, where the bullseye pattern (UV-BE) was pollinated only by bees and not swallowtail butterflies (Hirota et al., 2019), and varied pollinator responses were observed in floral UV reflectance manipulated experiments in *Hypoxis camerooniana* (Klomberg et al., 2019). The different ultraviolet patterns exhibited by plants facilitate specific interactions with different pollinators, irrespective of visible flower colors (Hansen et al., 2012; Horth et al., 2014; Papiorek et al., 2016).

The comparison of the different visible flower color groups (VIB, White, and YOR) associations with different classes of pollinators shows melittophily (34%) to be the most preferred mode of pollination, followed by generalists (27%), and entomophily (20%) among the studied plant species. About 46% of the studied plant species had no discernible patterns (VIS-P) in the visible spectrum, compared to 5% in the ultraviolet spectrum (UV-P). Both the VIS-P and UV-P patterns were predominant in the White color group, which is associated with generalists and melittophily & psychophily-mediated pollination. Among all the flower color groups (VIB, White, and YOR), plants with no visible pattern (VIS-P) were found to be predominant. The number of flowers that display no visible pattern was mostly yellow, white, or whiteish-yellow in color. Similar works have indicated that patternless yellow or white colored flowers frequently had ultraviolet patterns perceivable only to their intended pollinators (Hansen et al., 2012; Papiorek et al., 2016). The proportions of plant distribution under the different flower color group categories (VIB, White, and YOR) were found to be almost similar among all the studied visible patterns.

The contribution to the ultraviolet absorbing (UV-A) pattern was greater among the plants in the White flower color group. A similar study in the Neotropical Savanna found an abundance of ultraviolet absorbing patterns among the plant species studied (Tunes et al., 2021). Flowers in the White color groups were pollinated almost equally by generalist and both melittophily & psychophily. Plants from the VIB and YOR flower color groups were almost equally distributed toward ultraviolet absorbing (UV-A), ultraviolet bullseye (UV-BE), and ultraviolet contrasting reproductive structures (UV-CR) patterns. Most flowers in the VIB flower color groups were bicolored with contrasting markings on their petals that served as nectar guides (Leonard & Papaj, 2011). Concurrently, 40% and 38% of the VIB and YOR flower color groups, respectively, were primarily associated with melittophily and secondarily with melittophily & psychophily-mediated pollination. The results indicate that flower color groups such as VIB and YOR, where ultraviolet bullseye (UV-BE) and ultraviolet contrasting reproductive structures (UV-CR) together are more prominent when compared to White color groups, are primarily pollinated by bees (Melittophily).

Around 43% of the studied plant species (N = 188) collected in this study were found to be invasive. This is in concurrence with the results from the recent report from Tamil Nadu Policy on Invasive Plants and Ecological Restoration (TN PIPER) showed that around 36.6% of the recorded 6723 species of plants in Tamil Nadu were reported to be invasive, specifically concentrated in and around the Western Ghats region of the state (Tamil Nadu Policy on Invasive Alien Plant Species and Ecological Restoration of Habitats Short Title Tamil Nadu Policy on Invasive Plants and Ecological Restoration (TN PIPER, 2021). About 49% of India’s total geographic area has been predicted to be invasion prone (Adhikari et al., 2015). Observation of the flower colors among the invasive and non-invasive species of the studied plants shows that invasive species predominantly had flower colors in the YOR and VIB ranges of the visible color spectrum. Non-invasive plants (N = 108) were predominantly found to have flower color in the White group. This result is consistent with the findings from the previous chapter, where the YOR flower color group was predominant in invasive plants and the White flower color group was greater among non-invasive plants from across the world.

Both invasive and non-invasive plant species were observed to predominantly display ultraviolet absorbing (UV-A) and ultraviolet bullseye (UV-BE) patterns in the UV spectrum. Ultraviolet bullseye (UV-BE) and ultraviolet absorbing (UV-A) patterns have most commonly been associated with bee-and butterfly-mediated pollination (Chittka et al., 1994; Hirota et al., 2019; Horth et al., 2014). There was a significant variation among the ultraviolet patterns and pollinators in relation to the invasion status of the studied plants, but no variation was observed between visible patterns and pollinators in relation to the invasion status. Invasive plants have been found to benefit bees due to the year-round availability of flowers (Kovács-Hostyánszki et al., 2022), and butterflies and bees have been found to be better integrated among the invasive plant population (Anu et al., 2021; Vilà et al., 2009). This finding is consistent with the findings of Saidanyan et al. (2016), who discovered that majority of the invasive plant species have multi-seasonal flowering and flower color polymorphism. The results showed that invasive plants were associated with a greater diversity of pollinators when compared to non-invasive plants. Further analysis revealed that invasive species with flower colors in the VIB and YOR ranges of the visible spectrum along with ultraviolet absorbing (UV-A) and ultraviolet bullseye (UV-BE) patterns were most associated with bee-and butterfly-mediated pollination (Horth et al., 2014; Papiorek et al., 2016; Peterson et al., 2015; Tunes et al., 2021). Whereas non-invasive plants from the study were observed to be mostly associated with the White flower color group and pollinated by non-specific insects and bees in plants from around the Western Ghats region (Devy & Davidar, 2003; Hobbhahn et al., 2006; Kuriakose et al., 2018). The results suggest that specific colors and ultraviolet patterns displayed by invasive species might result in a pollinator selection driven reproductive advantage over non-invasive species of plants.

The phylogeny of plants in relation to their visible flower color showed that there was no specific phylogenetic link between the two and that the flower color trait is variable (Bergamo et al., 2018; McEwen & Vamosi, 2010; Shrestha et al., 2013b). Plant families such as Leguminosae, Compositae, Lamiaceae, Solanaceae, Apocynaceae, and Convolvulaceae (listed in descending order of invasive species abundance) had the most abundance in terms of invasive species among the studied plants. Results from the present study showed that a significant degree of phylogenetic signal was observed between ultraviolet floral patterns and plant species phylogeny. Among the 80 invasive species in the study, 33% were found to exhibit ultraviolet absorbing (UV-A) patterns, and another 33% were found to exhibit ultraviolet bullseye (UV-BE) patterns that were predominant among them. Plants belonging to the Compositae and Leguminosae families were found to contribute to about half of the ultraviolet absorbing (UV-A) floral pattern among the invasive plants. Among invasive plants, almost half of the plants with an ultraviolet bullseye (UV-BE) pattern were found to be in the Apocynaceae, Compositae, Convolvulaceae, and Verbenaceae (N = 3) plant families. The invasive plants belonging to the Lamiaceae family were found to display all ultraviolet floral patterns except for ultraviolet bullseye (UV-BE). The distribution of ultraviolet floral patterns (UV-A, UV-BE, and UV-CR) among invasive species in the Apocynaceae and Compositae plant families was found to be similar. Ultraviolet floral patterns were reported to be phylogenetically conserved among the studied plant (N = 80) species in the Neotropical savanna of Brazil (Tunes et al., 2021). Invasive species have been found to be phylogenetically conserved among the angiosperm species, where highly harmful species of invasive plants were reported to be strongly clustered (Divíšek et al., 2018; Qian et al., 2022). To conclude, plant species collected from around the Western Ghats and Eastern Ghats regions of Tamil Nadu revealed that invasive plant species from the same plant family were most likely to display similar floral UV patterns.

Invasive plant species have been reported to have multi-seasonal flowering and exhibit flower color polymorphism (Saidanyan et al., 2016). Plant species that exhibit flower color polymorphism were found to maintain colors in different flower color groups. Flower color polymorphic plants in the study (N = 10) were observed to maintain similar visible patterns, ultraviolet floral patterns, and pollinators among the species, except in the case of the invasive species *Lantana camara*. Among the different flowers of *L. camara*, dark-pink and white-pink morphs showed UV bullseye (UV-BE) floral pattern, whereas orange-red and red flower colors showed UV absorbing (UV-A) floral pattern. *L. camara* has been reported to show variations in the floral colors with respect to their pollination status (Lunau, 1996; Ram & Mathur, 1984; Weiss, 1991). The different flower color morphs of *L. camara* studied (dark-pink, white-pink, orange-red, and red), were found to be pollinated by different categories of pollinators (Andersson & Dobson, 2003; Willmer et al., 2009). The orange-red and red flower color morphs of *L. camara* were found to be pollinated by psychophily, whereas the dark-pink and white-pink morphs of *L. camara* were found to be pollinated by sphingophily and melittophily, respectively. This indicates that plant species that could employ different ultraviolet patterns might hold a reproductive advantage over plant species that exhibit only one UV floral pattern. The purpose of plant signaling towards pollinators is to direct them towards unpollinated flowers with floral rewards (Davies et al., 2022; Koski & Ashman, 2014; Lunau, 1992; Sooraj et al., 2019), and prevent energy spent by pollinators in visiting already pollinated flowers with little or no reward (Lucas-Barbosa et al., 2016; Papiorek et al., 2016; Primack, 1982; Young et al., 1991).

In conclusion, the results from the present study showed that there was no significant difference among visible (VIS) patterns and pollinators with relation to invasion status. A significant variation was observed between the ultraviolet (UV) patterns and pollinators among the invasive plants, whereas no such significant variation was observed in the case of non-invasive species. This result clearly suggests that the variation in the floral ultraviolet signature among invasive species plays a vital role in plant-pollinator interaction and invasion success. A significant degree of phylogenetic signal was observed between ultraviolet floral patterns and plant species phylogeny. The ultraviolet floral patterns were also found to vary among the different flower color morphs in *L. camara*. There is still a lacuna in our understanding of the variations in the ultraviolet signature and floral metabolites. Thus, the following chapter is an attempt to address the temporal variations in the ultraviolet patterns with respect to pollination status and floral metabolite variations among the flower color morphs of *L. camara*.

## References

1. Adhikari, D., Tiwary, R., & Barik, S. K. (2015). Modelling hotspots for invasive alien plants in India. PLoS ONE, 10(7), e0134665. https://doi.org/10.1371/JOURNAL.PONE.0134665

2. Albrecht, M., Ramis, M. R., & Traveset, A. (2016). Pollinator-mediated impacts of alien invasive plants on the pollination of native plants: the role of spatial scale and distinct behaviour among pollinator guilds. Biological Invasions, 18(7), 1801–1812. https://doi.org/10.1007/S10530-016-1121-6/FIGURES/5

3. Andersson, S., & Dobson, H. E. M. (2003). Behavioral foraging responses by the butterfly Heliconius melpomene to Lantana camara floral scent. Journal of Chemical Ecology, 29(10), 2303–2318. https://doi.org/10.1023/A:1026226514968

4. Anu, B., & Vishwakarma, R. (2021). Invasive Exotic Plant-Pollinator Interactions. In A. Rustagi, & B. Chaudhry (Eds.), Plant Reproductive Ecology - Recent Advances. IntechOpen. https://doi.org/10.5772/intechopen.100895

5. Aravindhan, V., & Rajendran, A. (2014). Impact of invasive species Lantana camara (L.) on the vegetation of Velliangiri Hills, the Southern Western Ghats, India. Global Journal of Environmental Research, 8(3), 35–40. https://doi.org/10.5829/idosi.gjer.2014.8.3.1107

6. Balaguru, B., Soosairaj, S., Nagamurugan, N., Ravindran, R., & Ahamed Khaleel, A. (2016). Native vegetation pattern and the spread of three invasive species in Palani Hill National Park, Western Ghats of India. Acta Ecologica Sinica, 36(5), 367–376. https://doi.org/10.1016/J.CHNAES.2016.05.005

7. Beans, C. M., & Roach, D. A. (2015). An invasive plant alters pollinator-mediated phenotypic selection on a native congener. American Journal of Botany, 102(1), 50–57. https://doi.org/10.3732/AJB.1400385

8. Bergamo, P. J., Telles, F. J., Arnold, S. E. J., & de Brito, V. L. G. (2018). Flower colour within communities shifts from overdispersed to clustered along an alpine altitudinal gradient. Oecologia, 188(1), 223–235. https://doi.org/10.1007/S00442-018-4204-5/TABLES/4

9. Boynton, R.M. (1990). Human color perception. In: Leibovic, K.N. (eds) Science of Vision. Springer, New York, NY. https://doi.org/10.1007/978-1-4612-3406-7_8

10. Briscoe, A. D., & Chittka, L. (2001). The evolution of color vision in insects. Annual Review of Entomology, 46, 471–510. https://doi.org/10.1146/ANNUREV.ENTO.46.1.471

11. CABI Compendium Invasive Species. (2022). CABI Digital Library. https://www.cabidigitallibrary.org/product/qi

12. Chandrasekaran, S., & Swamy, P. S. (2010). Growth patterns of Chromolaena odorata in varied ecosystems at Kodayar in the Western Ghats, India. Acta Oecologica, 36(4), 383–392. https://doi.org/10.1016/J.ACTAO.2010.03.006

13. Chandrasekaran, S., & Swamy, P. S. (2002). Biomass, litterfall and aboveground net primary productivity of herbaceous communities in varied ecosystems at Kodayar in the western ghats of Tamil Nadu. Agriculture, Ecosystems and Environment, 88(1), 61–71. https://doi.org/10.1016/S0167-8809(01)00162-1

14. Chandrasekaran, S., Sundarapandian, S. M., Chandrasekar, P., & Swamy, P. S. (2001). Exotic plant invasions in disturbed and man-modified forest ecosystems in the Western Ghats of Tamil Nadu. Tropical Forestry Research: Challenges in the New Millennium. Proceedings of the International Symposium, Peechi, India, 2-4 August, 2000, 32–39.

15. Chandrasekaran, S., & Swamy, P. S. (1995). Changes in herbaceous vegetation following disturbance due to biotic interference in natural and man-made ecosystems in Western ghats. Tropical Ecology, 36(2), 213–220. https://eurekamag.com/research/008/283/008283548.php

16. Chen, Z., Liu, C. Q., Sun, H., & Niu, Y. (2020). The ultraviolet colour component enhances the attractiveness of red flowers of a bee-pollinated plant. Journal of Plant Ecology, 13(3), 354–360. https://doi.org/10.1093/JPE/RTAA023

17. Chittka, L., Shmida, A., Troje, N., & Menzel, R. (1994). Ultraviolet as a component of flower reflections, and the colour perception of hymenoptera. Vision Research, 34(11), 1489–1508. https://doi.org/10.1016/0042-6989(94)90151-1

18. Chittka, L., & Menzel, R. (1992). The evolutionary adaptation of flower colours and the insect pollinators’ colour vision. Journal of Comparative Physiology A 1992 171:2, 171(2), 171–181. https://doi.org/10.1007/BF00188925

19. Cuthill, I. C., Partridge, J. C., Bennett, A. T. D., Church, S. C., Hart, N. S., & Hunt, S. (2000). Ultraviolet vision in birds. Advances in the Study of Behavior, 29, 159–214. https://doi.org/10.1016/S0065-3454(08)60105-9

20. Data and digital resources | Kew. (2022). www.kew.org. https://www.kew.org/science/collections-and-resources/data-and-digital

21. Davies, K. M., Landi, M., van Klink, J. W., Schwinn, K. E., Brummell, D. A., Albert, N. W., Chagné, D., Jibran, R., Kulshrestha, S., Zhou, Y., & Bowman, J. L. (2022). Evolution and function of red pigmentation in land plants. Annals of Botany, 130(5), 613–636. https://doi.org/10.1093/AOB/MCAC109

22. Davis, E. S., Kelly, R., Maggs, C. A., & Stout, J. C. (2018). Contrasting impacts of highly invasive plant species on flower-visiting insect communities. Biodiversity and Conservation, 27(8), 2069– 2085. https://doi.org/10.1007/S10531-018-1525-Y/FIGURES/5

23. Davies, A. (2017). Digital Ultraviolet and Infrared Photography (1st ed.). Routledge, New York https://doi.org/10.4324/9781315515090

24. Devy, M. S., & Davidar, P. (2003). Pollination systems of trees in Karachi, a mid-elevation wet evergreen forest in Western Ghats, India. American Journal of Botany, 90(4), 650–657. https://doi.org/10.3732/AJB.90.4.650

25. Divíšek, J., Chytrý, M., Beckage, B., Gotelli, N. J., Lososová, Z., Pyšek, P., Richardson, D. M., & Molofsky, J. (2018). Similarity of introduced plant species to native ones facilitates naturalization, but differences enhance invasion success. Nature Communications, 9(1). https://doi.org/10.1038/S41467-018-06995-4

26. Dyer, A. G., Boyd-Gerny, S., Mcloughlin, S., Rosa, M. G. P., Simonov, V., & Wong, B. B. M. (2012). Parallel evolution of angiosperm colour signals: Common evolutionary pressures linked to hymenopteran vision. Proceedings of the Royal Society B: Biological Sciences, 279(1742), 3606– 3615. https://doi.org/10.1098/RSPB.2012.0827

27. Flowers of India. (2022). https://www.flowersofindia.net/

28. Garcia, J. E., Greentree, A. D., Shrestha, M., Dorin, A., & Dyer, A. G. (2014). Flower colours through the lens: Quantitative measurement with visible and ultraviolet digital photography. PLoS ONE, 9(5), e96646. https://doi.org/10.1371/JOURNAL.PONE.0096646

29. Gibson, M. R., Richardson, D. M., & Pauw, A. (2012). Can floral traits predict an invasive plant’s impact on native plant–pollinator communities? Journal of Ecology, 100(5), 1216–1223. https://doi.org/10.1111/J.1365-2745.2012.02004.X

30. GISD. (2019). Iucngisd.org. http://www.iucngisd.org/gisd/

31. Go Botany: Native Plant Trust. (2019). Nativeplanttrust.org. https://gobotany.nativeplanttrust.org/

32a. Godar, D. E., Landry, R. J., & Lucas, A. D. (2009). Increased UVA exposures and decreased cutaneous

32. Vitamin D3 levels may be responsible for the increasing incidence of melanoma. Medical Hypotheses, 72(4), 434–443. https://doi.org/10.1016/J.MEHY.2008.09.056

33. Goodell, K., & Parker, I. M. (2017). Invasion of a dominant floral resource: effects on the floral community and pollination of native plants. Ecology, 98(1), 57–69. https://doi.org/10.1002/ECY.1639

34. Gronquist, M., Bezzerides, A., Attygalle, A., Meinwald, J., Eisner, M., & Eisner, T. (2001). Attractive and defensive functions of the ultraviolet pigments of a flower (Hypericum calycinum). Proceedings of the National Academy of Sciences, 98(24), 13745–13750. https://doi.org/10.1073/PNAS.231471698

35. Hansen, D. M., van der Niet, T., & Johnson, S. D. (2012). Floral signposts: testing the significance of visual ‘nectar guides’ for pollinator behaviour and plant fitness. Proceedings of the Royal Society B: Biological Sciences, 279(1729), 634–639. https://doi.org/10.1098/RSPB.2011.1349

36. Harborne, J. B. (1981). Two gossypetin methyl ethers as ultraviolet patterning guides in the flowers of Coronilla valentina. Phytochemistry, 20(5), 1117–1119. https://doi.org/10.1016/0031-9422(81)83038-5

37. Harinarayan, C. V., Holick, M. F., Prasad, U. V., Vani, P. S., & Himabindu, G. (2013). Vitamin D status and sun exposure in India. Dermato-Endocrinology, 5(1), 130. https://doi.org/10.4161/DERM.23873

38. Hirota, S. K., Miki, N., Yasumoto, A. A., & Yahara, T. (2019). UV bullseye contrast of Hemerocallis flowers attracts hawkmoths but not swallowtail butterflies. Ecology and Evolution, 9(1), 52. https://doi.org/10.1002/ECE3.4604

39. Hobbhahn, N., Küchmeister, H., & Porembski, S. (2006). Pollination biology of mass flowering terrestrial Utricularia species (lentibulariaceae) in the Indian Western Ghats. Plant Biology (Stuttgart, Germany), 8(6), 791–804. https://doi.org/10.1055/S-2006-924566

40. Horth, L., Campbell, L., & Bray, R. (2014). Wild bees preferentially visit Rudbeckia flower heads with exaggerated ultraviolet absorbing floral guides. Biology Open, 3(3), 221–230. https://doi.org/10.1242/BIO.20146445

41. Jin, Y., & Qian, H. (2019). V.PhyloMaker: an R package that can generate very large phylogenies for vascular plants. Ecography, 42(8), 1353–1359. https://doi.org/10.1111/ecog.04434

42. Karanth, K. K., Sankararaman, V., Dalvi, S., Srivathsa, A., Parameshwaran, R., Sharma, S., Robbins, P., & Chhatre, A. (2016). Producing diversity: Agroforests sustain avian richness and abundance in India’s Western Ghats. Frontiers in Ecology and Evolution, 4(SEP), 111. https://doi.org/10.3389/FEVO.2016.00111/BIBTEX

43. Kelber, A., Vorobyev, M., & Osorio, D. (2003). Animal colour vision - Behavioural tests and physiological concepts. Biological Reviews of the Cambridge Philosophical Society, 78(1), 81–118. https://doi.org/10.1017/S1464793102005985

44. Keller, R. P., Geist, J., Jeschke, J. M., & Kühn, L. (2011). Invasive species in Europe: Ecology, status, and policy. Environmental Sciences Europe, 23(1), 1–17. https://doi.org/10.1186/2190-4715-23-23/FIGURES/1

45. Kevan, P., Giurfa, M., & Chittka, L. (1996). Why are there so many and so few white flowers? Trends in Plant Science, 1(8), 252. https://doi.org/10.1016/1360-1385(96)20008-1

46. Kevan, P. G. (1979). Vegetation and floral colors revealed by ultraviolet light: Interpretational difficulties for functional significance. American Journal of Botany, 66(6), 749–751. https://doi.org/10.1002/J.1537-2197.1979.TB06280.X

47. KFRI - KFSTAT Database. (2022). https://www.kfri.res.in/kfri_database.asp

48. King, V. M., & Sargent, R. D. (2012). Presence of an invasive plant species alters pollinator visitation to a native. Biological Invasions, 14(9), 1809–1818. https://doi.org/10.1007/S10530-012-0191-3/FIGURES/4

49. Klomberg, Y., Dywou Kouede, R., Bartoš, M., Mertens, J. E. J., Tropek, R., Fokam, E. B., & Janeček, Š. (2019). The role of ultraviolet reflectance and pattern in the pollination system of Hypoxis camerooniana (Hypoxidaceae). AoB PLANTS, 11(5). https://doi.org/10.1093/AOBPLA/PLZ057

50. Koski, M. H. (2020). Macroevolution of flower color patterning: Biased transition rates and correlated evolution with flower size. Frontiers in Plant Science, 11, 945. https://doi.org/10.3389/fpls.2020.00945

51. Koski, M. H., MacQueen, D., & Ashman, T. L. (2020). Floral pigmentation has responded rapidly to global change in ozone and temperature. Current Biology, 30(22), 4425–4431.e3. https://doi.org/10.1016/J.CUB.2020.08.077

52. Koski, M. H., & Ashman, T. (2016). Macroevolutionary patterns of ultraviolet floral pigmentation explained by geography and associated bioclimatic factors. New Phytologist, 211(2), 708–718. https://doi.org/10.1111/nph.13921

53. Koski, M. H., & Ashman, T. L. (2014). Dissecting pollinator responses to a ubiquitous ultraviolet floral pattern in the wild. Functional Ecology, 28(4), 868–877. https://doi.org/10.1111/1365-2435.12242

54. Kovács-Hostyánszki, A., Szigeti, V., Miholcsa, Z., Sándor, D., Soltész, Z., Török, E., & Fenesi, A. (2022). Threats and benefits of invasive alien plant species on pollinators. Basic and Applied Ecology, 64, 89–102. https://doi.org/10.1016/J.BAAE.2022.07.003

55. Kuriakose, G., Sinu, P. A., & Shivanna, K. R. (2018). Ant pollination of Syzygium occidentale, an endemic tree species of tropical rain forests of the Western Ghats, India. Arthropod-Plant Interactions, 12(5), 647–655. https://doi.org/10.1007/S11829-018-9613-1/FIGURES/5

56. Leonard, A. S., & Papaj, D. R. (2011). ‘X’ marks the spot: The possible benefits of nectar guides to bees and plants. Functional Ecology, 25(6), 1293–1301. https://doi.org/10.1111/J.1365-2435.2011.01885.X

57. Lucas-Barbosa, D., Sun, P., Hakman, A., van Beek, T. A., van Loon, J. J. A., & Dicke, M. (2016). Visual and odour cues: plant responses to pollination and herbivory affect the behaviour of flower visitors. Functional Ecology, 30(3), 431–441. https://doi.org/10.1111/1365-2435.12509

58. Lunau, K. (1996). Unidirectionality of floral colour changes. Plant Systematics and Evolution, 200(1), 125–140. https://doi.org/10.1007/BF00984753

59. Lunau, K. (1992). A new interpretation of flower guide colouration: Absorption of ultraviolet light enhances colour saturation. Plant Systematics and Evolution, 183(1), 51–65. https://doi.org/10.1007/BF00937735

60. Matthew, K. M. (1981). Flora of the Tamil Nadu Carnatic. Rapinat Herbarium, St. Joseph’s College. https://doi.org/10.3/JQUERY-UI.JS

61. McEwen, J. R., & Vamosi, J. C. (2010). Floral colour versus phylogeny in structuring subalpine flowering communities. Proceedings of the Royal Society B: Biological Sciences, 277(1696), 2957– 2965. https://doi.org/10.1098/RSPB.2010.0501

62. McKenzie, R., Smale, D., Bodeker, G. and Claude, H. (2003). Ozone profile differences between Europe and New Zealand: Effects on surface UV irradiance and its estimation from satellite sensors. Journal of Geophysical Research, 108(D6), 4179. https://doi.org/10.1029/2002JD002770

63. Milet-Pinheiro, P., Ayasse, M., & Dötterl, S. (2015). Visual and olfactory floral cues of Campanula (Campanulaceae) and their significance for host recognition by an oligolectic bee pollinator. PLoS ONE, 10(6). https://doi.org/10.1371/JOURNAL.PONE.0128577

64. Moyroud, E., & Glover, B. J. (2017). The physics of pollinator attraction. New Phytologist, 216(2), 350–354. https://doi.org/10.1111/NPH.14312

65. Napoleone, F., Manzini, D., & Burrascano, S. (2022). How to measure flower ultraviolet reflectance using digital photography. Applied Vegetation Science, 25(1), e12648. https://doi.org/10.1111/AVSC.12648

66. Narbona, E., Arista, M., Whittall, J. B., Camargo, M. G. G., & Shrestha, M. (2021a). Editorial: The role of flower color in Angiosperm evolution. Frontiers in Plant Science, 12, 736998. https://doi.org/10.3389/FPLS.2021.736998

67. Narbona, E., del Valle, J. C., Arista, M., Buide, M. L., & Ortiz, P. L. (2021b). Major flower pigments originate different colour signals to pollinators. Frontiers in Ecology and Evolution, 9, 686. https://doi.org/10.3389/FEVO.2021.743850/BIBTEX

68. Ödeen, A., & Håstad, O. (2010). Pollinating birds differ in spectral sensitivity. Journal of Comparative Physiology A: Neuroethology, Sensory, Neural, and Behavioral Physiology, 196(2), 91–96. https://doi.org/10.1007/S00359-009-0474-Z/FIGURES/1

69. Origin Pro (2021). OriginLab Corporation, Northampton, MA, USA.

70. Papiorek, S., Junker, R. R., Alves-dos-Santos, I., Melo, G. A. R., Amaral-Neto, L. P., Sazima, M., Wolowski, M., Freitas, L., & Lunau, K. (2016). Bees, birds and yellow flowers: pollinator-dependent convergent evolution of UV patterns. Plant Biology, 18(1), 46–55. https://doi.org/10.1111/PLB.12322

71. Papiorek, S., Rohde, K., & Lunau, K. (2013). Bees’ subtle colour preferences: How bees respond to small changes in pigment concentration. Naturwissenschaften, 100(7), 633–643. https://doi.org/10.1007/S00114-013-1060-3/TABLES/2

72. Parra-Tabla, V., & Arceo-Gómez, G. (2021). Impacts of plant invasions in native plant–pollinator networks. New Phytologist, 230(6), 2117–2128. https://doi.org/10.1111/NPH.17339

73. Peterson, M. L., Miller, T. J., & Kay, K. M. (2015). An ultraviolet floral polymorphism associated with life history drives pollinator discrimination in Mimulus guttatus. American Journal of Botany, 102(3), 396–406. https://doi.org/10.3732/AJB.1400415

74. Primack, R. (1982). Ultraviolet patterns in flowers, or flowers as viewed by insects. Arnoldia (USA), 42(3), 139–146.

75. R Core Team (2022). R: A language and environment for statistical computing. R Foundation for Statistical Computing, Vienna, Austria. URL https://www.R-project.org/.

76. Raguso, R. A., & Willis, M. A. (2002). Synergy between visual and olfactory cues in nectar feeding by naive hawkmoths, Manduca sexta. Animal Behaviour, 64(5), 685–695. https://doi.org/10.1006/ANBE.2002.4010

77. Rajpurohit, S., & Schmidt, P. S. (2019). Latitudinal pigmentation variation contradicts ultraviolet radiation exposure: A case study in tropical Indian Drosophila melanogaster. Frontiers in Physiology, 10(FEB), 84. https://doi.org/10.3389/FPHYS.2019.00084/BIBTEX

78. Ram, H. Y. M., & Mathur, G. (1984). Flower colour changes in Lantana camara. Journal of Experimental Botany, 35(11), 1656–1662. https://doi.org/10.1093/JXB/35.11.1656

79. Saidanyan, R. I., Kamaladhasan, N., Krishnankutty, N., & Chandrasekaran, S. (2016). Influence of flower colour and seasonality on plant invasion success. Current Science, 111(4), 617–619.

80. Schlangen, K., Miosic, S., Castro, A., Freudmann, K., Luczkiewicz, M., Vitzthum, F., Schwab, W., Gamsjäger, S., Musso, M., & Halbwirth, H. (2009). Formation of UV-honey guides in Rudbeckia hirta. Phytochemistry, 70(7), 889–898. https://doi.org/10.1016/J.PHYTOCHEM.2009.04.017

81. Schnapf, J. L., Kraft, T. W., & Baylor, D. A. (1987). Spectral sensitivity of human cone photoreceptors. Nature, 325(6103), 439–441. https://doi.org/10.1038/325439a0

82. Schulte, A. J., Mail, M., Hahn, L. A., & Barthlott, W. (2019). Ultraviolet patterns of flowers revealed in polymer replica - caused by surface architecture. Beilstein Journal of Nanotechnology, 10(1), 459–466. https://doi.org/10.3762/bjnano.10.45

83. Shrestha, M., Dyer, A. G., & Burd, M. (2013a). Evaluating the spectral discrimination capabilities of different pollinators and their effect on the evolution of flower colors. Communicative & Integrative Biology, 6(3), e24000. https://doi.org/10.4161/CIB.24000

84. Shrestha, M., Dyer, A. G., Boyd-Gerny, S., Wong, B. B. M., & Burd, M. (2013b). Shades of red: Bird-pollinated flowers target the specific colour discrimination abilities of avian vision. New Phytologist, 198(1), 301–310. https://doi.org/10.1111/NPH.12135

85. Sooraj, N. P., Jaishanker, R., Athira, K., Sajeev, C. R., Lijimol, D., Saroj, K.V., Ammini, J., Pillai, M. S., & Dadhwal, V. K. (2019). Comparative study on the floral spectral reflectance of invasive and non-invasive plants. Ecological Informatics, 53, 100990. https://doi.org/10.1016/J.ECOINF.2019.100990

86. Swamy, P. S., Sundarapandian, S. M., Chandrasekar, P., & Chandrasekaran, S. (2000). Plant species diversity and tree population structure of a humid tropical forest in Tamil Nadu, India. Biodiversity and Conservation, 9(12), 1643–1669. https://doi.org/10.1023/A:1026511812878

87. Thompson, W. R., Meinwald, J., Aneshansley, D., & Eisner, T. (1972). Flavonols: pigments responsible for ultraviolet absorption in nectar guide of flower. Science (New York, N.Y.), 177(4048), 528–530. https://doi.org/10.1126/SCIENCE.177.4048.528

88. Trunschke, J., Lunau, K., Pyke, G. H., Ren, Z. X., & Wang, H. (2021). Flower color evolution and the evidence of pollinator-mediated selection. Frontiers in Plant Science, 12, 1096. https://doi.org/10.3389/FPLS.2021.617851/BIBTEX

89. TRY Plant Trait Database. (2022). https://www.try-db.org/TryWeb/Home.php

90. Tunes, P., Camargo, M. G. G., & Guimarães, E. (2021). Floral UV features of plant species from a neotropical Savanna. Frontiers in Plant Science, 12, 771. https://doi.org/10.3389/fpls.2021.618028

91. Useful Tropical Plants. (2022). Tropical.theferns.info. https://tropical.theferns.info/

92. Utech, F. H., & Kawano, S. (1975). Spectral polymorphisms in angiosperm flowers determined by differential ultraviolet reflectance. The Botanical Magazine = Shokubutsu-Gaku-Zasshi, 88(1), 9–30 https://doi.org/10.1007/BF02498877

93. van der Kooi, C. J., & Spaethe, J. (2022). Caution with colour calculations: spectral purity is a poor descriptor of flower colour visibility. Annals of Botany, 130(1), 1–9. https://doi.org/10.1093/AOB/MCAC069

94. van der Kooi, C. J., Stavenga, D. G., Arikawa, K., Belušič, G., & Kelber, A. (2021). Evolution of insect color vision: From spectral sensitivity to visual ecology. Annual Review of Entomology, 66(66), 435–461. https://doi.org/10.1146/annurev-ento-061720-071644

95. van der Kooi, C. J., Dyer, A. G., Kevan, P. G., & Lunau, K. (2019). Functional significance of the optical properties of flowers for visual signalling. Annals of Botany, 123(2), 263–276. https://doi.org/10.1093/AOB/M CY119

96. van der Kooi, C. J., Elzenga, J. T. M., Staal, M., & Stavenga, D. G. (2016). How to colour a flower: on the optical principles of flower coloration. Proceedings of the Royal Society B: Biological Sciences, 283(1830). https://doi.org/10.1098/RSPB.2016.0429

97. Vilà, M., Espinar, J. L., Hejda, M., Hulme, P. E., Jarošík, V., Maron, J. L., Pergl, J., Schaffner, U., Sun, Y., & Pyšek, P. (2011). Ecological impacts of invasive alien plants: A meta-analysis of their effects on species, communities and ecosystems. Ecology Letters, 14(7), 702–708. https://doi.org/10.1111/J.1461-0248.2011.01628.X

98. Vilà, M., Bartomeus, I., Dietzsch, A. C., Petanidou, T., Steffan-Dewenter, I., Stout, J. C., & Tscheulin, T. (2009). Invasive plant integration into native plant–pollinator networks across Europe. Proceedings of the Royal Society B: Biological Sciences, 276(1674), 3887. https://doi.org/10.1098/RSPB.2009.1076

99. Weiss, M. R., & Lamont, B. B. (2013). Floral color change and insect pollination: A dynamic relationship. Brill, 45(2–3), 185–199. https://doi.org/10.1080/07929978.1997.10676683

100. Weiss, M. R. (1995). Floral color change: A widespread functional convergence. American Journal of Botany, 82(2), 167–185. https://doi.org/10.2307/2445525

101. Weiss, M. R. (1991). Floral colour changes as cues for pollinators. Nature, 354(6350), 227–229. https://doi.org/10.1038/354227a0

102. Willmer, P., Stanley, D. A., Steijven, K., Matthews, I. M., & Nuttman, C. V. (2009). Bidirectional flower color and shape changes allow a second opportunity for pollination. Current Biology, 19(11), 919–923. https://doi.org/10.1016/J.CUB.2009.03.070

103. Wojcik, V. (2021). Pollinators: Their evolution, ecology, management, and conservation. in (ed.), arthropods - are they beneficial for mankind?. In Ranz, R. E. R (ed). Arthropods. IntechOpen. https://doi.org/10.5772/intechopen.97153

104. Yan, J., Wang, G., Sui, Y., Wang, M., & Zhang, L. (2016). Pollinator responses to floral colour change, nectar and scent promote reproductive fitness in Quisqualis indica (Combretaceae). Scientific Reports, 6(1), 1–10. https://doi.org/10.1038/srep24408

105. Young, P. A. V, Magure, P. K. H., Evans, J. R., Lafolley, N., Mooney, W. D., Andrews, M. C., Ginzburg, A., Peters, D. A., & Hamilton, R. M. (1991). Floral colour changes as cues for pollinators. Nature, 354(6350), 227–229. https://doi.org/10.1038/354227a0

