## Supplementary Data for "VARIATIONS IN THE ULTRAVIOLET FLORAL PATTERNS AND POLLINATOR PREFERENCE AMONG SELECTED NON-INVASIVE AND INVASIVE PLANTS OF TAMIL NADU, INDIA"

**Supplementary Table 1:** List of 188 plant species along with their respective invasive status, floral ultraviolet patterns, floral visible patterns, visible flower color group and pollinator category studied in parts of the Western Ghats and Eastern Ghats regions of Tamil Nadu, India.

| S. No. | Family and Species Name | Invasive Status | Ultraviolet Pattern | Visible Pattern | Visible Color Group | Pollinator |
|---|---|---|---|---|---|---|
| | **Acanthaceae** | | | | | |
| 1. | *Andrographis echioides* (L.) Nees | Non-invasive | UV-CC | VIS-CC | White Group | Melittophily & Psychophily |
| 2. | *Asystasia gangetica* (L.) T. Anderson | Invasive | UV-A | VIS-P | YOR Group | Melittophily & Psychophily |
| 3. | *Barleria cristata* L. | Invasive | UV-BE | VIS-P | VIB Group | Melittophily & Psychophily |
| 4. | *Barleria mysorensis* Heyne ex Roth | Non-invasive | UV-CR | VIS-P | VIB Group | Melittophily |
| 5. | *Blepharis maderaspatensis* (L.) Heyne ex Roth | Non-invasive | UV-CC | VIS-CC | White Group | Melittophily & Psychophily |
| 6. | *Justicia aurea* Schltdl. | Non-invasive | UV-A | VIS-CR | YOR Group | Ornithophily |
| 7. | *Justicia glauca* Rottl. | Non-invasive | UV-A | VIS-CC | VIB Group | Generalist |
| 8. | *Justicia scandens* Vahl | Non-invasive | UV-A | VIS-CC | White Group | Generalist |
| 9. | *Justicia tranquebariensis* Roxb. | Non-invasive | UV-A | VIS-CC | White Group | Psychophily |
| 10. | *Lepidagathis barberi* Gamble | Non-invasive | UV-A | VIS-CC | White Group | Generalist |
| 11. | *Ruellia caroliniensis* (J.F.Gmel.) Steud. | Non-invasive | UV-BE | VIS-P | VIB Group | Psychophily |
| 12. | *Ruellia patula* Jacq. | Non-invasive | UV-BE | VIS-P | VIB Group | Generalist |
| 13. | *Ruellia prostrata* Poir. | Non-invasive | UV-BE | VIS-P | VIB Group | Generalist |
| 14. | *Ruellia tuberosa* L. | Non-invasive | UV-BE | VIS-BE | VIB Group | Ornithophily |
| 15. | *Thunbergia coccinea* Wall. ex D.Don | Non-invasive | UV-CR | VIS-CR | YOR Group | Ornithophily |
| 16. | *Thunbergia laevis Wall. ex* Nees | Non-invasive | UV-BE | VIS-P | White Group | Generalist |
| | **Amaranthaceae** | | | | | |
| 17. | *Amaranthus spinosus* L. | Invasive | UV-CR | VIS-P | YOR Group | Anemophily |
| 18. | *Celosia argentea* L. | Non-invasive | UV-CR | VIS-CR | White Group | Melittophily & Psychophily |

| | | | | | | |
|---|---|---|---|---|---|---|
| 19. | *Celosia cristata* L. | Non-invasive | UV-A | VIS-P | YOR Group | Generalist |
| | **Apocynaceae** | | | | | |
| 20. | *Adenium obesum* (Forssk.) Roem. & Schult. | Non-invasive | UV-BE | VIS-CC | White Group | Generalist |
| 21. | *Asclepias curassavica* L. | Invasive | UV-A | VIS-CR | YOR Group | Melittophily & Psychophily |
| 22. | *Calotropis gigantea* (L.) W.T.Aiton | Invasive | UV-CR | VIS-P | VIB Group | Melittophily |
| 23. | *Catharanthus pusillus* (Murray) G.Don | Non-invasive | UV-A | VIS-P | White Group | Melittophily & Psychophily |
| 24. | *Catharanthus roseus* (L.) G.Don | Invasive | UV-BE | VIS-BE | VIB Group | Melittophily & Psychophily |
| 25. | *Ichnocarpus frutescens* (L.) W.T.Aiton | Non-invasive | UV-A | VIS-P | White Group | Melittophily |
| 26. | *Nerium oleander* L. | Invasive | UV-BE | VIS-P | VIB Group | Melittophily |
| 27. | *Plumeria alba* L. | Non-invasive | UV-BE | VIS-BE | White Group | Sphingophily |
| 28. | *Tabernaemontana divaricate* (L.) R.Br. ex Roem. & Schult. | Non-invasive | UV-A | VIS-P | White Group | Melittophily & Psychophily |
| 29. | *Vinca major* L. | Invasive | UV-BE | VIS-CC | VIB Group | Melittophily & Psychophily |
| | **Balsaminaceae** | | | | | |
| 30. | *Impatiens balsamina* L. | Invasive | UV-A | VIS-P | YOR Group | Melittophily & Psychophily |
| 31. | *Impatiens walleriana* Hook. f. | Invasive | UV-A | VIS-P | VIB Group | Melittophily & Psychophily |
| | **Bignoniaceae** | | | | | |
| 32. | *Tecoma stans* (L.) Juss. ex Kunth | Invasive | UV-BE | VIS-P | YOR Group | Melittophily |
| | **Boraginaceae** | | | | | |
| 33. | *Cynoglossum lanceolatum* Forssk. | Non-invasive | UV-BE | VIS-BE | White Group | Generalist |
| 34. | *Heliotropium marifolium* J.Koenig ex Retz. | Non-invasive | UV-A | VIS-BE | White Group | Psychophily |
| 35. | *Trichodesma indicum* (L.) Sm. | Non-invasive | UV-BE | VIS-BE | White Group | Melittophily |
| | **Brassicaceae** | | | | | |

| | | | | | | |
|---|---|---|---|---|---|---|
| 36. | *Brassica juncea* (L.) Czern. | Invasive | UV-BE | VIS-P | YOR Group | Autogamy |
| 37. | *Brassica rapa* L. | Invasive | UV-BE | VIS-P | YOR Group | Melittophily & Psychophily |
| | **Burseraceae** | | | | | |
| 38. | *Boswellia serrata* Roxb. | Non-invasive | UV-A | VIS-CR | White Group | Melittophily |
| | **Cactaceae** | | | | | |
| 39. | *Opuntia elata* Link & Otto ex Salm-Dyck | Non-invasive | UV-CC | VIS-CC | YOR Group | Generalist |
| | **Cannaceae** | | | | | |
| 40. | *Canna indica* L. | Invasive | UV-CC | VIS-CC | YOR Group | Ornithophily |
| 41. | *Canna paniculata* Ruiz & Pav. | Non-invasive | UV-A | VIS-CC | YOR Group | Ornithophily |
| | **Cleomaceae** | | | | | |
| 42. | *Cleome viscosa* L. | Invasive | UV-BE | VIS-P | YOR Group | Melittophily |
| | **Commelinaceae** | | | | | |
| 43. | *Commelina benghalensis* L. | Invasive | UV-A | VIS-CR | VIB Group | Autogamy |
| | **Compositae** | | | | | |
| 44. | *Ageratum conyzoides* L. | Invasive | UV-A | VIS-P | White Group | Generalist |
| 45. | *Bidens pilosa* L. | Invasive | UV-A | VIS-CR | White Group | Melittophily & Psychophily |
| 46. | *Cyanthillium cinereum* (L.) H.Rob. | Invasive | UV-A | VIS-CR | VIB Group | Anemophily |
| 47. | *Dahlia pinnata* Cav. | Non-invasive | UV-CR | VIS-CR | VIB Group | Melittophily |
| 48. | *Eclipta prostrata* (L.) L. | Invasive | UV-BE | VIS-P | White Group | Melittophily |
| 49. | *Erigeron karvinskianus* DC. | Invasive | UV-BE | VIS-BE | White Group | Melittophily |
| 50. | *Glebionis segetum* (L.) Fourr. | Non-invasive | UV-A | VIS-P | YOR Group | Melittophily & Psychophily |
| 51. | *Lipochaeta succulenta* (Hook. & Arn.) DC. | Non-invasive | UV-BE | VIS-P | YOR Group | Melittophily |
| 52. | *Parthenium hysterophorus* L. | Invasive | UV-A | VIS-P | White Group | Psychophily |

| | | | | | | |
|---|---|---|---|---|---|---|
| 53. | *Synedrella nodiflora* (L.) Gaertn. | Invasive | UV-CR | VIS-CR | YOR Group | Melittophily & Psychophily |
| 54. | *Tagetes erecta* L. | Invasive | UV-A | VIS-P | YOR Group | Melittophily & Psychophily |
| 55. | *Tagetes tenuifolia* Kunth | Invasive | UV-A | VIS-CC | YOR Group | Melittophily & Psychophily |
| 56. | *Tithonia diversifolia* (Hemsl.) A.Gray | Invasive | UV-BE | VIS-P | YOR Group | Melittophily & Psychophily |
| 57. | *Tridax procumbens* L. | Invasive | UV-A | VIS-CR | White Group | Melittophily & Psychophily |
| | **Convolvulaceae** | | | | | |
| 58. | *Cuscuta pentagona* Engelm. | Non-invasive | UV-A | VIS-P | White Group | Melittophily & Psychophily |
| 59. | *Evolvulus alsinoides* (L.) L. | Non-invasive | UV-CR | VIS-CR | VIB Group | Melittophily & Psychophily |
| 60. | *Ipomoea aquatica* Forssk. | Invasive | UV-BE | VIS-BE | White Group | Melittophily & Psychophily |
| 61. | *Ipomoea indica* (Burm.) Merr. | Invasive | UV-BE | VIS-BE | VIB Group | Melittophily |
| 62. | *Ipomoea littoralis* Blume | Non-invasive | UV-BE | VIS-BE | VIB Group | Melittophily |
| 63. | *Ipomoea obscura* (L.) Ker Gawl. | Invasive | UV-BE | VIS-BE | White Group | Generalist |
| 64. | *Ipomoea triloba* L. | Invasive | UV-A | VIS-BE | VIB Group | Melittophily |
| | **Crassulaceae** | | | | | |
| 65. | *Kalanchoe delagoensis* Eckl. & Zeyh. | Invasive | UV-P | VIS-CC | YOR Group | Generalist |
| | **Cucurbitaceae** | | | | | |
| 66. | *Citrullus lanatus* (Thunb.) Matsum. & Nakai | Non-invasive | UV-BE | VIS-P | YOR Group | Melittophily |
| 67. | *Coccinia grandis* (L.) Voigt | Invasive | UV-BE | VIS-CR | White Group | Melittophily |
| 68. | *Corallocarpus epigaeus* (Rottler) Hook.f. | Non-invasive | UV-A | VIS-P | YOR Group | Generalist |
| 69. | *Cucumis melo* L. | Non-invasive | UV-BE | VIS-P | YOR Group | Melittophily |
| 70. | *Momordica charantia* L. | Invasive | UV-CR | VIS-P | YOR Group | Melittophily |
| | **Cyperaceae** | | | | | |
| 71. | *Cyperus mindorensis* (Steud.) Huygh | Invasive | UV-A | VIS-P | White Group | Generalist |

| | | | | | | |
|---|---|---|---|---|---|---|
| 72. | *Kyllinga nemoralis* (J.R.Forst. & G.Forst.) Dandy ex Hutch. & Dalziel | Invasive | UV-A | VIS-P | White Group | Anemophily |
| | **Euphorbiaceae** | | | | | |
| 73. | *Croton bonplandianus* Baill. | Non-invasive | UV-A | VIS-P | White Group | Melittophily |
| 74. | *Euphorbia pulcherrima* Willd. ex Klotzsch | Non-invasive | UV-A | VIS-CR | YOR Group | Ornithophily |
| 75. | *Euphorbia rosea* Retz. | Non-invasive | UV-A | VIS-CR | White Group | Generalist |
| 76. | *Jatropha glandulifera* Roxb. | Non-invasive | UV-CC | VIS-P | YOR Group | Melittophily |
| 77. | *Jatropha gossypiifolia* L. | Invasive | UV-CC | VIS-BE | YOR Group | Melittophily & Psychophily |
| | **Gesneriaceae** | | | | | |
| 78. | *Henckelia humboldtiana* (Gardner) A.Weber & B.L.Burtt | Non-invasive | UV-A | VIS-P | VIB Group | Generalist |
| | **Hydrangeaceae** | | | | | |
| 79. | *Hydrangea macrophylla* (Thunb.) Ser. | Non-invasive | UV-P | VIS-CR | White Group | Melittophily & Psychophily |
| | **Lamiaceae** | | | | | |
| 80. | *Anisomeles malabarica* (L.) R.Br. | Non-invasive | UV-CC | VIS-CC | VIB Group | Generalist |
| 81. | *Clerodendrum infortunatum* L. | Non-invasive | UV-A | VIS-CC | White Group | Melittophily & Psychophily |
| 82. | *Coleus barbatus* (Andrews) Benth. ex G.Don | Non-invasive | UV-CR | VIS-CR | VIB Group | Generalist |
| 83. | *Endostemon viscosus* (Roth) M.R.Ashby | Non-invasive | UV-CC | VIS-P | White Group | Generalist |
| 84. | *Gmelina asiatica* L. | Non-invasive | UV-CC | VIS-P | YOR Group | Melittophily |
| 85. | *Hyptis suaveolens* (L.) Poit. | Invasive | UV-CC | VIS-CC | VIB Group | Melittophily |
| 86. | *Leucas aspera* (Willd.) Link | Non-invasive | UV-P | VIS-P | White Group | Psychophily |
| 87. | *Leucas biflora* (Vahl) Sm | Non-invasive | UV-P | VIS-P | White Group | Melittophily |
| 88. | *Ocimum tenuiflorum* L. | Invasive | UV-A | VIS-CR | White Group | Melittophily & Psychophily |
| 89. | *Orthosiphon thymiflorus* (Roth) Sleesen | Non-invasive | UV-P | VIS-P | White Group | Melittophily |
| 90. | *Salvia coccinea* Buc'hoz ex Etl. | Invasive | UV-CC | VIS-CC | YOR Group | Ornithophily |

| | | | | | | |
|---|---|---|---|---|---|---|
| 91. | *Salvia leucantha* Cav. | Invasive | UV-P | VIS-CC | White Group | Ornithophily |
| 92. | *Salvia splendens* Sellow ex Nees | Non-invasive | UV-P | VIS-P | YOR Group | Ornithophily |
| 93. | *Tectona grandis* L.f. | Invasive | UV-CR | VIS-CR | White Group | Melittophily |
| | **Leguminosae** | | | | | |
| 94. | *Abrus precatorius* L. | Invasive | UV-A | VIS-CC | VIB Group | Melittophily |
| 95. | *Acacia farnesiana* (L.) Willd. | Invasive | UV-A | VIS-P | YOR Group | Melittophily & Psychophily |
| 96. | *Acacia leucophloea* (Roxb.) Willd. | Non-invasive | UV-A | VIS-P | White Group | Melittophily |
| 97. | *Acacia nilotica* (L.) Willd. ex Delile | Invasive | UV-A | VIS-P | YOR Group | Melittophily & Psychophily |
| 98. | *Canavalia gladiate* (Jacq.) DC. | Invasive | UV-CC | VIS-CC | VIB Group | Melittophily |
| 99. | *Cassia mimosoides* L. | Non-invasive | UV-CR | VIS-P | YOR Group | Generalist |
| 100. | *Chamaecrista pumila* (Lam.) V.Singh | Non-invasive | UV-CR | VIS-P | YOR Group | Melittophily |
| 101. | *Clitoria ternatea* L. | Invasive | UV-CC | VIS-CC | VIB Group | Melittophily & Psychophily |
| 102. | *Crotalaria grahamiana* Wight & Arn. | Non-invasive | UV-CC | VIS-P | YOR Group | Melittophily |
| 103. | *Crotalaria pallida* Aiton | Non-invasive | UV-CC | VIS-CC | YOR Group | Melittophily |
| 104. | *Crotalaria paniculata* Willd. | Non-invasive | UV-CC | VIS-P | YOR Group | Melittophily |
| 105. | *Crotalaria verrucosa* L. | Non-invasive | UV-CC | VIS-CC | White Group | Melittophily |
| 106. | *Desmanthus virgatus* (L.) Willd. | Non-invasive | UV-CR | VIS-CR | White Group | Psychophily |
| 107. | *Erythrina variegate* L. | Non-invasive | UV-CR | VIS-CC | YOR Group | Ornithophily |
| 108. | *Indigofera aspalathoides* Vahl ex DC. | Non-invasive | UV-CC | VIS-CC | YOR Group | Generalist |
| 109. | *Indigofera tinctoria* Forssk. | Invasive | UV-CC | VIS-CC | YOR Group | Generalist |
| 110. | *Macroptilium atropurpureum* (DC.) Urb. | Invasive | UV-A | VIS-CR | YOR Group | Autogamy |
| 111. | *Mimosa pudica* L. | Invasive | UV-CR | VIS-P | VIB Group | Melittophily |
| 112. | *Peltophorum pterocarpum* (DC.) Backer ex K.Heyne | Invasive | UV-CR | VIS-P | YOR Group | Melittophily |

| | | | | | | |
|---|---|---|---|---|---|---|
| 113. | *Prosopis juliflora* (Sw.) DC. | Invasive | UV-A | VIS-CR | YOR Group | Melittophily |
| 114. | *Pseudarthria viscida* (L.) Wight & Arn. | Non-invasive | UV-CC | VIS-CC | VIB Group | Generalist |
| 115. | *Senna auriculata* (L.) Roxb. | Non-invasive | UV-CR | VIS-CR | YOR Group | Melittophily |
| 116. | *Senna occidentalis* (L.) Link | Invasive | UV-CR | VIS-CR | YOR Group | Melittophily |
| 117. | *Senna septemtrionalis* (Viv.) H.S.Irwin & Barneby | Invasive | UV-CR | VIS-CR | YOR Group | Melittophily |
| 118. | *Senna siamea* (Lam.) H.S.Irwin & Barneby | Invasive | UV-CR | VIS-CR | YOR Group | Melittophily |
| 119. | *Stylosanthes mucronata* Willd. | Non-invasive | UV-CC | VIS-CC | YOR Group | Melittophily & Psychophily |
| 120. | *Tamarindus indica* L. | Invasive | UV-A | VIS-CC | YOR Group | Melittophily |
| 121. | *Tephrosia purpurea* (L.) Pers. | Non-invasive | UV-A | VIS-CC | VIB Group | Melittophily |
| | **Lophiocarpaceae** | | | | | |
| 122. | *Corbichonia decumbens* (Forssk.) Exell | Non-invasive | UV-BE | VIS-CR | VIB Group | Melittophily |
| | **Loranthaceae** | | | | | |
| 123. | *Dendrophthoe falcata* (L.f.) Ettingsh. | Non-invasive | UV-A | VIS-CR | YOR Group | Ornithophily |
| | **Lythraceae** | | | | | |
| 124. | *Cuphea hyssopifolia* Griseb. | Invasive | UV-BE | VIS-P | VIB Group | Melittophily |
| 125. | *Lagerstroemia speciosa* (L.) Pers. | Invasive | UV-CR | VIS-CR | VIB Group | Melittophily |
| 126. | *Punica granatum* L. | Non-invasive | UV-CR | VIS-CR | YOR Group | Melittophily & Psychophily |
| | **Malvaceae** | | | | | |
| 127. | *Hibiscus calyphyllus* Cav. | Non-invasive | UV-A | VIS-P | White Group | Generalist |
| 128. | *Hibiscus micranthus* L.f. | Non-invasive | UV-A | VIS-P | White Group | Generalist |
| 129. | *Hibiscus rosa-sinensis* L. | Non-invasive | UV-BE | VIS-BE | YOR Group | Generalist |
| 130. | *Hibiscus solandra* L'Hér. | Non-invasive | UV-A | VIS-P | White Group | Generalist |
| 131. | *Hibiscus vitifolius* Mill. | Non-invasive | UV-A | VIS-BE | White Group | Generalist |

| | | | | | | |
|---|---|---|---|---|---|---|
| 132. | *Malvaviscus arboreus var. arboreus* Cav. | Invasive | UV-A | VIS-P | YOR Group | Ornithophily |
| 133. | *Sida acuta* Burm.f. | Invasive | UV-BE | VIS-P | YOR Group | Melittophily & Psychophily |
| 134. | *Sida cordata* (Burm.f.) Borss.Waalk. | Non-invasive | UV-BE | VIS-P | YOR Group | Melittophily & Psychophily |
| 135. | *Sida cordifolia* L. | Non-invasive | UV-BE | VIS-CR | White Group | Melittophily & Psychophily |
| 136. | *Sida ovata* Forssk. | Non-invasive | UV-BE | VIS-P | YOR Group | Melittophily |
| 137. | *Sida rhombifolia* L. | Non-invasive | UV-BE | VIS-P | YOR Group | Melittophily |
| 138. | *Waltheria indica* L. | Invasive | UV-BE | VIS-P | YOR Group | Melittophily & Psychophily |
| | **Meliaceae** | | | | | |
| 139. | *Melia azedarach* L. | Invasive | UV-CR | VIS-CR | White Group | Melittophily & Psychophily |
| | **Nyctaginaceae** | | | | | |
| 140. | *Boerhavia coccinea* Mill. | Invasive | UV-A | VIS-P | VIB Group | Autogamy |
| 141. | *Boerhavia erecta* L. | Non-invasive | UV-A | VIS-P | White Group | Generalist |
| 142. | *Bougainvillea glabra* Choisy | Non-invasive | UV-A | VIS-CC | VIB Group | Melittophily & Psychophily |
| 143. | *Bougainvillea spectabilis* Willd. | Invasive | UV-A | VIS-CC | White Group | Melittophily & Psychophily |
| 144. | *Mirabilis jalapa* L. | Invasive | UV-CR | VIS-P | VIB Group | Phalaenophily |
| | **Oleaceae** | | | | | |
| 145. | *Jasminum griffithianum* L. | Non-invasive | UV-A | VIS-P | White Group | Melittophily |
| 146. | *Jasminum mesnyi* Hance | Invasive | UV-BE | VIS-P | YOR Group | Melittophily |
| 147. | *Jasminum multiflorum* (Burm.f.) Andrews | Invasive | UV-P | VIS-P | White Group | Phalaenophily |
| | **Onagraceae** | | | | | |
| 148. | *Fuchsia magellanica* Lam. | Non-invasive | UV-CC | VIS-CC | YOR Group | Ornithophily |
| | **Oxalidaceae** | | | | | |
| 149. | *Oxalis corniculata* L. | Invasive | UV-BE | VIS-P | YOR Group | Autogamy |

| | | | | | | |
|---|---|---|---|---|---|---|
| | **Passifloraceae** | | | | | |
| 150. | *Passiflora tarminiana* Coppens & V.E.Barney | Invasive | UV-A | VIS-CR | VIB Group | Ornithophily |
| 151. | *Passiflora vitifolia* Kunth | Non-invasive | UV-CR | VIS-CR | YOR Group | Melittophily |
| | **Pedaliaceae** | | | | | |
| 152. | *Pedalium murex* L. | Non-invasive | UV-A | VIS-BE | White Group | Melittophily & Psychophily |
| 153. | *Sesamum radiatum* Thonn. ex Hornem. | Non-invasive | UV-CC | VIS-CC | White Group | Autogamy |
| | **Plumbaginaceae** | | | | | |
| 154. | *Plumbago zeylanica* L. | Non-invasive | UV-BE | VIS-CR | White Group | Melittophily |
| | **Polygalaceae** | | | | | |
| 155. | *Polygala arvensis* Willd. | Non-invasive | UV-CR | VIS-CC | YOR Group | Generalist |
| | **Portulacaceae** | | | | | |
| 156. | *Portulaca oleracea* L. | Invasive | UV-BE | VIS-P | YOR Group | Autogamy |
| | **Rhamnaceae** | | | | | |
| 157. | *Ziziphus nummularia* (Burm.f.) Wight & Arn. | Non-invasive | UV-A | VIS-P | YOR Group | Melittophily |
| 158. | *Ziziphus oenopolia* (L.) Mill. | Non-invasive | UV-A | VIS-P | YOR Group | Melittophily |
| | **Rosaceae** | | | | | |
| 159. | *Prunus cerasoides* Koidz. | Non-invasive | UV-CR | VIS-CR | VIB Group | Melittophily |
| | **Rubiaceae** | | | | | |
| 160. | *Catunaregam spinosa* (Thunb.) Tirveng. | Non-invasive | UV-A | VIS-P | White Group | Melittophily |
| 161. | *Ixora chinensis* Lam | Non-invasive | UV-CR | VIS-CR | VIB Group | Melittophily & Psychophily |
| 162. | *Ixora coccinea* Curtis | Non-invasive | UV-CR | VIS-P | YOR Group | Melittophily & Psychophily |
| 163. | *Knoxia wightiana* Wall. ex Wight & Arn | Non-invasive | UV-A | VIS-CC | VIB Group | Generalist |
| 164. | *Mitracarpus hirtus* (L.) DC | Non-invasive | UV-A | VIS-P | White Group | Generalist |

| | | | | | | |
|---|---|---|---|---|---|---|
| 165. | *Morinda coreia* Buch. -Ham. | Non-invasive | UV-A | VIS-P | White Group | Generalist |
| 166. | *Spermacoce hispida* L. | Non-invasive | UV-A | VIS-P | White Group | Melittophily & Psychophily |
| 167. | *Spermacoce remota* Lam. | Non-invasive | UV-A | VIS-P | White Group | Melittophily & Psychophily |
| | **Rutaceae** | | | | | |
| 168. | *Ruta graveolens* L. | Non-invasive | UV-A | VIS-P | YOR Group | Melittophily & Psychophily |
| 169. | *Toddalia asiatica* (L.) Lam. | Non-invasive | UV-A | VIS-P | White Group | Melittophily |
| | **Sapindaceae** | | | | | |
| 170. | *Cardiospermum canescens* Wall. | Non-invasive | UV-A | VIS-CR | White Group | Melittophily & Psychophily |
| 171. | *Cardiospermum corindum* L. | Non-invasive | UV-A | VIS-CR | White Group | Melittophily |
| | **Solanaceae** | | | | | |
| 172. | *Brugmansia suaveolens* (Humb. & Bonpl. ex Willd.) Sweet | Invasive | UV-P | VIS-P | VIB Group | Chiropterophily |
| 173. | *Capsicum frutescens* L. | Non-invasive | UV-CR | VIS-P | YOR Group | Melittophily |
| 174. | *Datura innoxia* Mill. | Invasive | UV-BE | VIS-P | White Group | Melittophily & Psychophily |
| 175. | *Solanum americanum* Mill. | Invasive | UV-CR | VIS-CR | White Group | Melittophily & Psychophily |
| 176. | *Solanum crispum* Ruiz & Pav. | Non-invasive | UV-CR | VIS-CR | VIB Group | Melittophily |
| 177. | *Solanum erianthum* D.Don | Invasive | UV-CR | VIS-CR | White Group | Melittophily |
| 178. | *Solanum melongena* L. | Non-invasive | UV-CR | VIS-CR | VIB Group | Melittophily |
| 179. | *Solanum melongena var. insanum* (L.) Filov | Non-invasive | UV-CR | VIS-CR | VIB Group | Melittophily |
| 180. | *Solanum torvum* Buch.-Ham. ex Wall. | Invasive | UV-CR | VIS-CR | White Group | Generalist |
| | **Tropaeolaceae** | | | | | |
| 181. | *Tropaeolum majus* L. | Invasive | UV-CR | VIS-CC | YOR Group | Melittophily |
| | **Verbenaceae** | | | | | |
| 182. | *Lantana camara* L. | Invasive | UV-BE | VIS-BE | VIB Group | Melittophily & Psychophily |

| | | | | | | |
|---|---|---|---|---|---|---|
| 183. | *Lantana veronicifolia* Hayek | Non-invasive | UV-BE | VIS-BE | White Group | Generalist |
| 184. | *Priva cordifolia* (L.f.) Druce | Non-invasive | UV-BE | VIS-P | White Group | Generalist |
| 185. | *Stachytarpheta cayennensis* (Rich.) Vahl | Invasive | UV-BE | VIS-P | VIB Group | Psychophily |
| 186. | *Stachytarpheta jamaicensis* (L.) Vahl | Invasive | UV-BE | VIS-P | VIB Group | Psychophily |
| | **Violaceae** | | | | | |
| 187. | *Hybanthus enneaspermus* (L.) F.Muell. | Non-invasive | UV-A | VIS-CC | VIB Group | Autogamy |
| | **Zygophyllaceae** | | | | | |
| 188. | *Tribulus terrestris* L. | Invasive | UV-BE | VIS-P | YOR Group | Melittophily & Psychophily |

**Notes:**

- UV-A – Ultraviolet Absorbing
- UV-P – Ultraviolet Patternless
- UV-BE – Ultraviolet Bullseye pattern
- UV-CC – Ultraviolet Contrasting Corolla pattern
- UV-CR – Ultraviolet Contrasting Reproductive Structures pattern
- VIS-P – Visible Patternless
- VIS-BE – Visible Bullseye pattern
- VIS-CC – Visible Contrasting Corolla pattern
- VIS-CR – Visible Contrasting Reproductive Structures pattern
- VIB Group – Violet/ Indigo/ Blue spectrum of colors
- YOR Group – Yellow/ Orange/ Red spectrum of colors
- White Group – White/ Cream/ Greenish-white/ Yellowish-white/ Pinkish-white colors

**Supplimentay Figure 1:** Visible and ultraviolet patterns observed among 188 plant species collected in parts of the Western Ghats and Eastern Ghats regions of Tamil Nadu. Visible spectral and ultraviolet spectral images are displayed in the first and second columns respectively. Each row displays images from one species and can identified with their respective row numbers in each of the following pages.

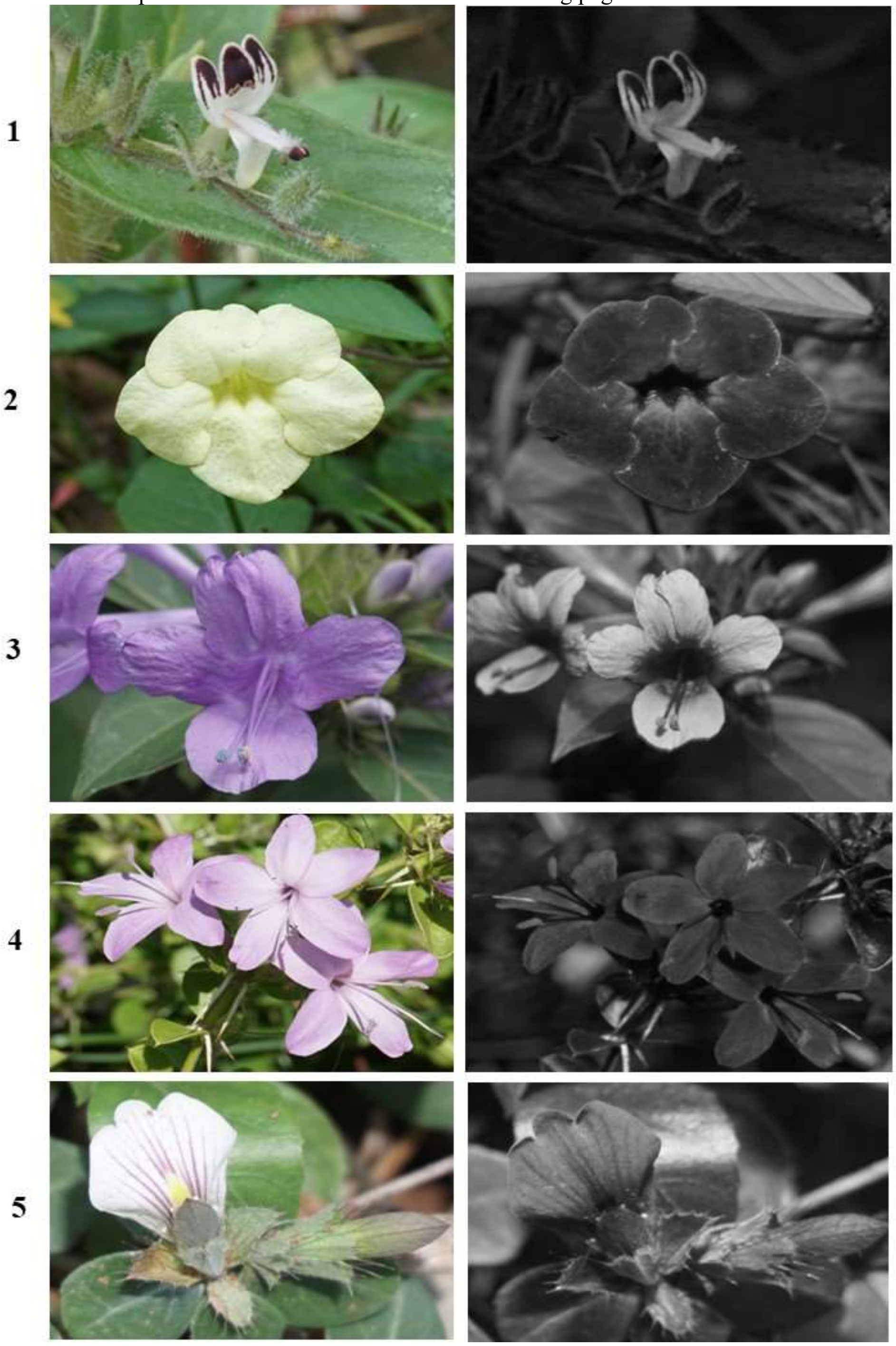


**Species list:** 1) *Andrographis echioides* (L.) Nees (Acanthaceae), 2) *Asystasia gangetica* (L.) T. Anderson (Acanthaceae), 3) *Barleria cristata* L. (Acanthaceae), 4) *Barleria mysorensis* Heyne ex Roth (Acanthaceae), and 5) *Blepharis maderaspatensis* (L.) Heyne ex Roth (Acanthaceae).

6
7
8
9
10

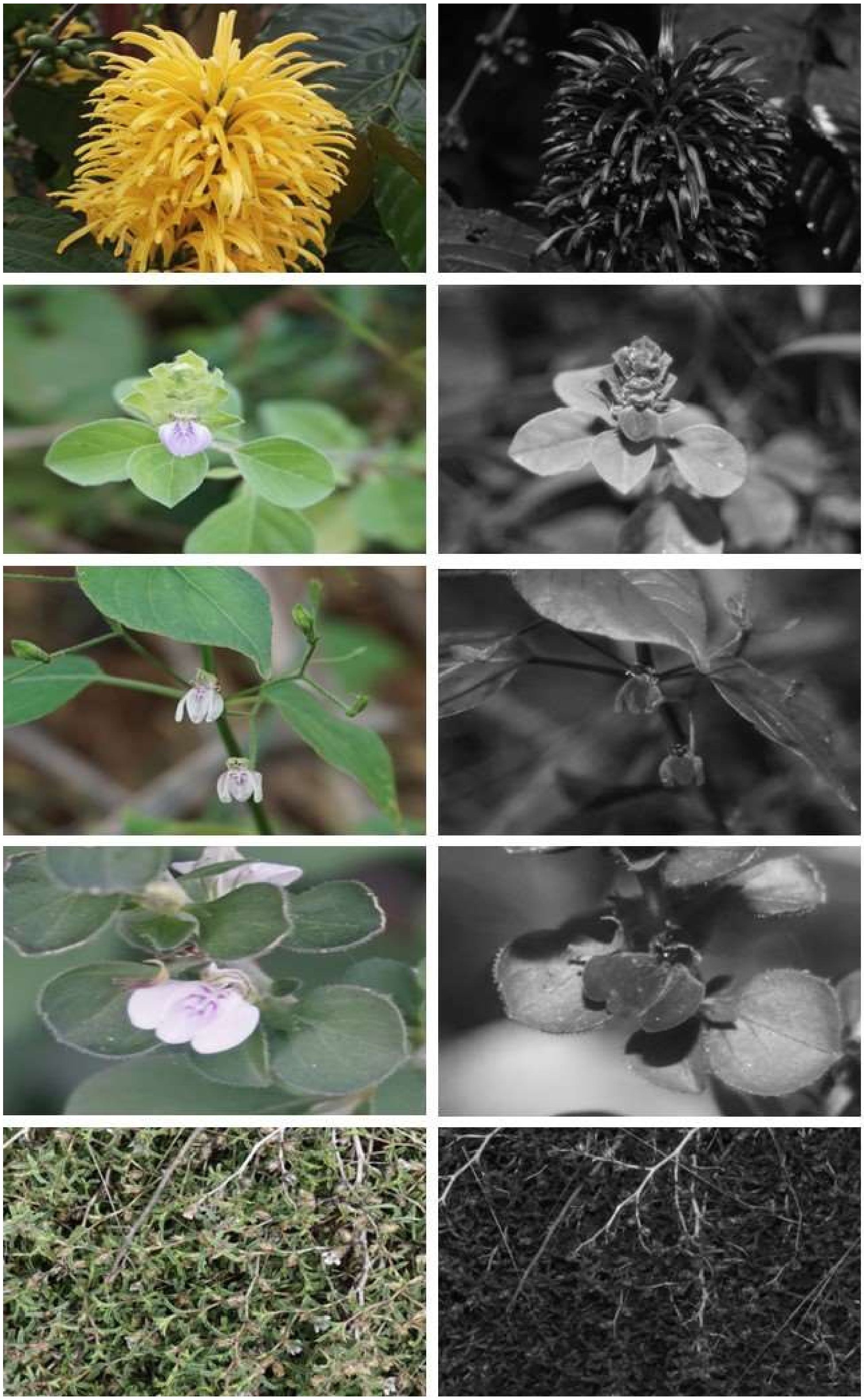

**Species list (cont.):** 6) *Justicia aurea* Schltdl. (Acanthaceae), 7) *Justicia glauca* Rottl. (Acanthaceae), 8) *Justicia scandens* Vahl (Acanthaceae), 9) *Justicia tranquebariensis* Roxb. (Acanthaceae), and 10) *Lepidagathis barberi* Gamble (Acanthaceae).

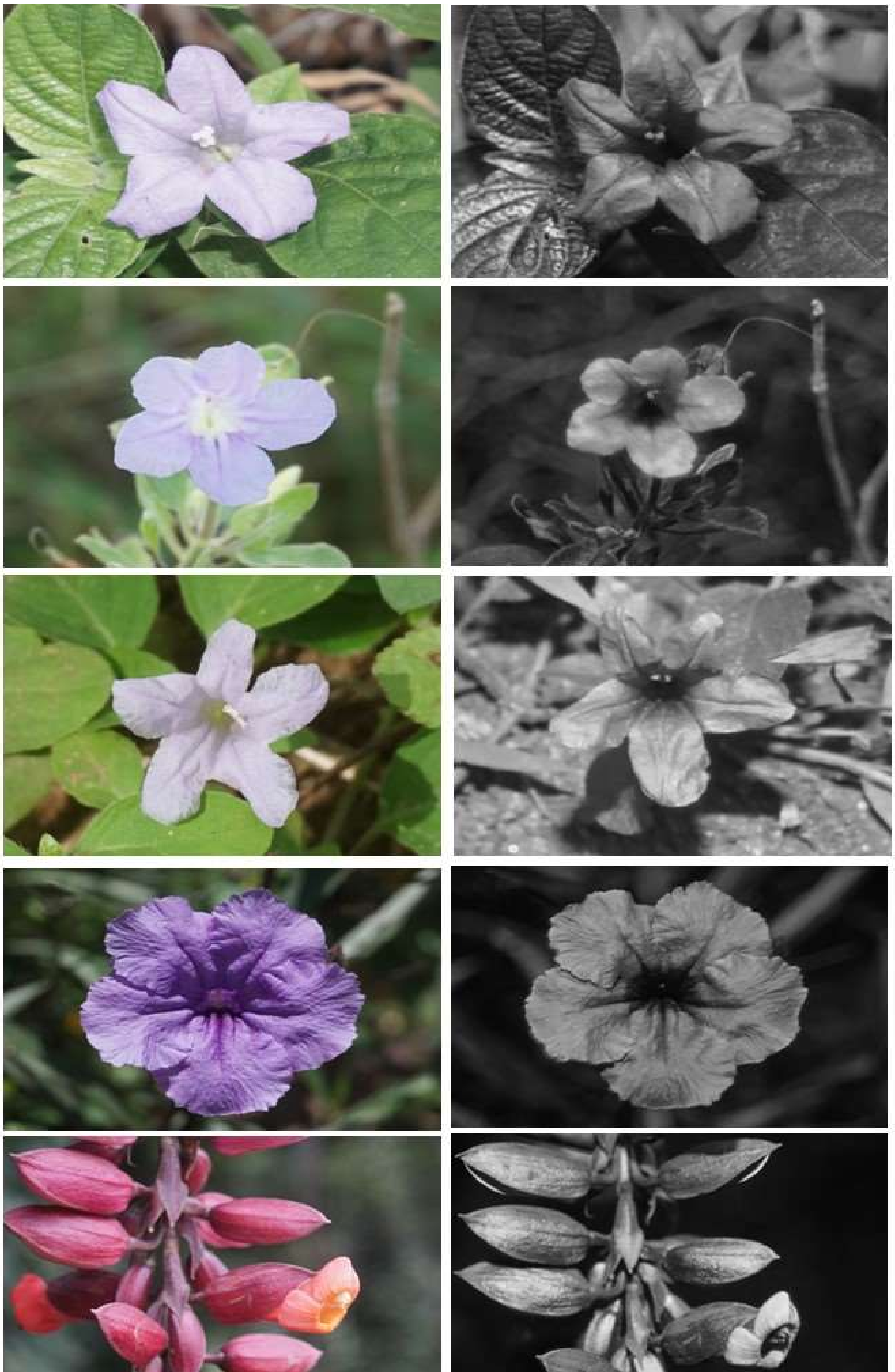

11

12

13

14

15

**Species list (cont.):** 11) *Ruellia caroliniensis* (J.F.Gmel.) Steud. (Acanthaceae), 12) *Ruellia patula* Jacq. (Acanthaceae), 13) *Ruellia prostrata* Poir. (Acanthaceae), 14) *Ruellia tuberosa* L. (Acanthaceae), and 15) *Thunbergia coccinea* Wall. ex D.Don (Acanthaceae).

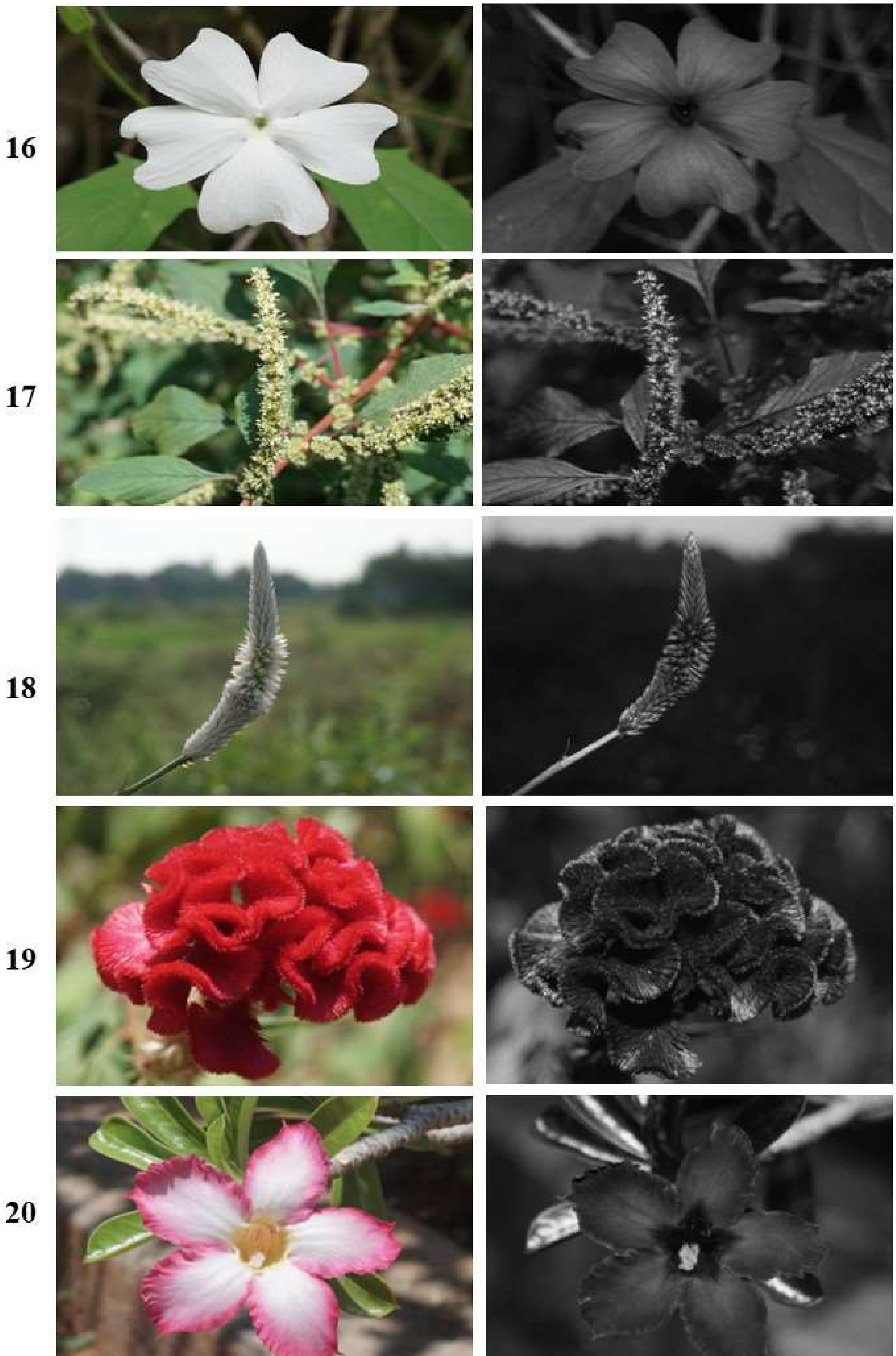


**Species list (cont.):** 16) *Thunbergia laevis* Wall. ex Nees (Acanthaceae), 17) *Amaranthus spinosus* L. (Amaranthaceae), 18) *Celosia argentea* L. (Amaranthaceae), 19) *Celosia cristata* L. (Amaranthaceae), and 20) *Adenium obesum* (Forssk.) Roem. & Schult. (Apocynaceae).

21

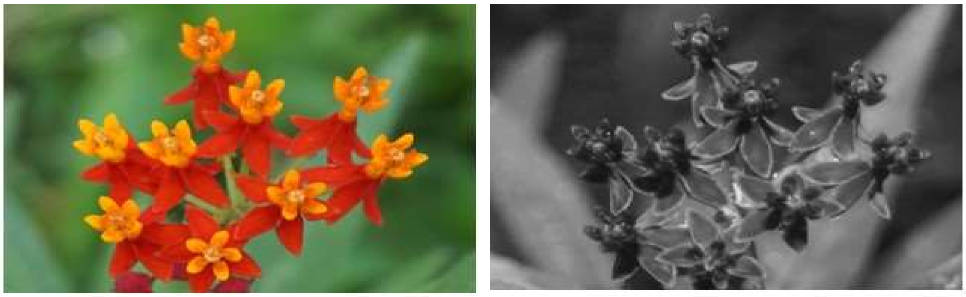

22

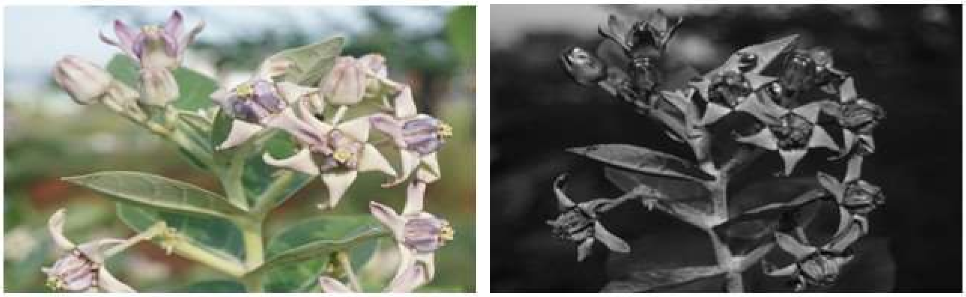

23

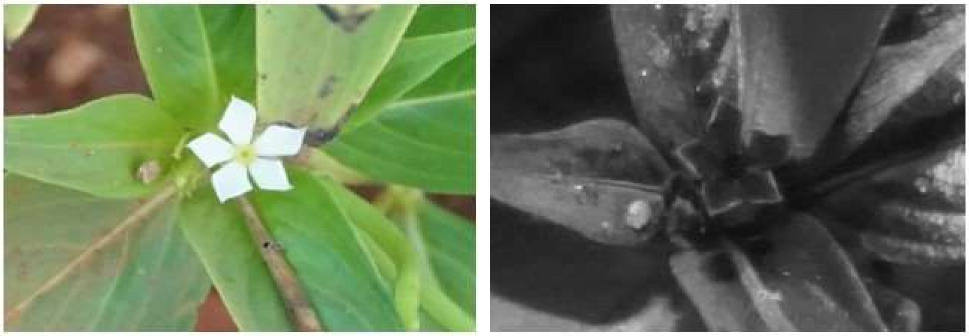

24

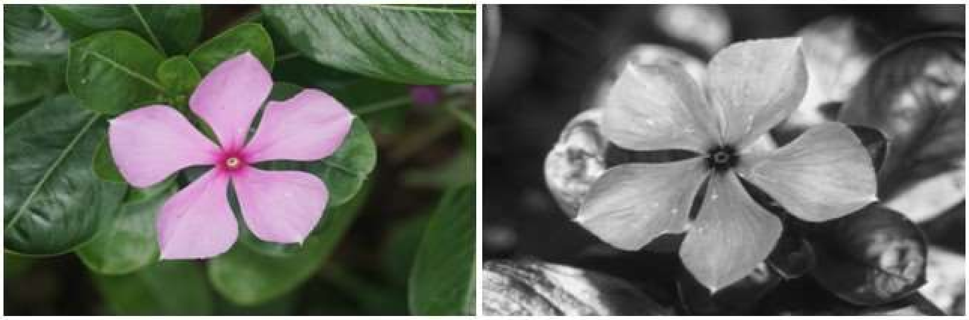

25

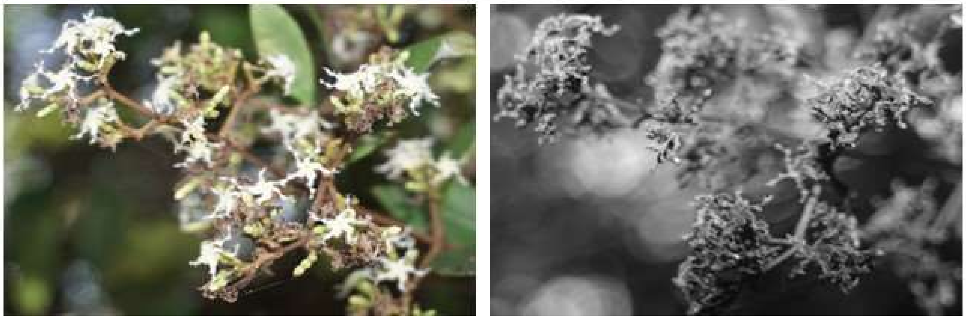

**Species list (cont.):** 21) *Asclepias curassavica* L. (Apocynaceae), 22) *Calotropis gigantea* (L.) W.T.Aiton (Apocynaceae), 23) *Catharanthus pusillus* (Murray) G.Don (Apocynaceae), 24) *Catharanthus roseus* (L.) G.Don (Apocynaceae), and 25) *Ichnocarpus frutescens* (L.) W.T.Aiton (Apocynaceae).

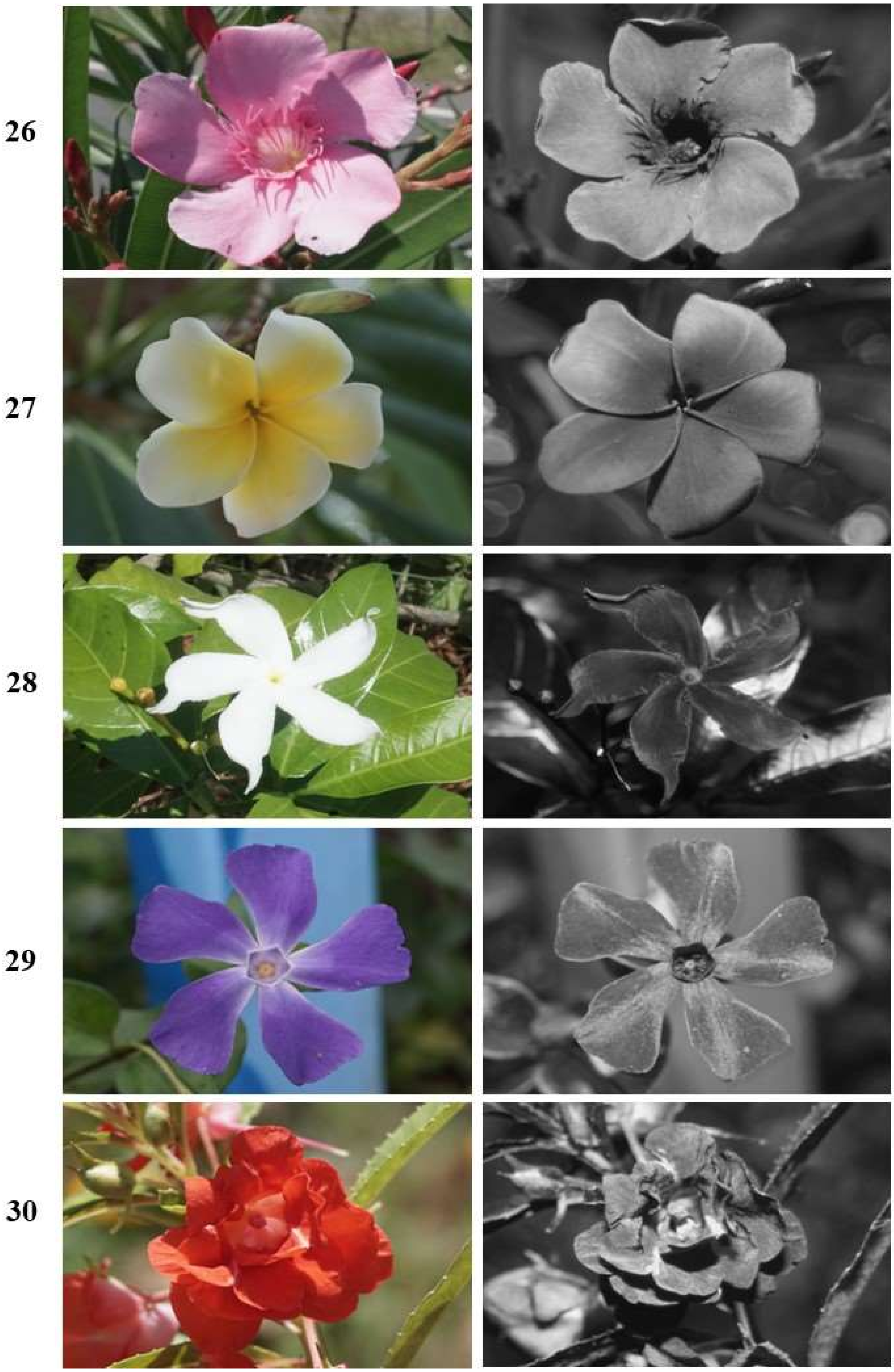


**Species list (cont.):** 26) *Nerium oleander* L. (Apocynaceae), 27) *Plumeria alba* L. (Apocynaceae), 28) *Tabernaemontana divaricate* (L.) R.Br. ex Roem. & Schult. (Apocynaceae), 29) *Vinca major* L. (Apocynaceae), and 30) *Impatiens balsamina* L. (Balsaminaceae).

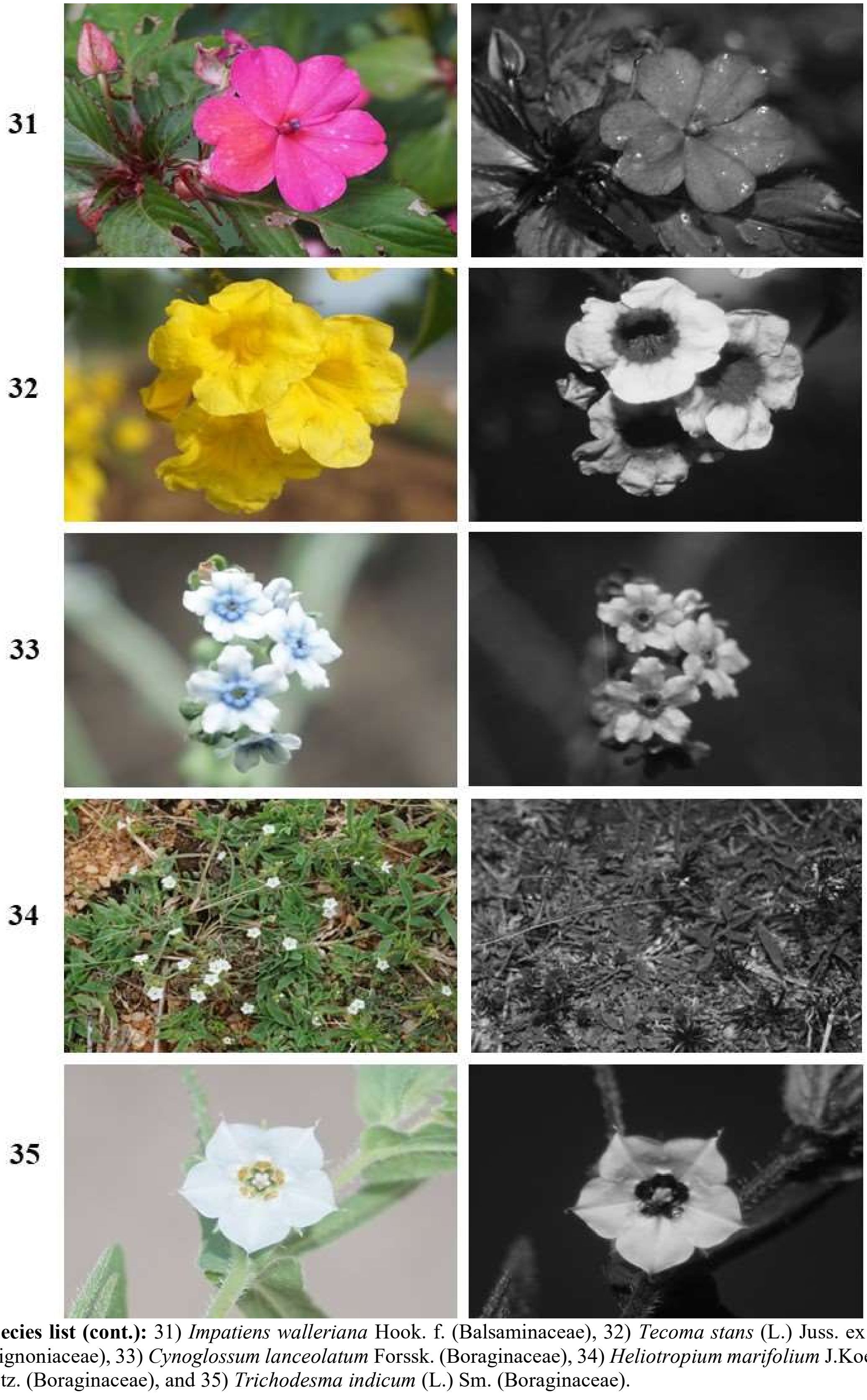


**Species list (cont.):** 31) *Impatiens walleriana* Hook. f. (Balsaminaceae), 32) *Tecoma stans* (L.) Juss. ex Kunth (Bignoniaceae), 33) *Cynoglossum lanceolatum* Forssk. (Boraginaceae), 34) *Heliotropium marifolium* J.Koenig ex Retz. (Boraginaceae), and 35) *Trichodesma indicum* (L.) Sm. (Boraginaceae).

36

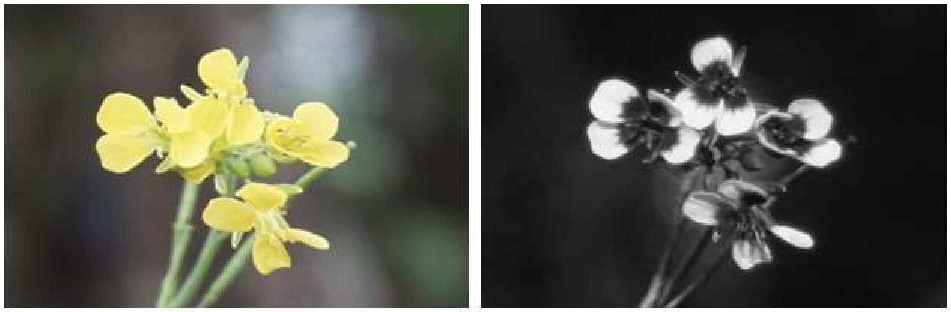

37

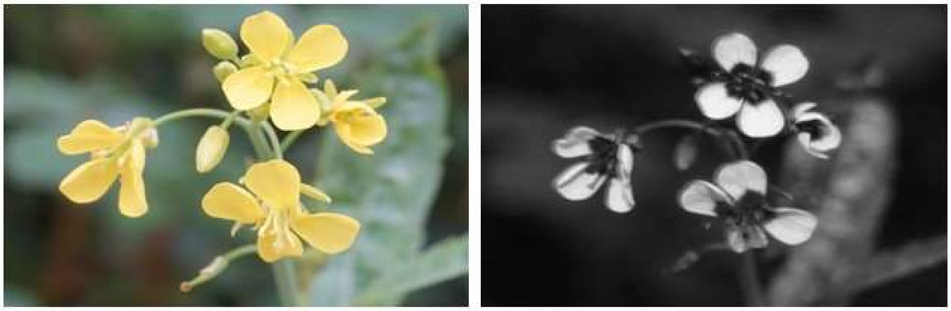

38

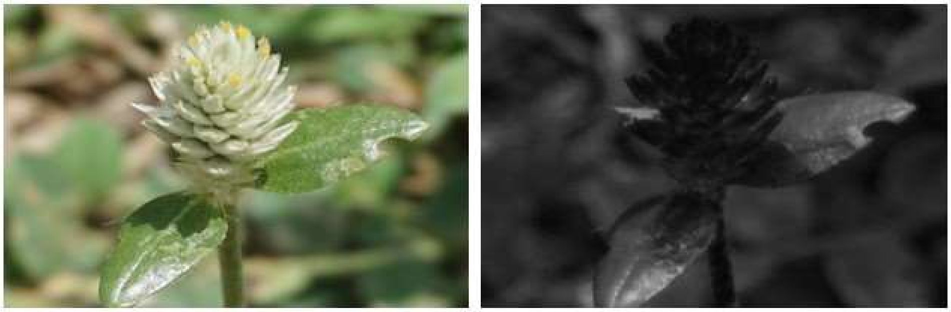

39

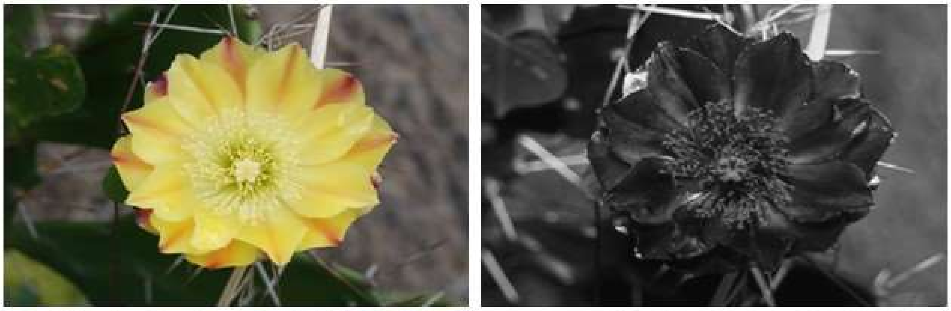

40

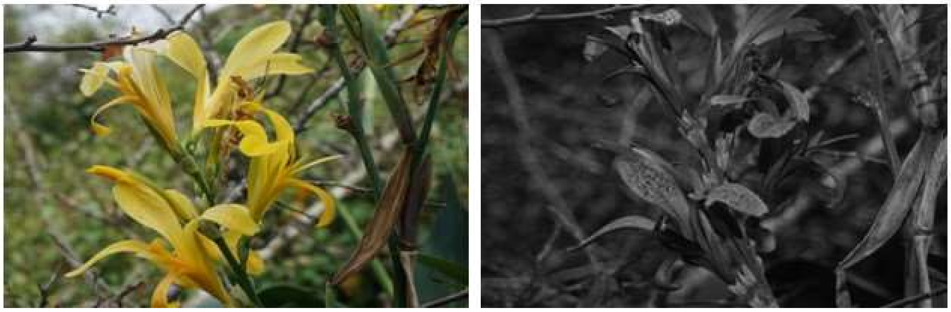

**Species list (cont.):** 36) *Brassica juncea* (L.) Czern. (Brassicaceae), 37) *Brassica rapa* L. (Brassicaceae), 38) *Boswellia serrata* Roxb. (Burseraceae), 39*) Opuntia elata* Link & Otto ex Salm-Dyck (Cactaceae), and 40) *Canna indica* L. (Cannaceae).

41 42 43 44 45

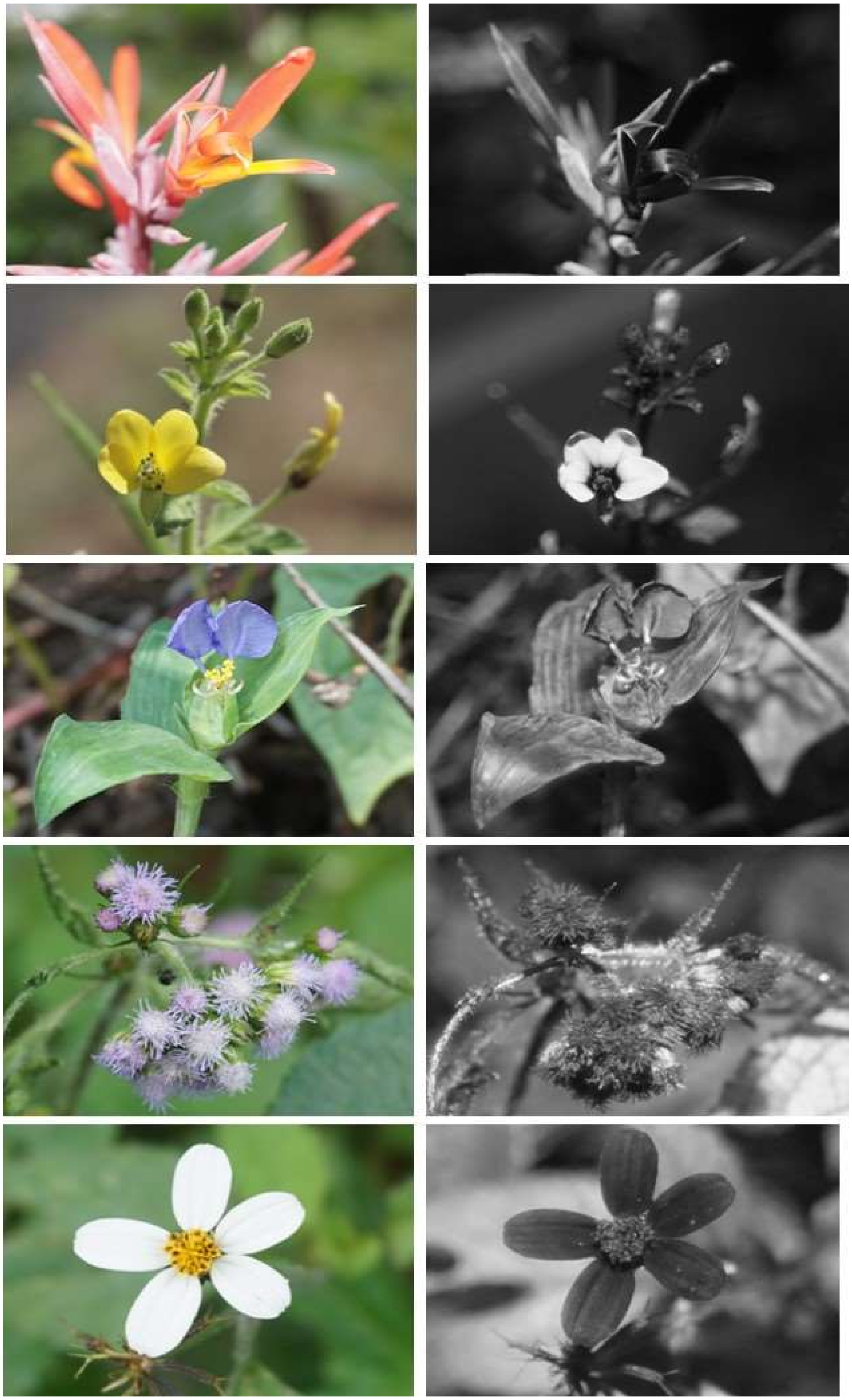

**Species list (cont.):** 41) *Canna paniculata* Ruiz & Pav. (Cannaceae), 42) *Cleome viscosa* L. (Cleomaceae), 43) *Commelina benghalensis* L. (Commelinaceae), 44) *Ageratum conyzoides* L. (Compositae), and 45) *Bidens pilosa* L. (Compositae).

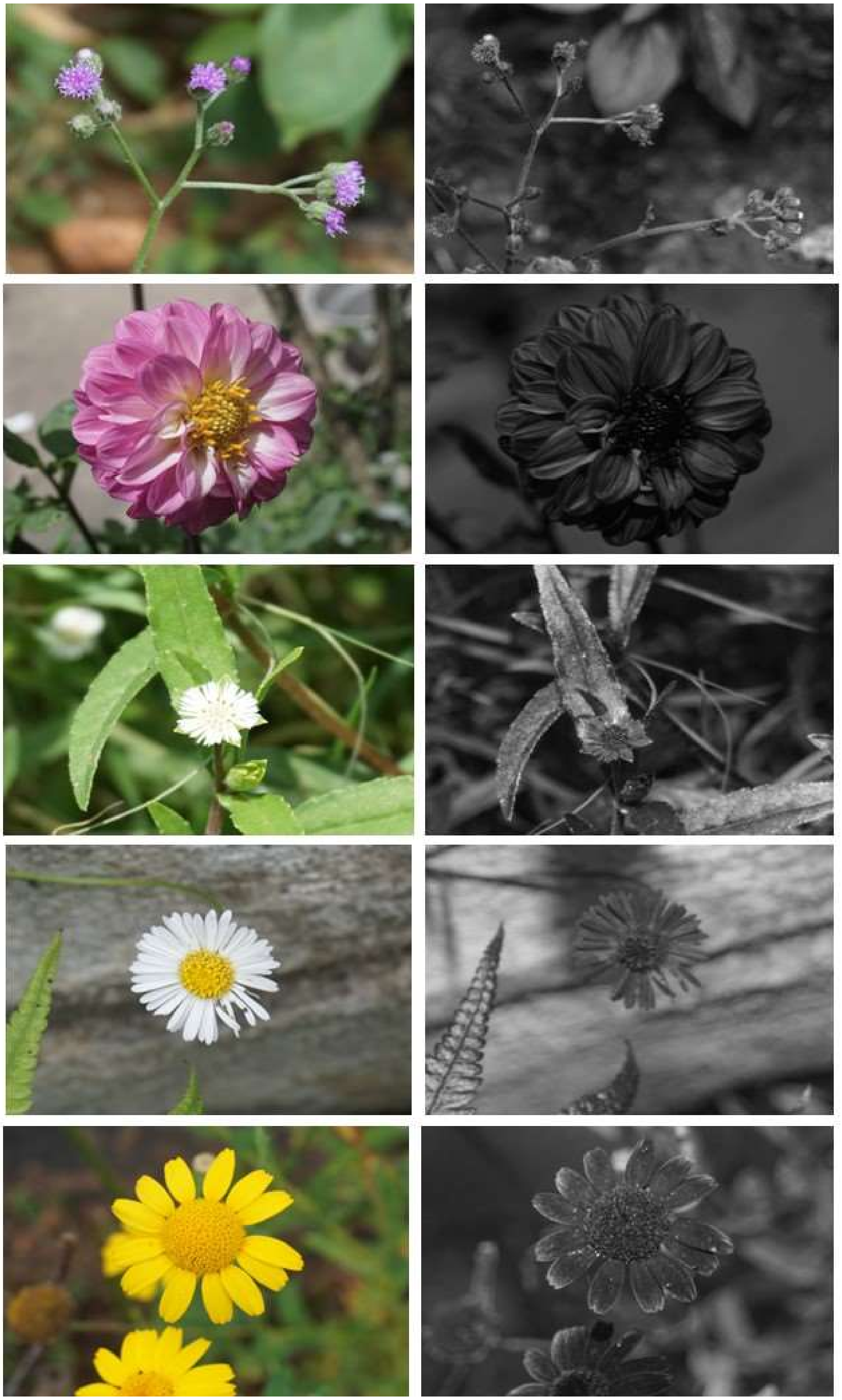
46

47

48

49

50

**Species list (cont.):** 46) *Cyanthillium cinereum* (L.) H.Rob. (Compositae), 47) *Dahlia pinnata* Cav. (Compositae), 48) *Eclipta prostrata* (L.) L. (Compositae), 49) *Erigeron karvinskianus* DC. (Compositae), and 50) *Glebionis segetum* (L.) Fourr. (Compositae).

51

52

53

54

55

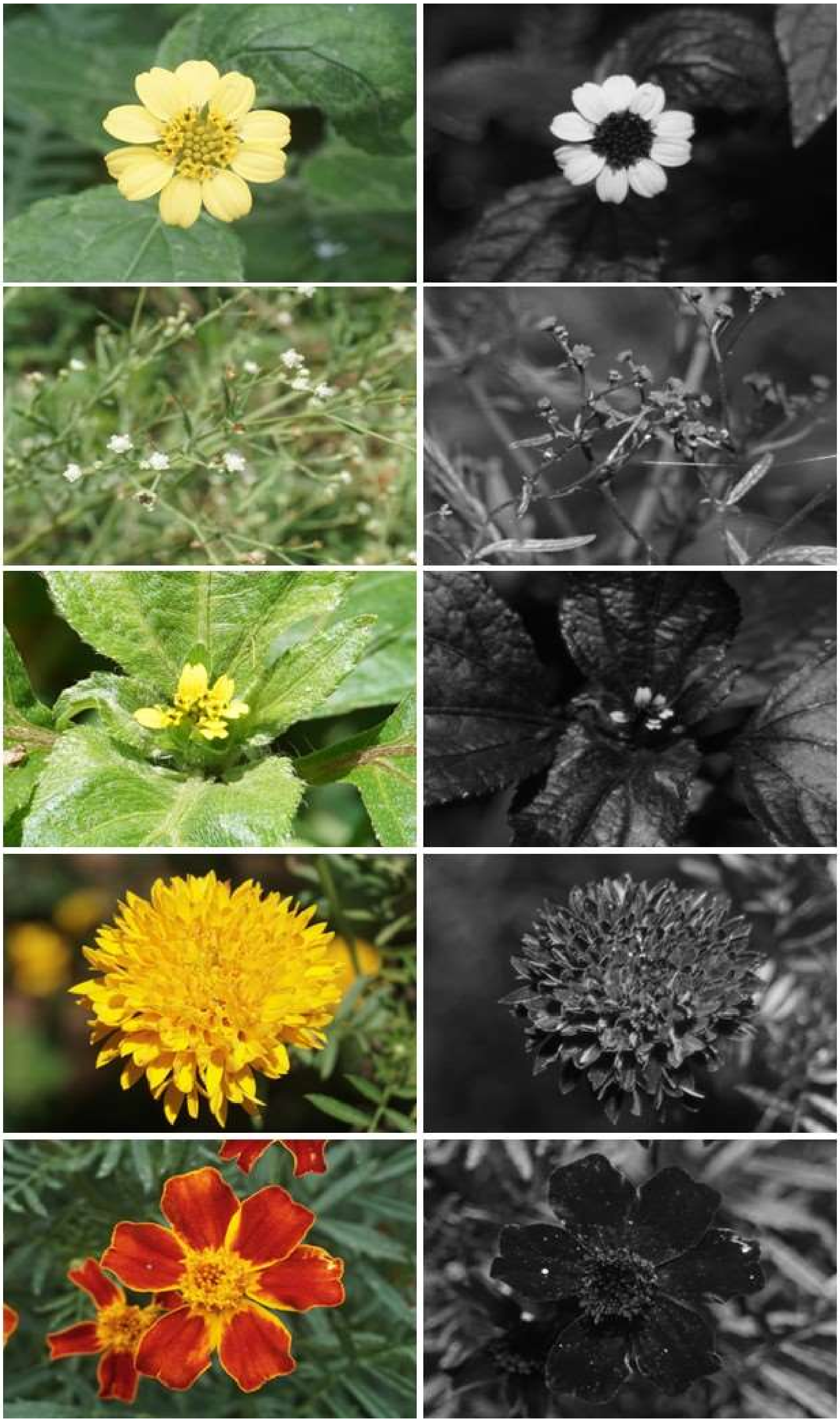

**Species list (cont.):** 51) *Lipochaeta succulenta* (Hook. & Arn.) DC. (Compositae), 52) P*arthenium hysterophorus* L. (Compositae), 53) *Synedrella nodiflora* (L.) Gaertn. (Compositae), 54) *Tagetes erecta* L. (Compositae), and 55) *Tagetes tenuifolia* Kunth (Compositae).

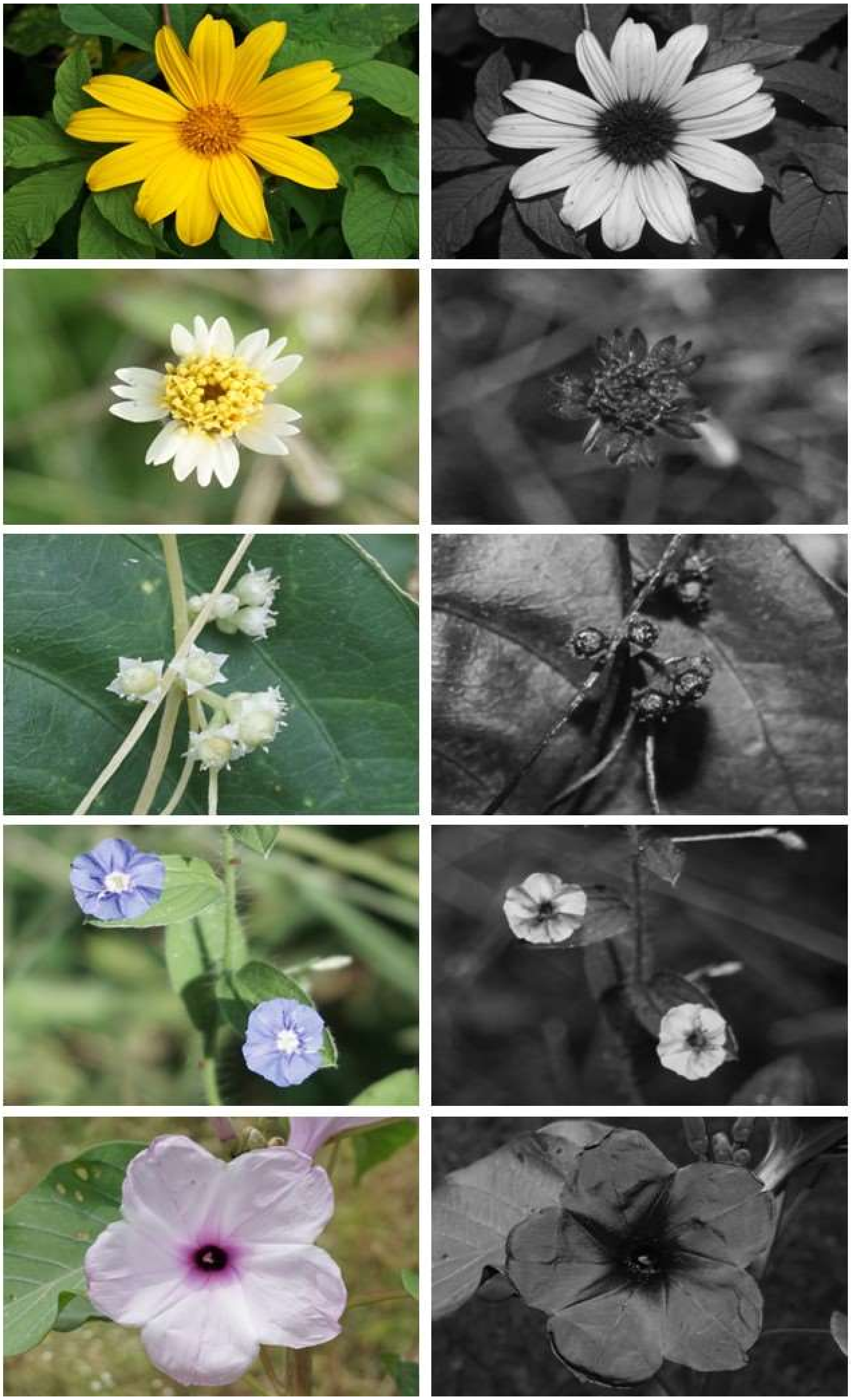

56

57

58

59

60

**Species list (cont.):** 56) *Tithonia diversifolia* (Hemsl.) A.Gray (Compositae), 57) *Tridax procumbens* L. (Compositae), 58) *Cuscuta pentagona* Engelm. (Convolvulaceae), 59) *Evolvulus alsinoides* (L.) L. (Convolvulaceae), and 60) *Ipomoea aquatica* Forssk. (Convolvulaceae).

61

62

63

64

65

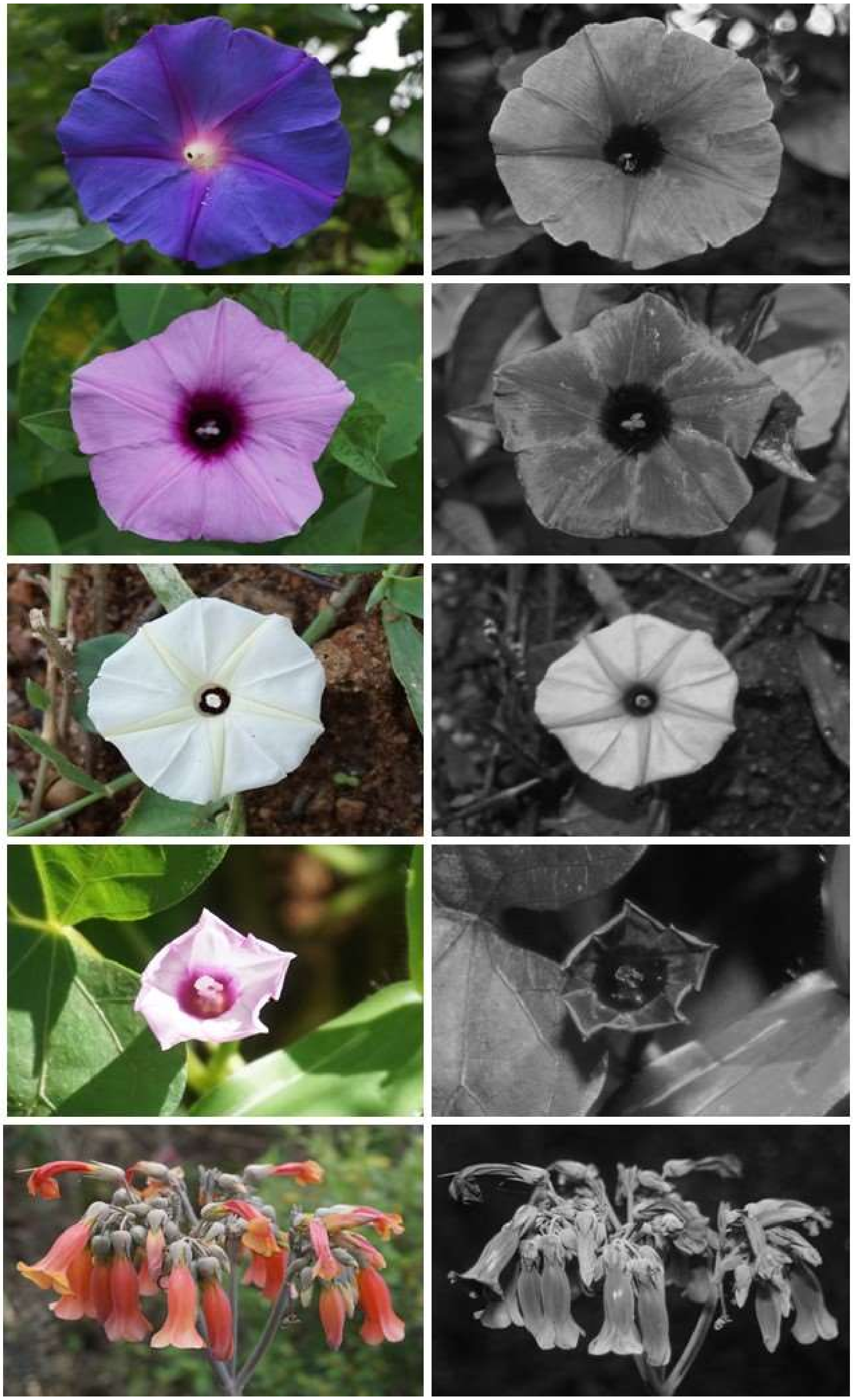

**Species list (cont.):** 61) *Ipomoea indica* (Burm.) Merr. (Convolvulaceae), 62) *Ipomoea littoralis* Blume (Convolvulaceae), 63) *Ipomoea obscura* (L.) Ker Gawl. (Convolvulaceae), 64) *Ipomoea triloba* L. (Convolvulaceae), and 65) *Kalanchoe delagoensis* Eckl. & Zeyh. (Crassulaceae).

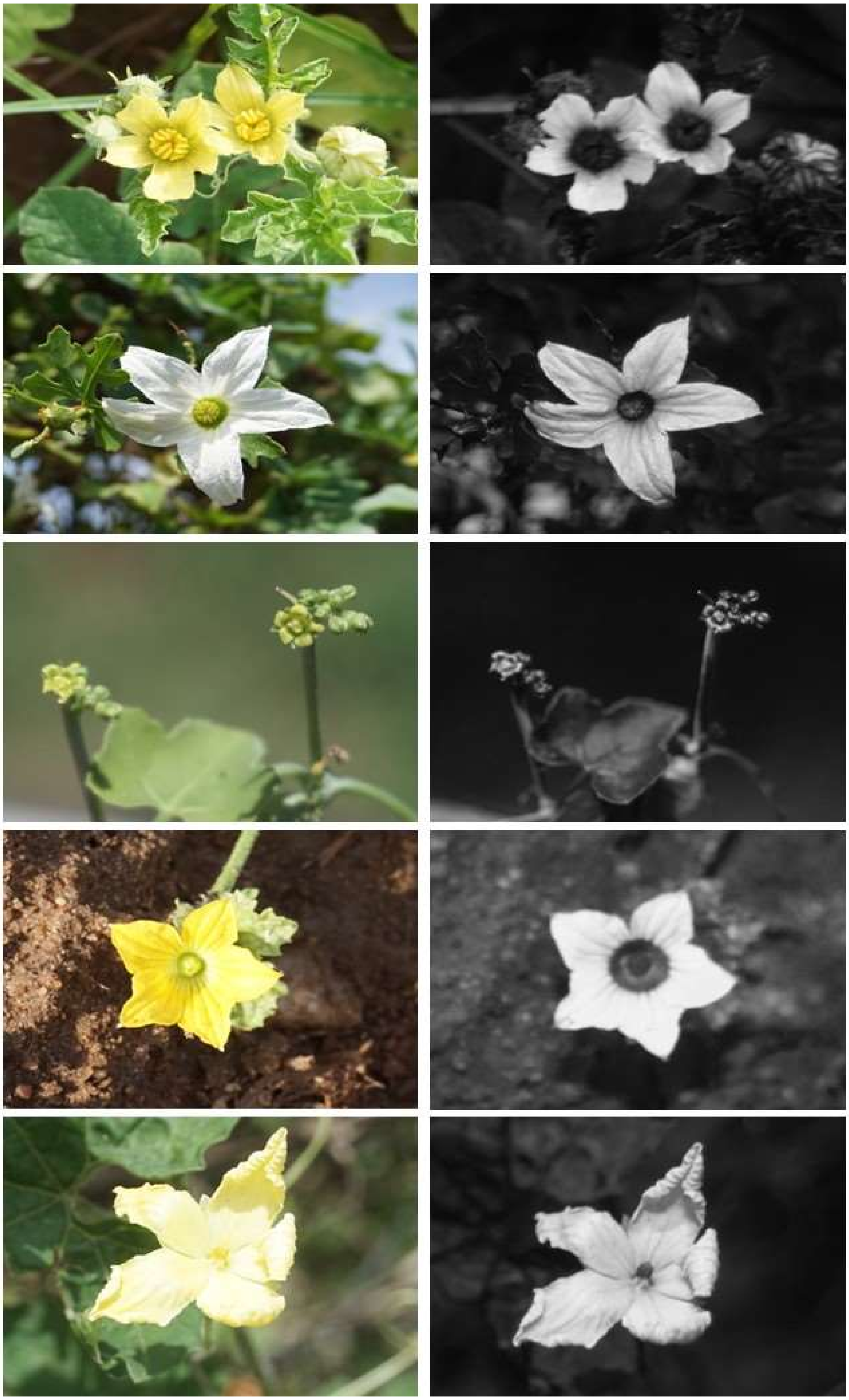

66

67

68

69

70

**Species list (cont.):** 66) *Citrullus lanatus* (Thunb.) Matsum. & Nakai (Cucurbitaceae), 67) *Coccinia grandis* (L.) Voigt (Cucurbitaceae), 68) *Corallocarpus epigaeus* (Rottler) Hook.f. (Cucurbitaceae), 69) *Cucumis melo* L. (Cucurbitaceae), and 70) *Momordica charantia* L. (Cucurbitaceae).

71

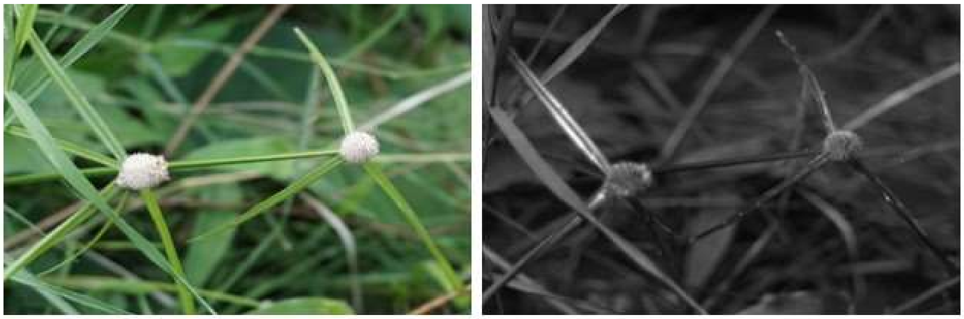

72

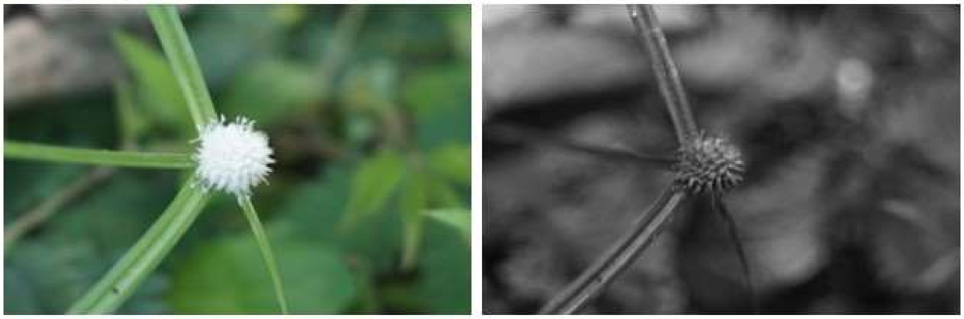

73

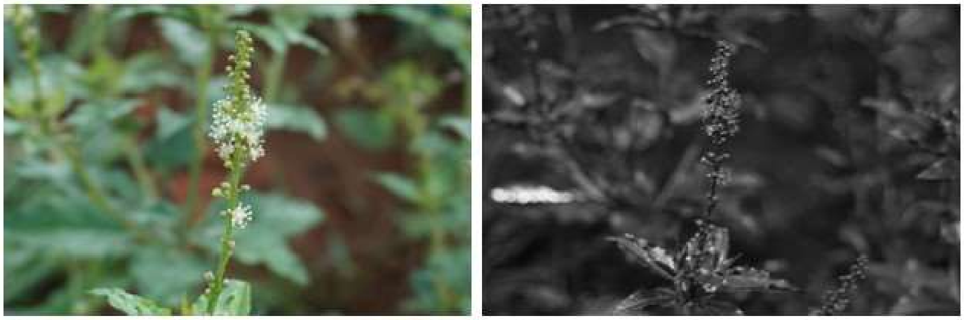

74

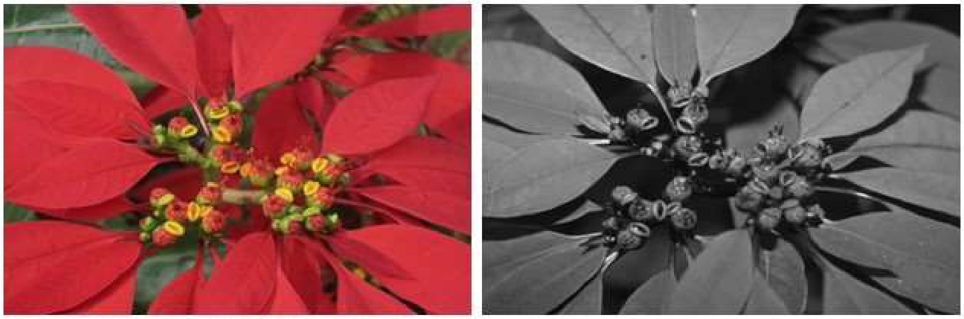

75

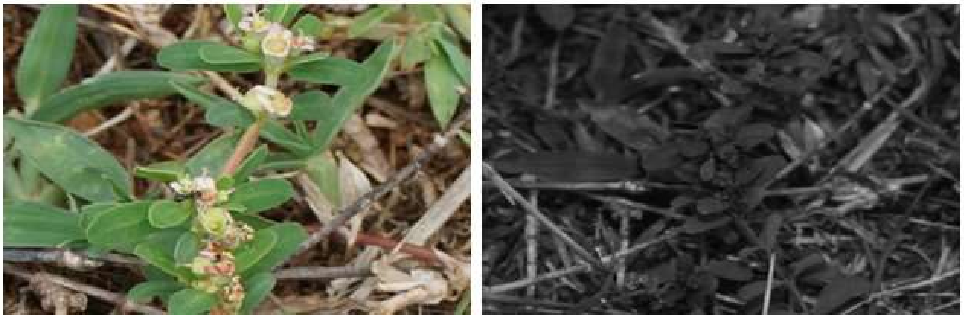

**Species list (cont.):** 71) *Cyperus mindorensis* (Steud.) Huygh (Cyperaceae), 72) *Kyllinga nemoralis* (J.R.Forst. & G.Forst.) Dandy ex Hutch. & Dalziel (Cyperaceae), 73) *Croton bonplandianus* Baill. (Euphorbiaceae), 74) *Euphorbia pulcherrima* Willd. ex Klotzsch (Euphorbiaceae), and 75) *Euphorbia rosea* Retz. (Euphorbiaceae).

76

77

78

79

80

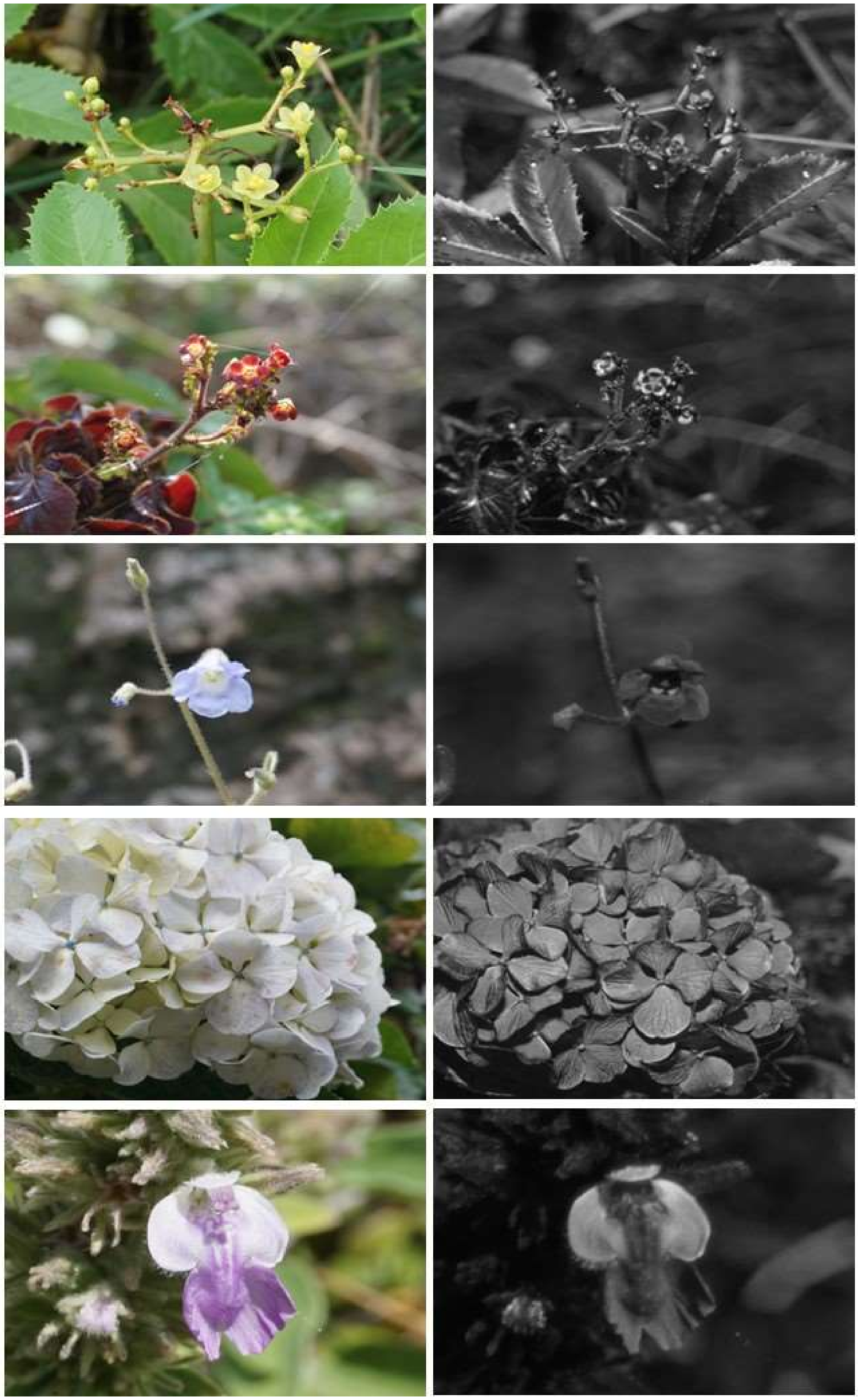

**Species list (cont.):** 76) *Jatropha glandulifera* Roxb. (Euphorbiaceae), 77) *Jatropha gossypiifolia* L. (Euphorbiaceae), 78) *Henckelia humboldtiana* (Gardner) A.Weber & B.L.Burtt (Gesneriaceae), 79) *Hydrangea macrophylla* (Thunb.) Ser. (Hydrangeaceae), and 80) *Anisomeles malabarica* (L.) R.Br. (Lamiaceae).

81

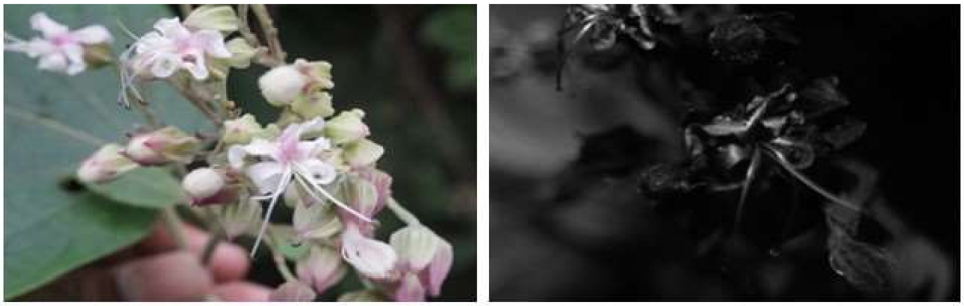

82

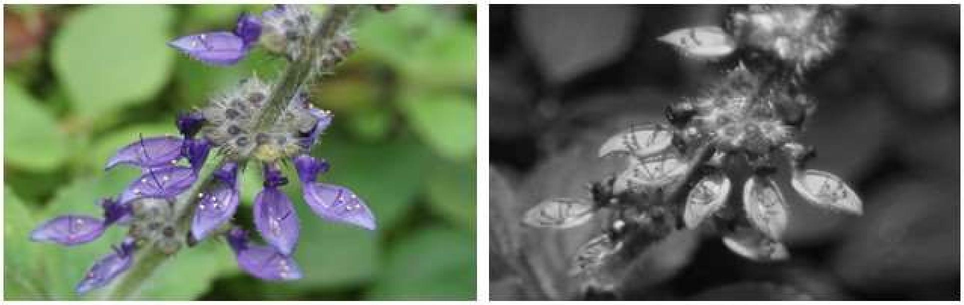

83

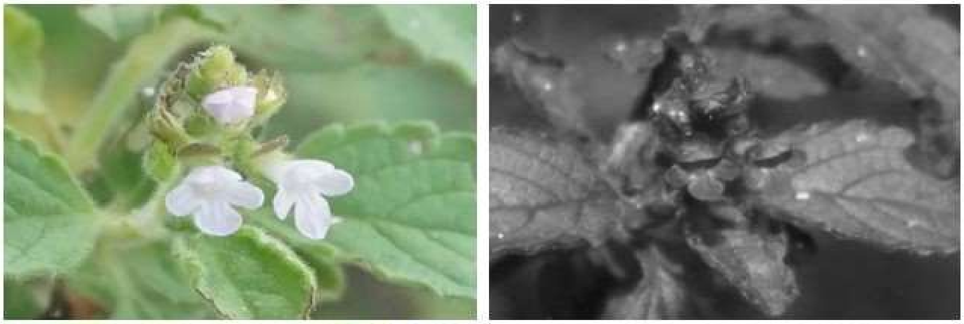

84

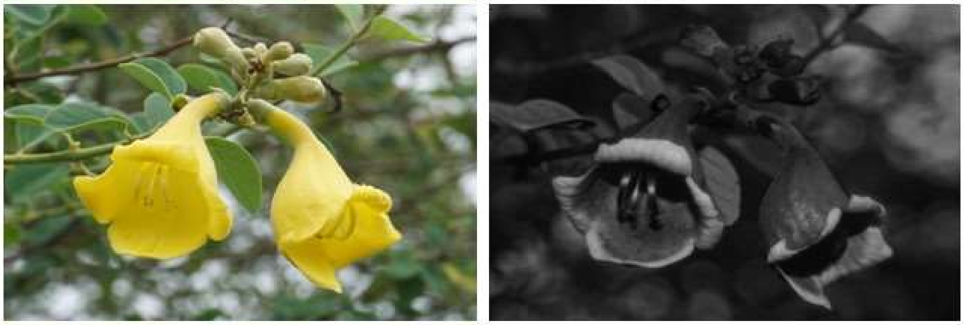

85

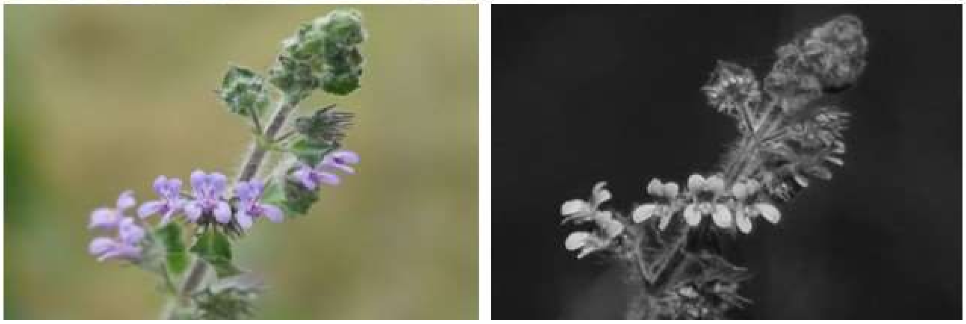

**Species list (cont.):** 81) *Clerodendrum infortunatum* L. (Lamiaceae), 82) *Coleus barbatus* (Andrews) Benth. ex G.Don (Lamiaceae), 83) *Endostemon viscosus* (Roth) M.R.Ashby (Lamiaceae), 84) *Gmelina asiatica* L. (Lamiaceae), and 85) *Hyptis suaveolens* (L.) Poit. (Lamiaceae).

86

87

88

89

90

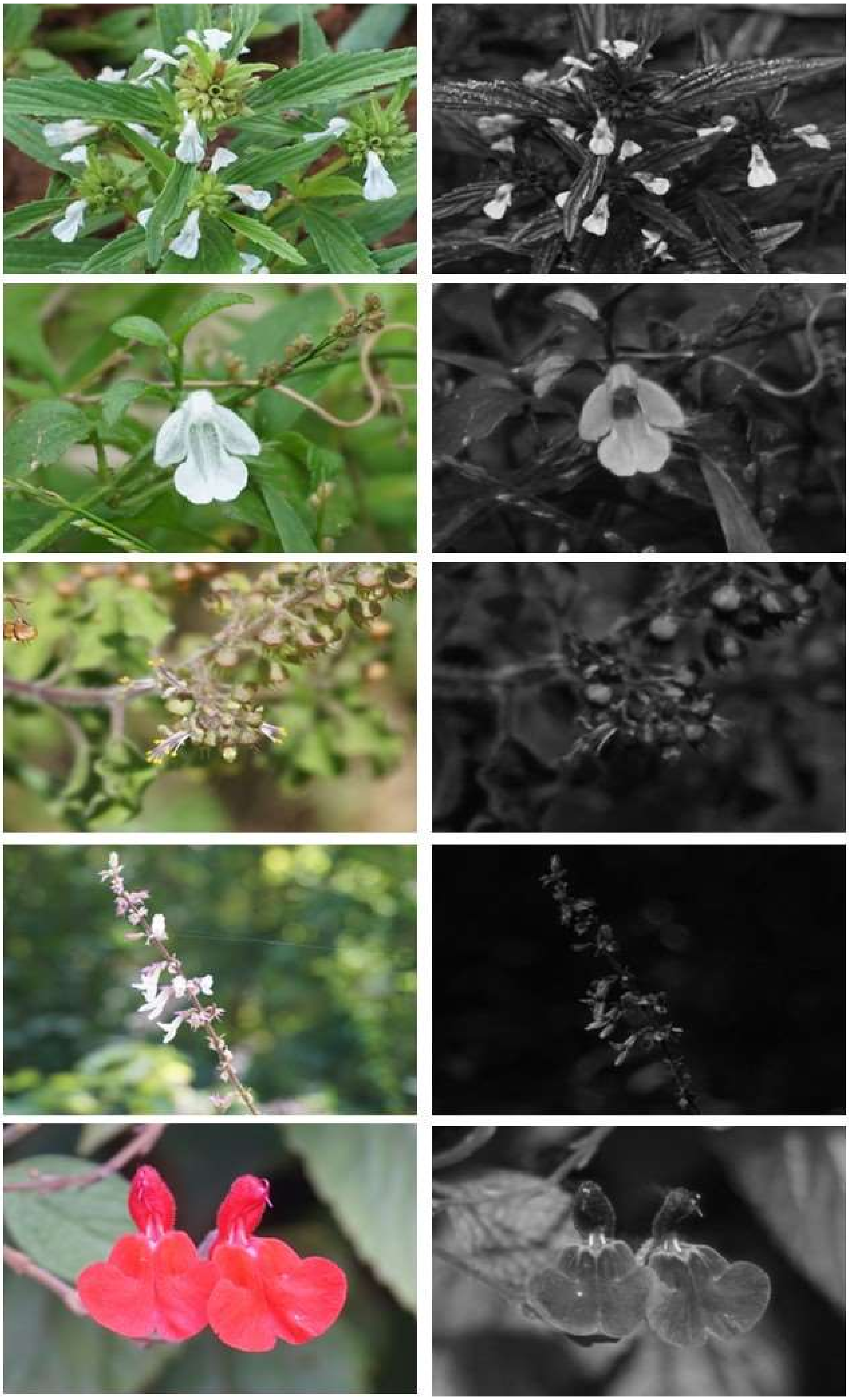

**Species list (cont.):** 86) *Leucas aspera* (Willd.) Link (Lamiaceae), 87) *Leucas biflora* (Vahl) Sm (Lamiaceae), 88) *Ocimum tenuiflorum* L. (Lamiaceae), 89) *Orthosiphon thymiflorus* (Roth) Sleesen (Lamiaceae), and 90) *Salvia coccinea* Buc'hoz ex Etl. (Lamiaceae).

91

92

93

94

95

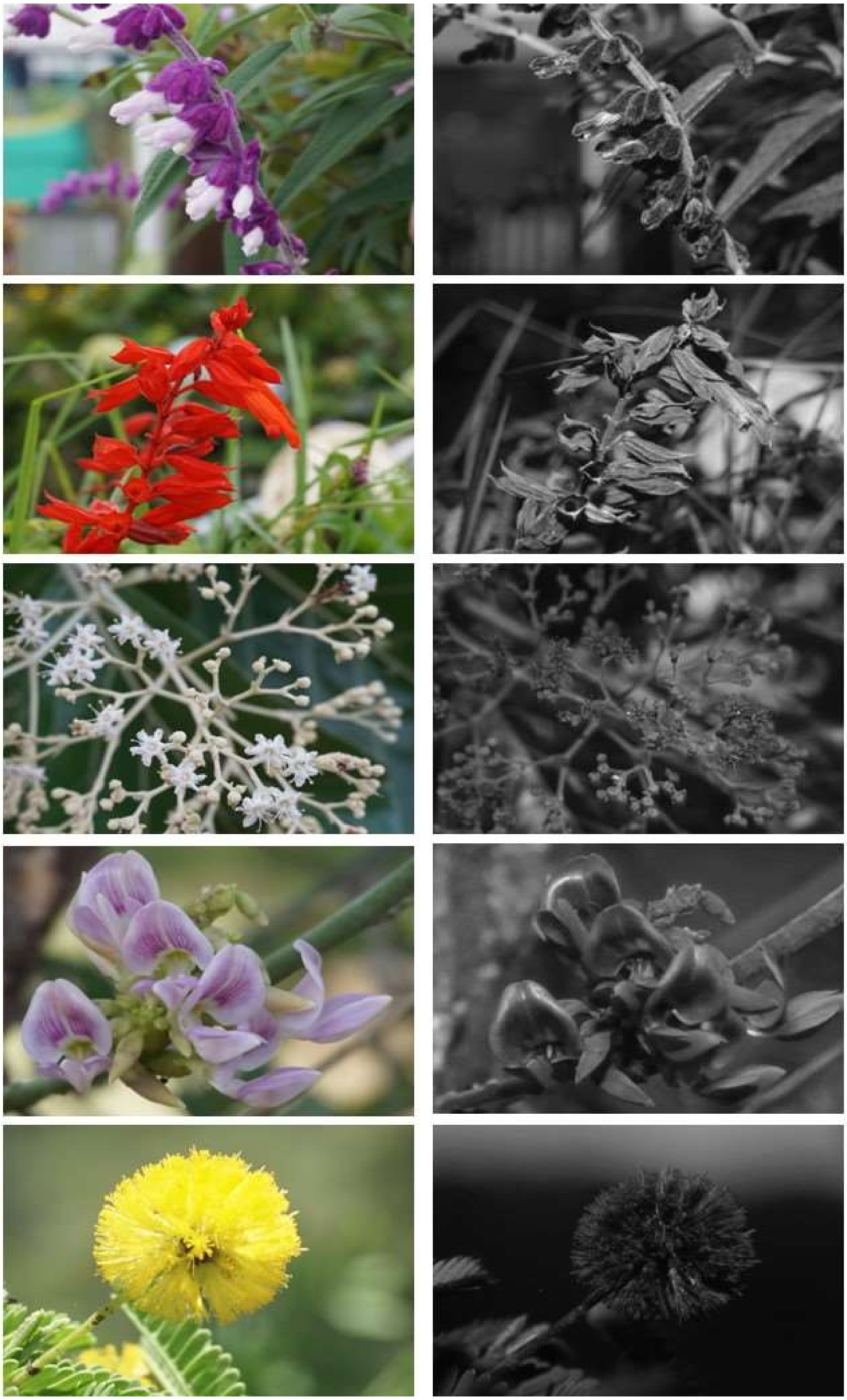

**Species list (cont.):** 91) *Salvia leucantha* Cav. (Lamiaceae), 92) *Salvia splendens* Sellow ex Nees (Lamiaceae), 93) *Tectona grandis* L.f. (Lamiaceae), 94) *Abrus precatorius* L. (Leguminosae), and 95) *Acacia farnesiana* (L.) Willd. (Leguminosae).

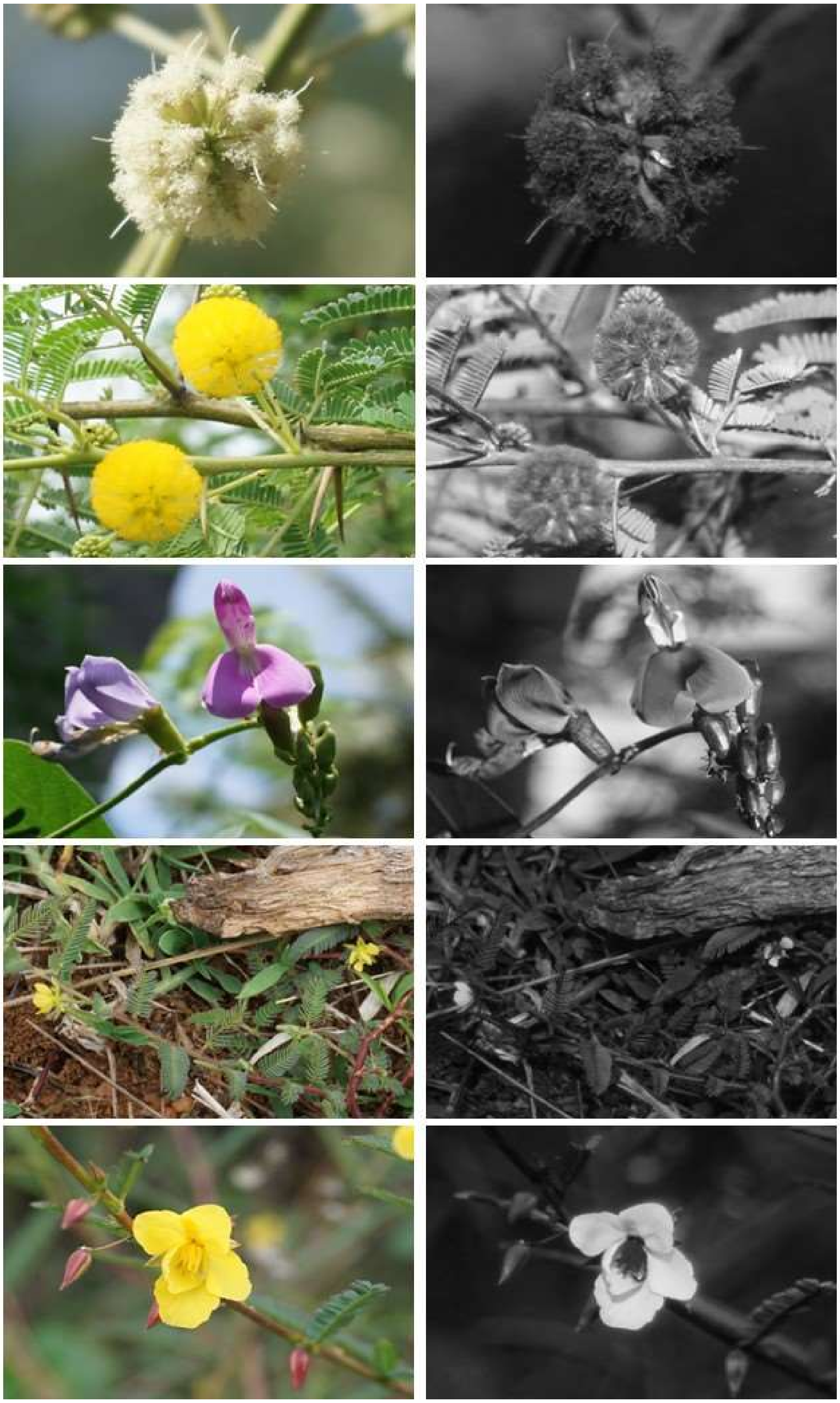


**Species list (cont.):** 96) *Acacia leucophloea* (Roxb.) Willd. (Leguminosae), 97) *Acacia nilotica* (L.) Willd. ex Delile (Leguminosae), 98) *Canavalia gladiate* (Jacq.) DC. (Leguminosae), 99) *Cassia mimosoides* L. (Leguminosae), and 100) *Chamaecrista pumila* (Lam.) V.Singh (Leguminosae).

101

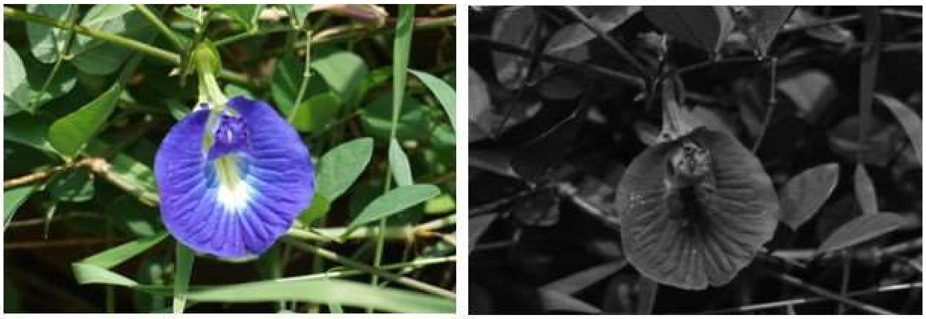

102

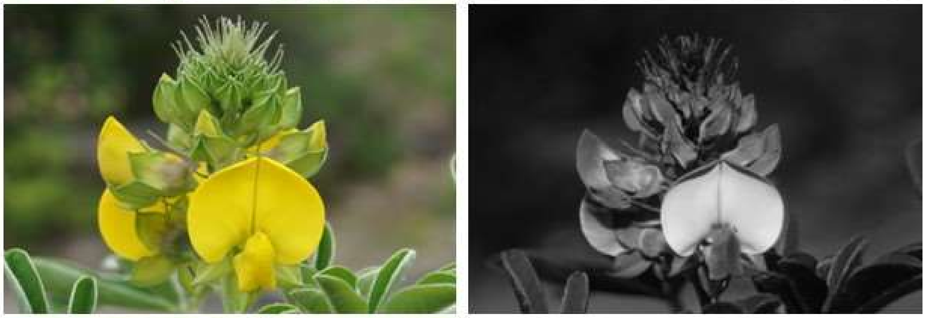

103

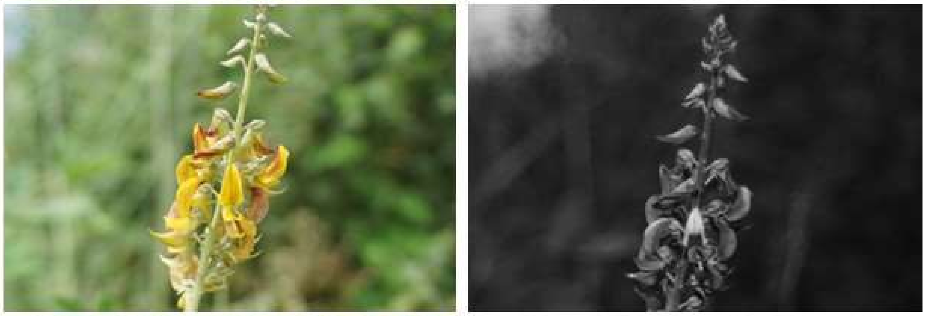

104

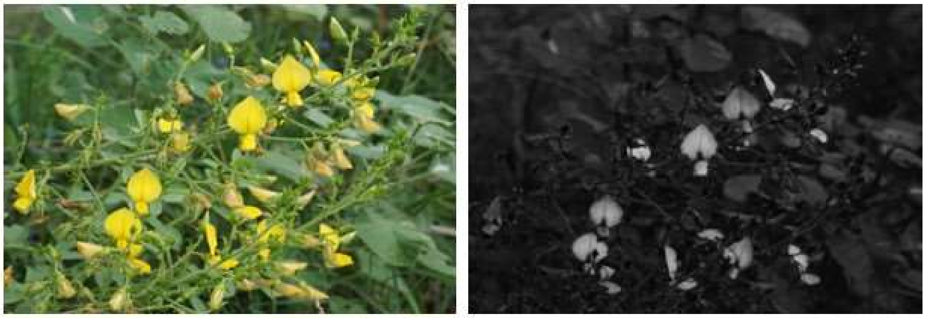

105

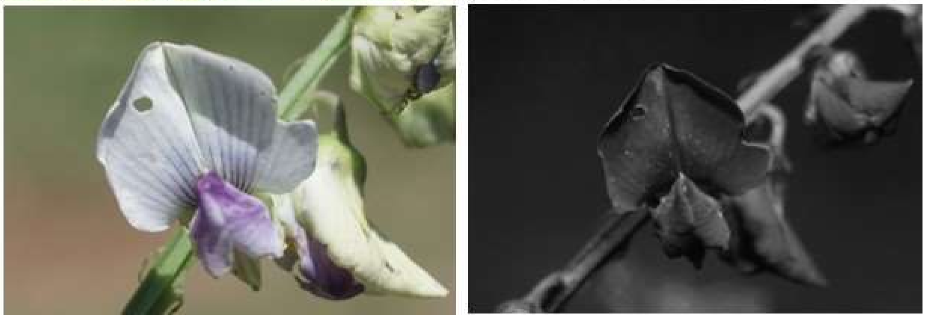

**Species list (cont.):** 101) *Clitoria ternatea* L. (Leguminosae), 102) *Crotalaria grahamiana* Wight & Arn. (Leguminosae), 103) *Crotalaria pallida* Aiton (Leguminosae), 104) *Crotalaria paniculata* Willd. (Leguminosae), and 105) *Crotalaria verrucosa* L. (Leguminosae).

106

107

108

109

110

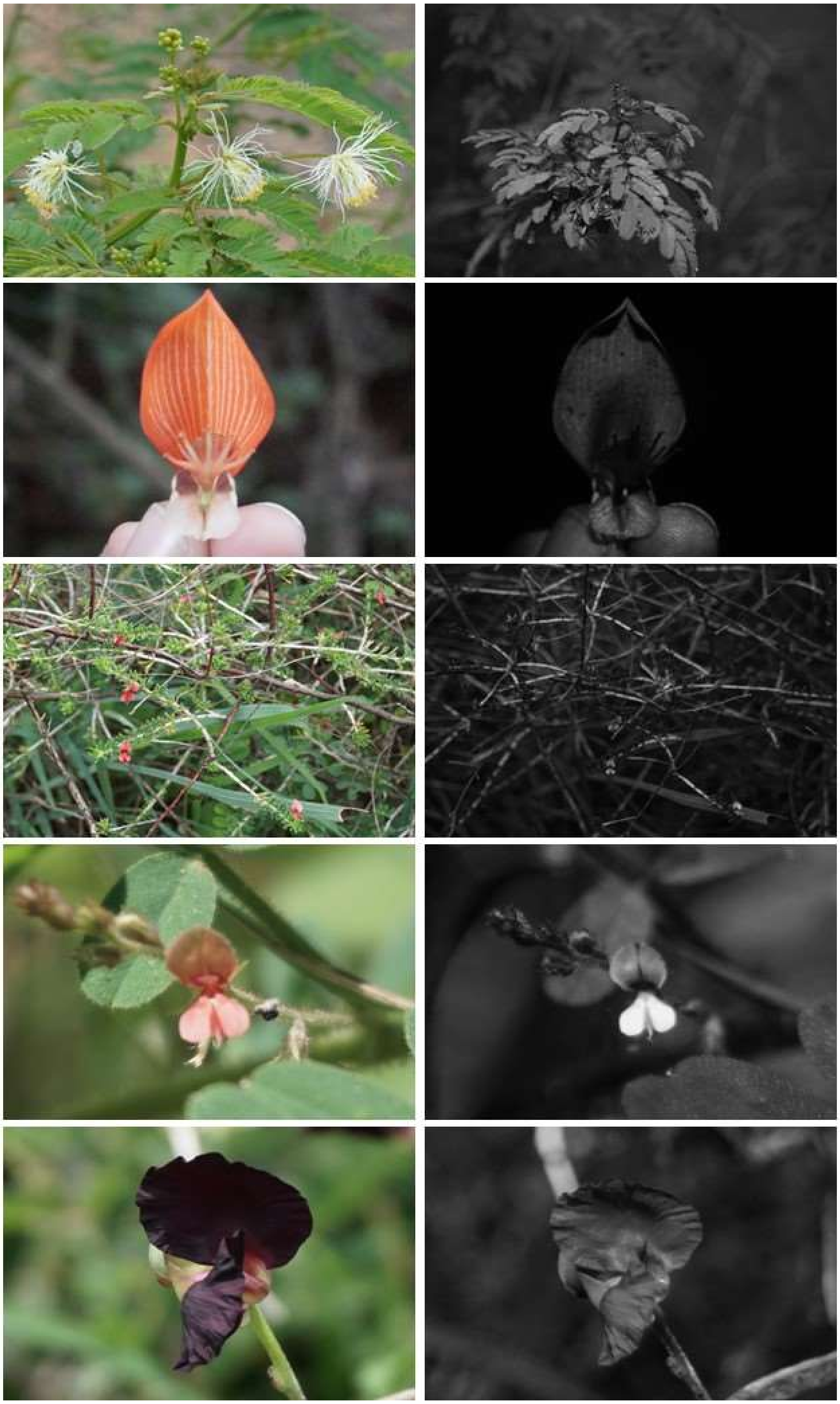

**Species list (cont.):** 106) *Desmanthus virgatus* (L.) Willd. (Leguminosae), 107) *Erythrina variegate* L. (Leguminosae), 108) *Indigofera aspalathoides* Vahl ex DC. (Leguminosae), 109) *Indigofera tinctoria* Forssk. (Leguminosae), and 110) *Macroptilium atropurpureum* (DC.) Urb. (Leguminosae).

111

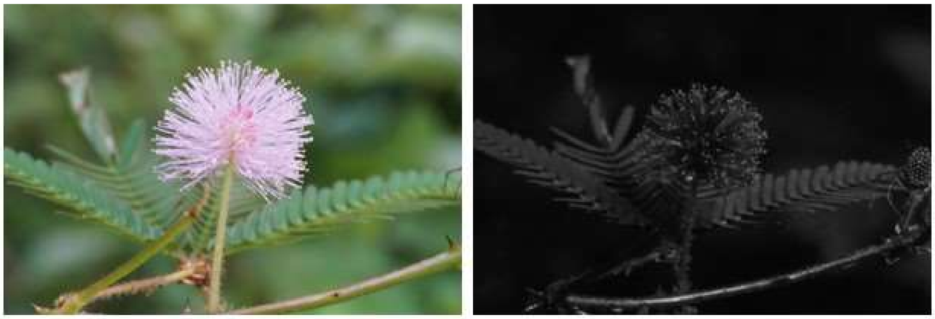

112

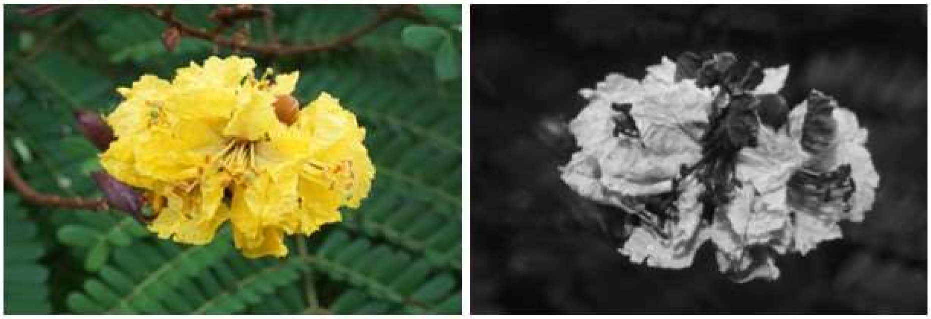

113

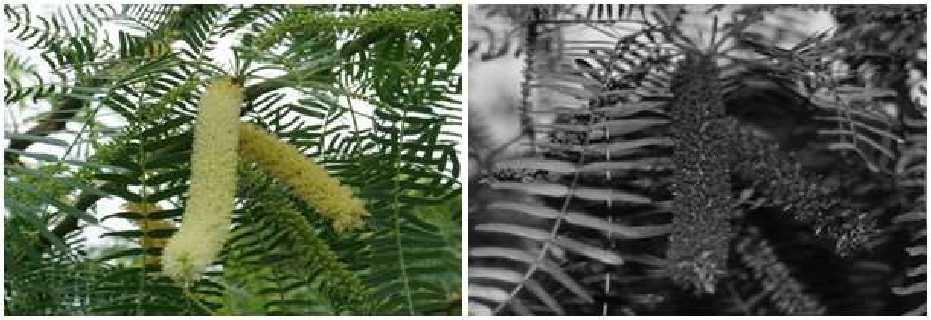

114

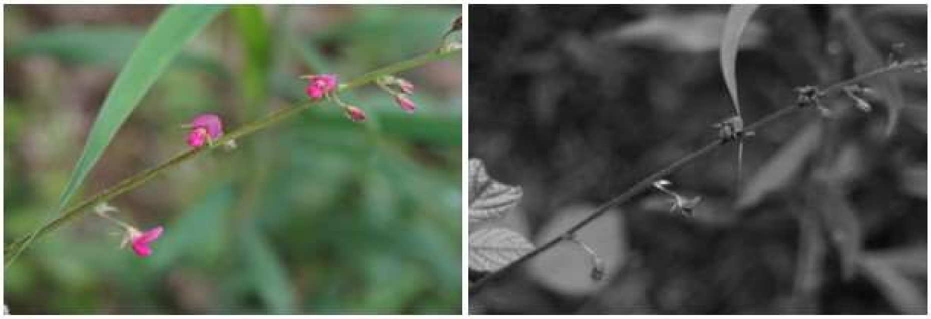

115

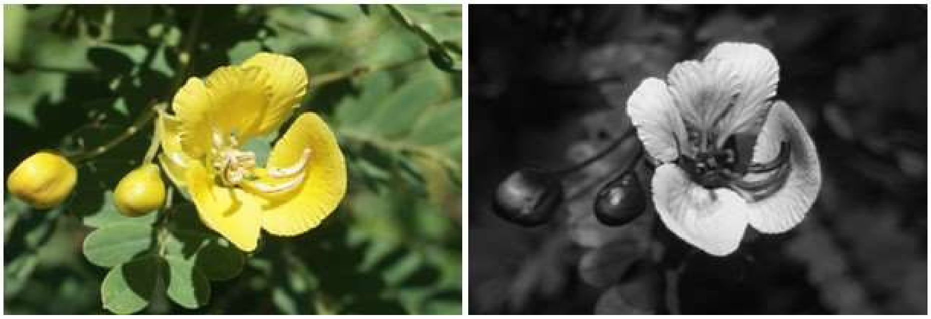

**Species list (cont.):** 111) *Mimosa pudica* L. (Leguminosae), 112) *Peltophorum pterocarpum* (DC.) Backer ex K.Heyne (Leguminosae), 113) *Prosopis juliflora* (Sw.) DC. (Leguminosae), 114) *Pseudarthria viscida* (L.) Wight & Arn. (Leguminosae), and 115) *Senna auriculata* (L.) Roxb. (Leguminosae).

116

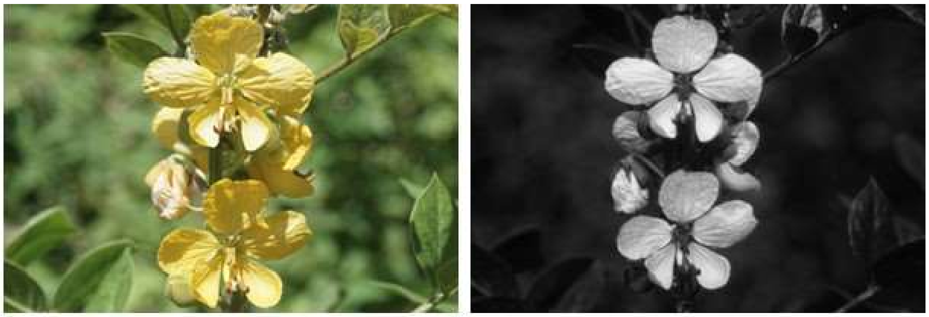

117

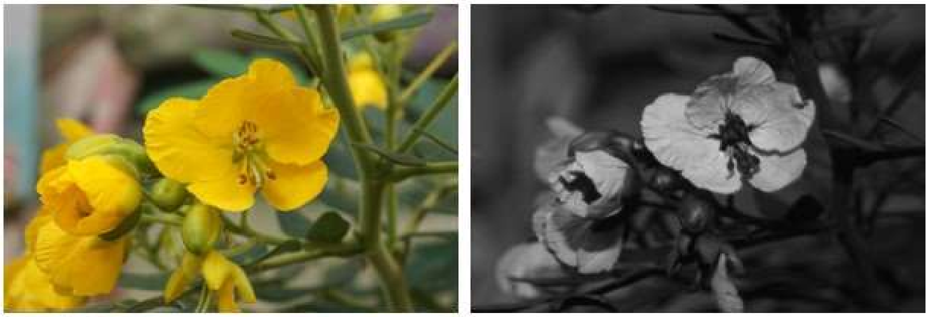

118

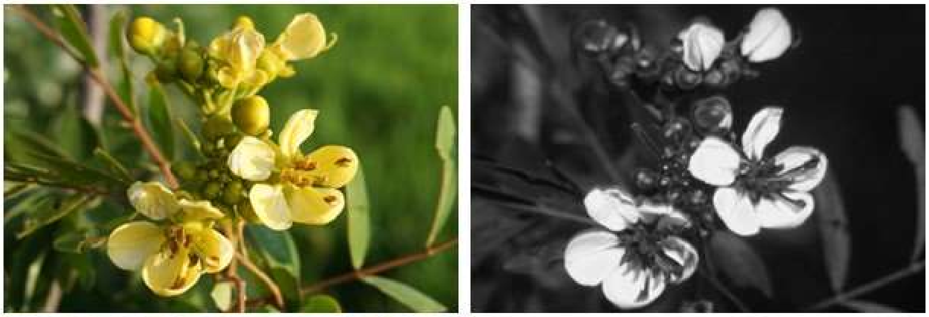

119

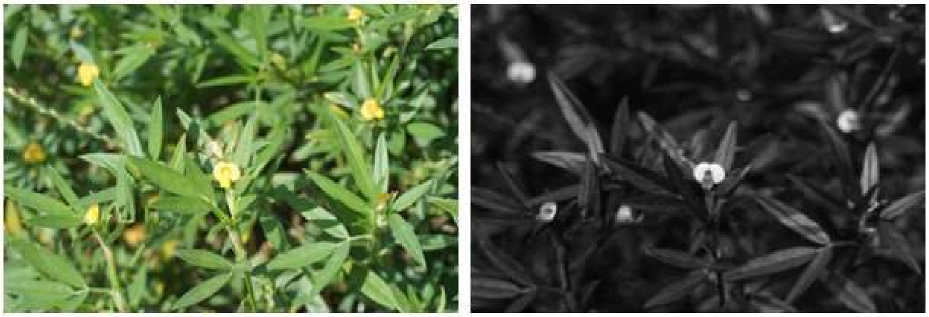

120

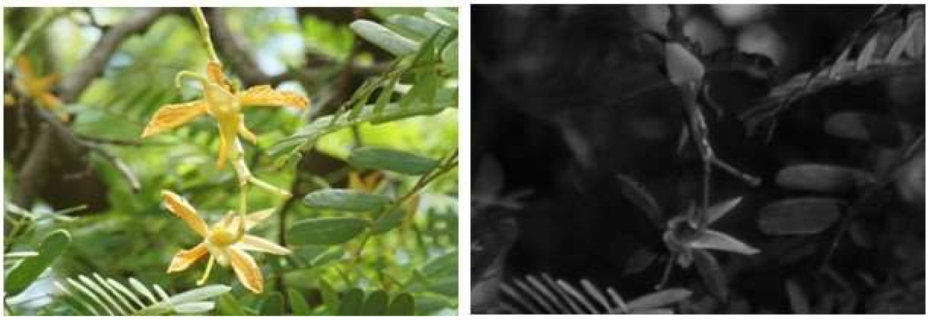

**Species list (cont.):** 116) *Senna occidentalis* (L.) Link (Leguminosae), 117) *Senna septemtrionalis* (Viv.) H.S.Irwin & Barneby (Leguminosae), 118) *Senna siamea* (Lam.) H.S.Irwin & Barneby (Leguminosae), 119) *Stylosanthes mucronata* Willd. (Leguminosae), and 120) *Tamarindus indica* L. (Leguminosae).

121

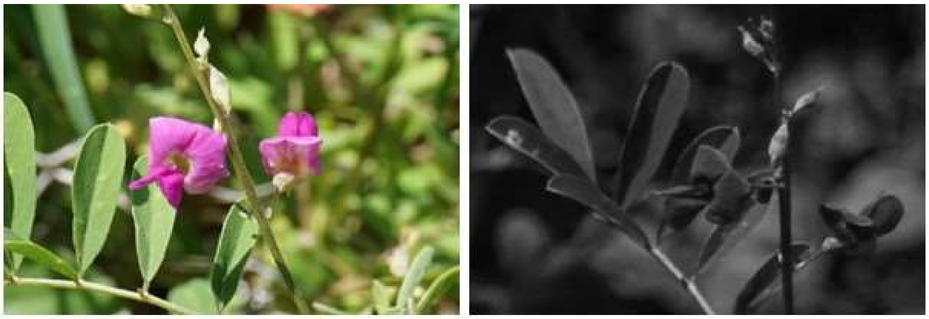

122

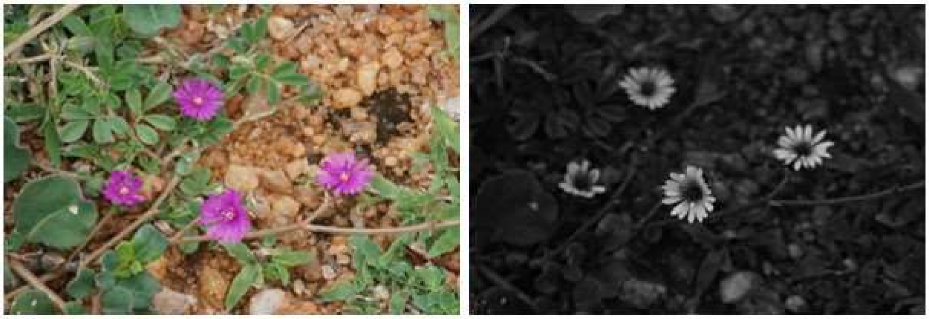

123

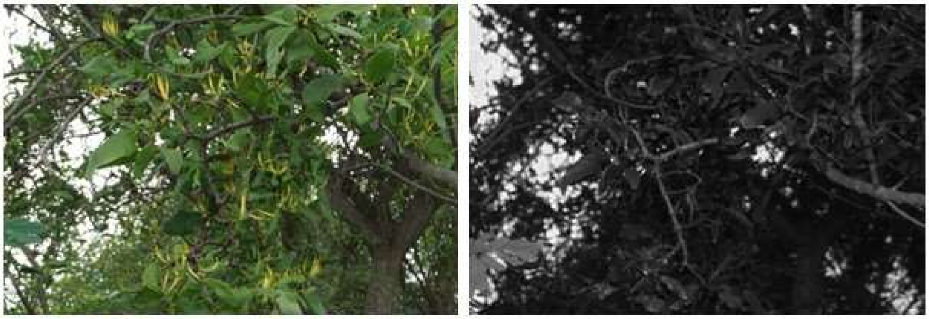

124

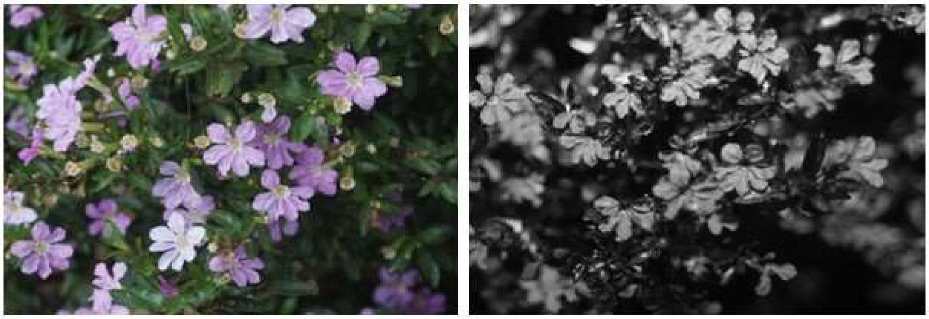

125

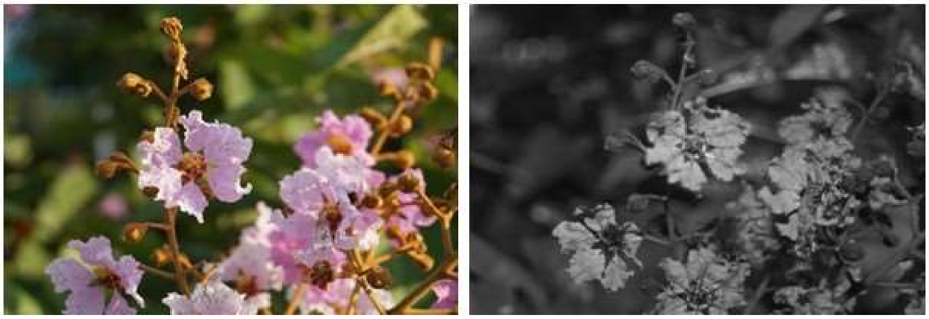

**Species list (cont.):** 121) *Tephrosia purpurea* (L.) Pers. (Leguminosae), 122) *Corbichonia decumbens* (Forssk.) Exell (Lophiocarpaceae), 123) *Dendrophthoe falcata* (L.f.) Ettingsh. (Loranthaceae), 124) *Cuphea hyssopifolia* Griseb. (Lythraceae), and 125) *Lagerstroemia speciosa* (L.) Pers. (Lythraceae).

126

127

128

129

130

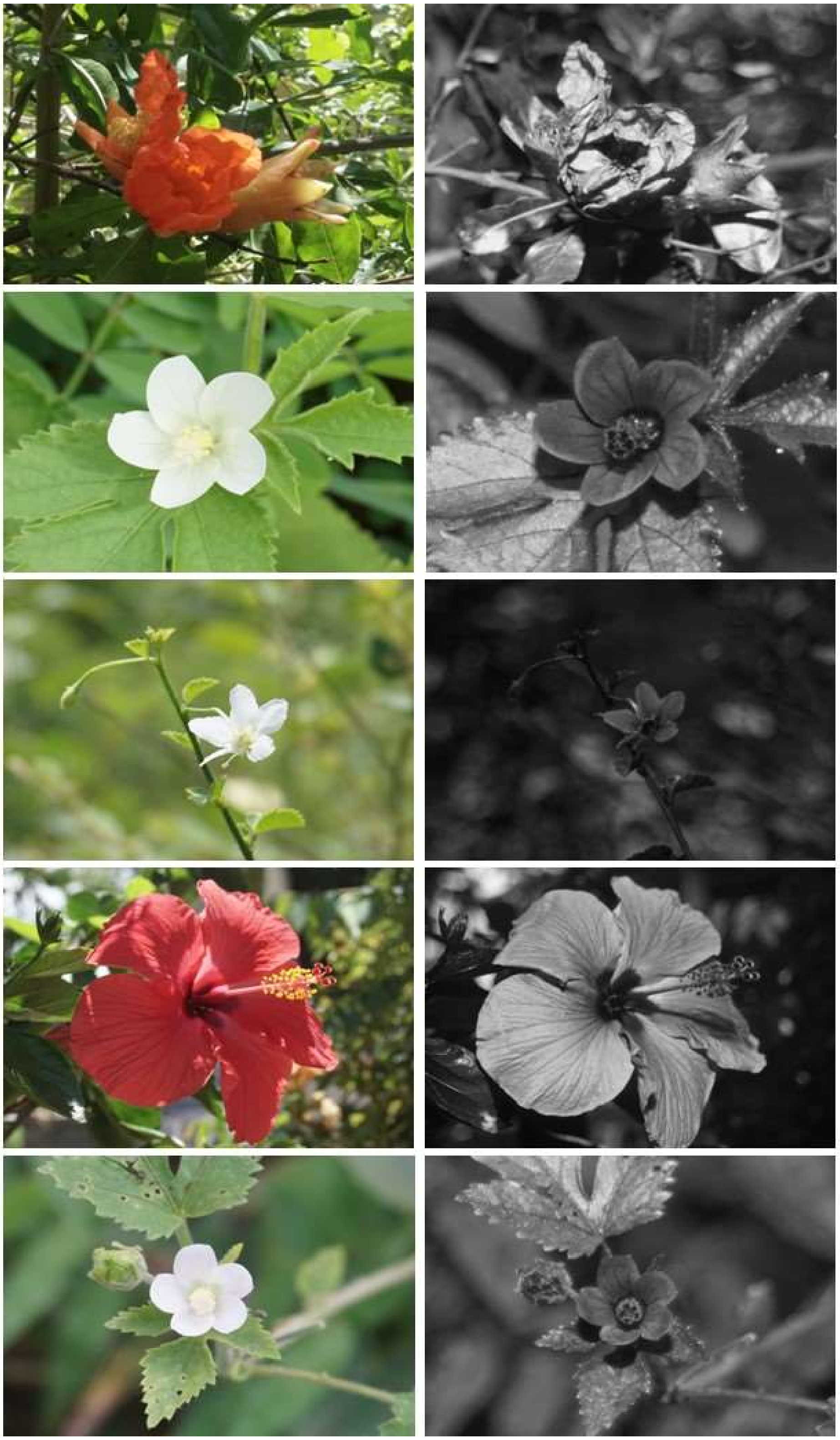

**Species list (cont.):** 126) *Punica granatum* L. (Lythraceae), 127) *Hibiscus calyphyllus* Cav. (Malvaceae), 128) *Hibiscus micranthus* L.f. (Malvaceae), 129) *Hibiscus rosa-sinensis* L. (Malvaceae), and 130) *Hibiscus solandra* L'Hér. (Malvaceae).

131

132

133

134

135

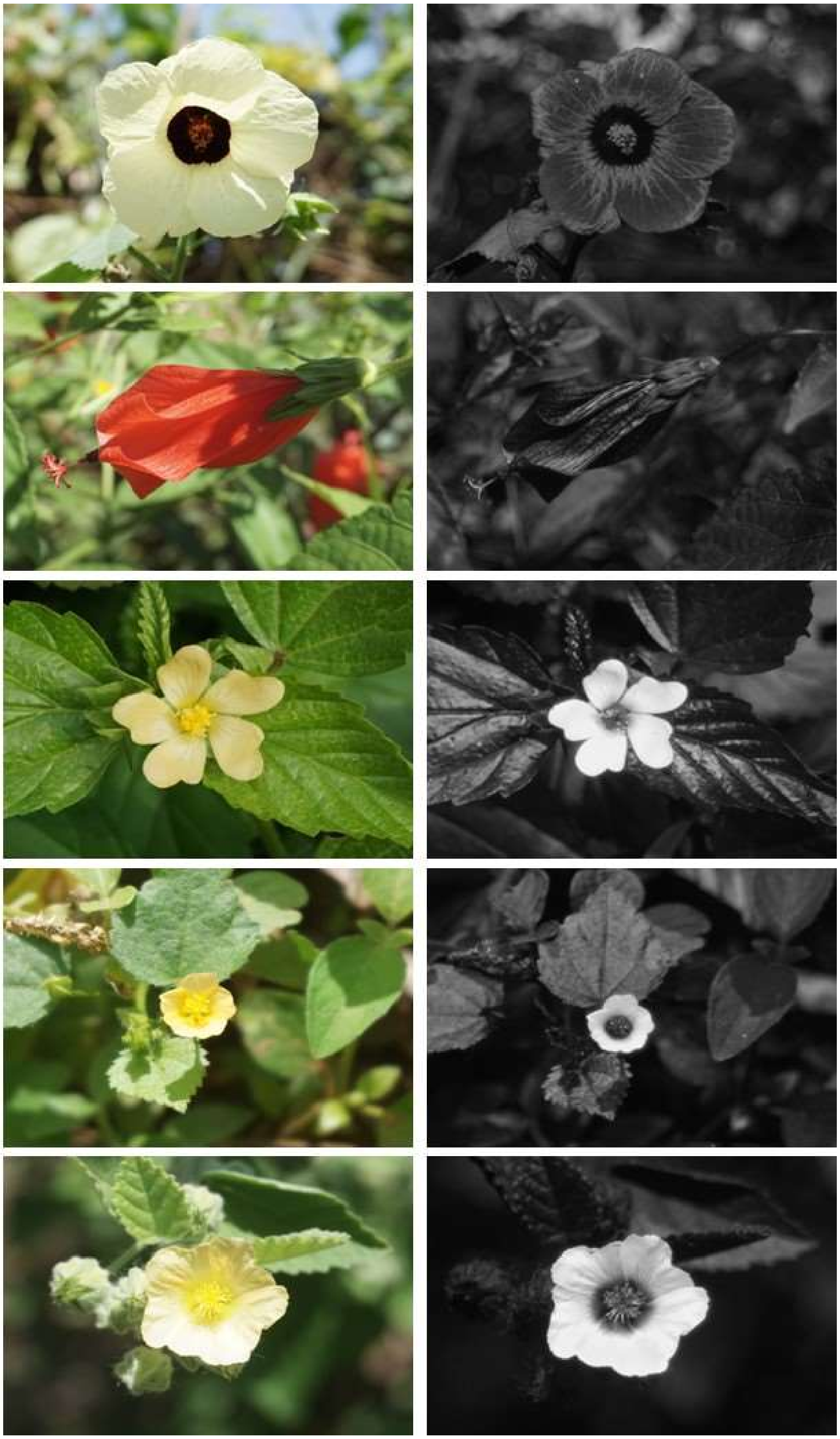

**Species list (cont.):** 131) *Hibiscus vitifolius* Mill. (Malvaceae), 132) *Malvaviscus arboreus var. arboreus* Cav. (Malvaceae), 133) *Sida acuta* Burm.f. (Malvaceae), 134) *Sida cordata* (Burm.f.) Borss.Waalk. (Malvaceae), and 135) *Sida cordifolia* L. (Malvaceae).

136

137

138

139

140

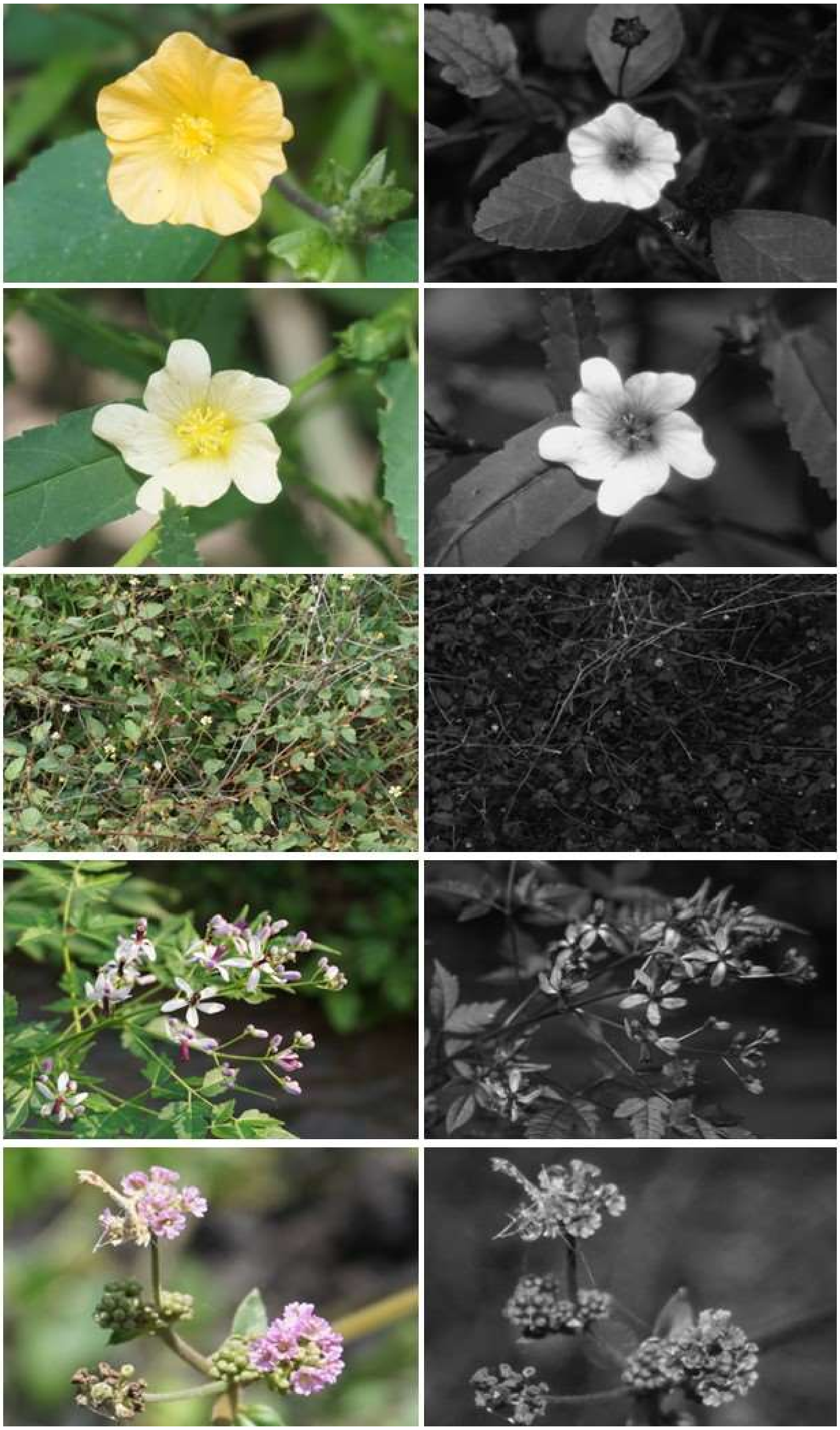

**Species list (cont.):** 136) *Sida ovata* Forssk. (Malvaceae), 137) *Sida rhombifolia* L. (Malvaceae), 138) *Waltheria indica* L. (Malvaceae), 139) *Melia azedarach* L. (Meliaceae), and 140) *Boerhavia coccinea* Mill. (Nyctaginaceae).

141

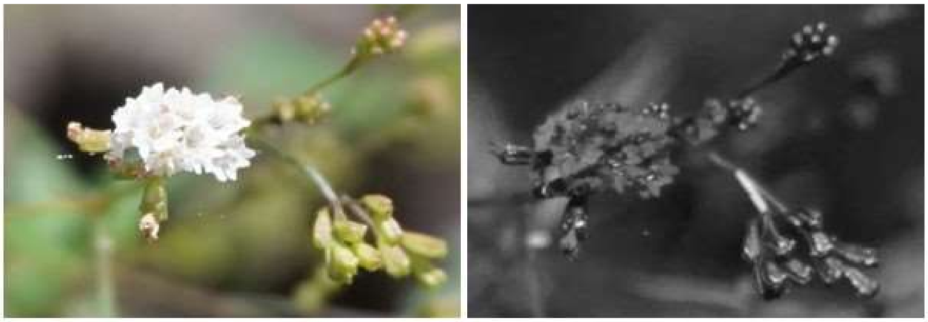

142

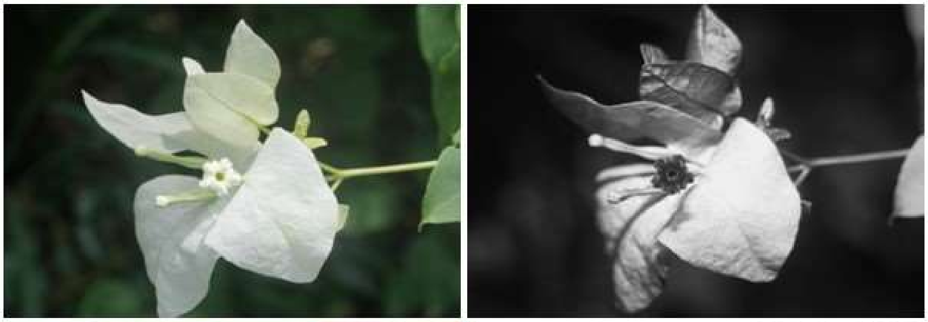

143

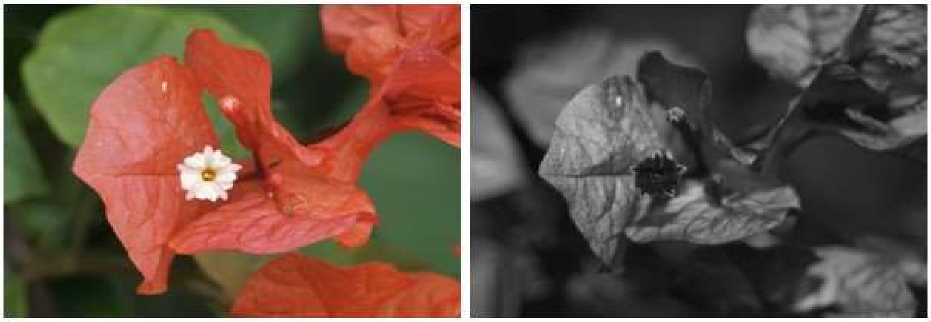

144

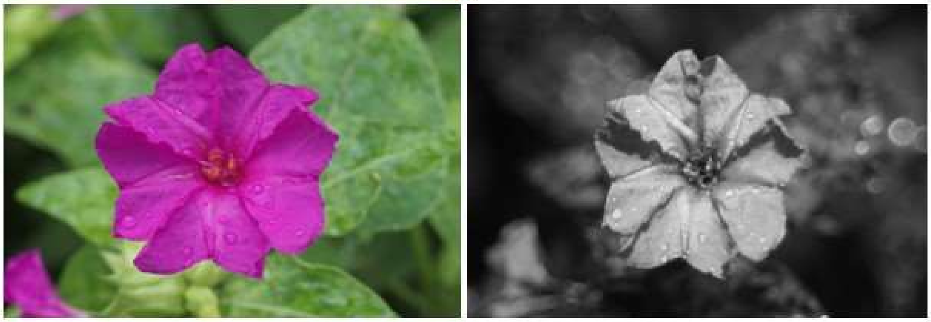

145

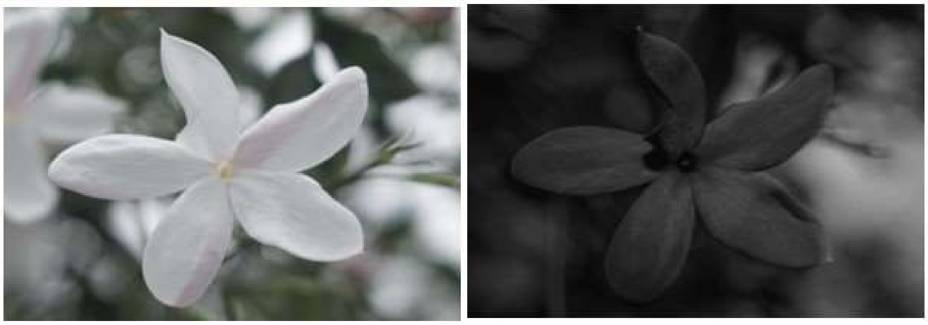

**Species list (cont.):** 141) *Boerhavia erecta* L. (Nyctaginaceae), 142) *Bougainvillea glabra* Choisy (Nyctaginaceae), 143) *Bougainvillea spectabilis* Willd. (Nyctaginaceae), 144) *Mirabilis jalapa* L. (Nyctaginaceae), and 145) *Jasminum griffithianum* L. (Oleaceae).

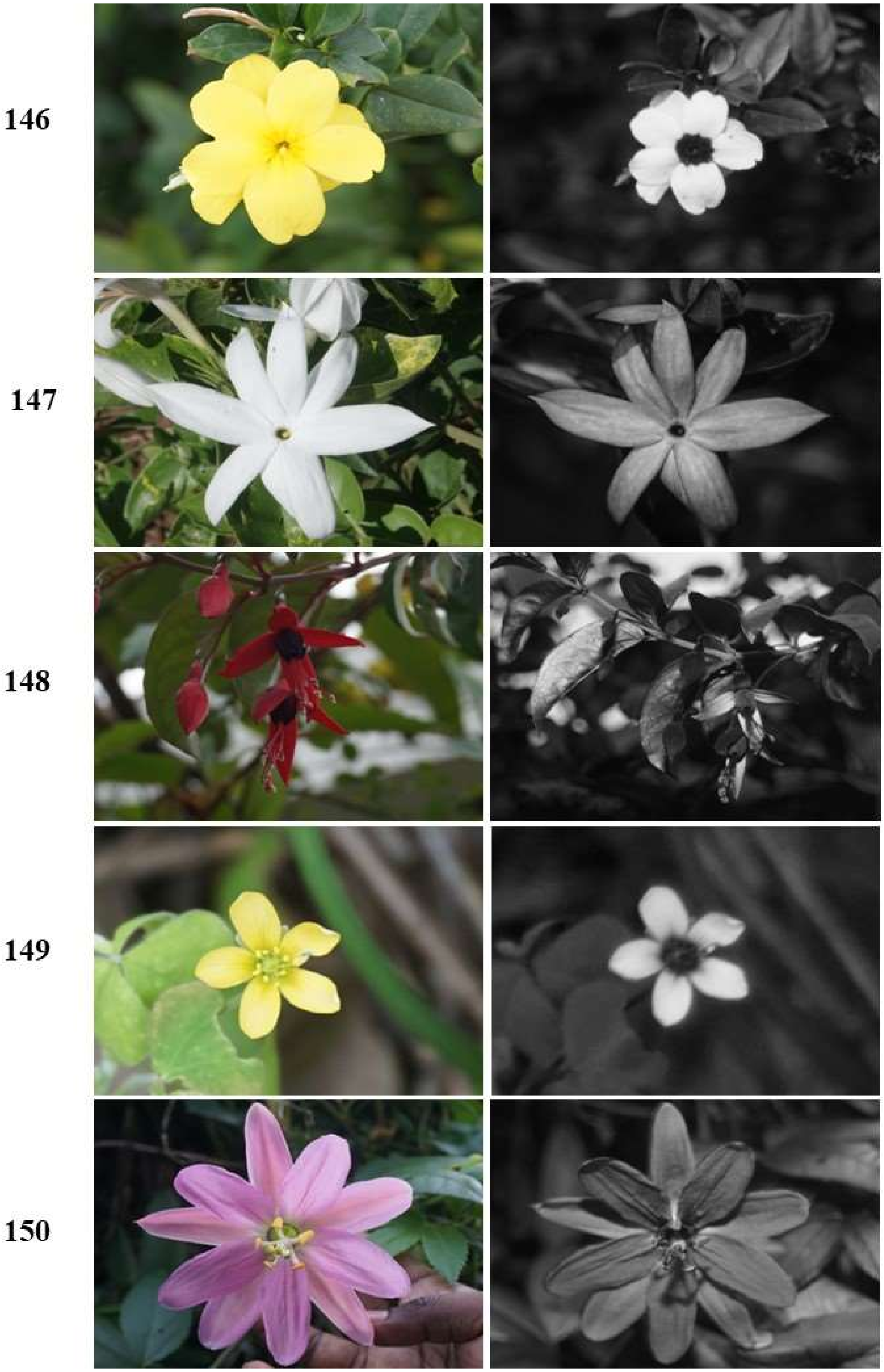


**Species list (cont.):** 146) *Jasminum mesnyi* Hance (Oleaceae), 147) *Jasminum multiflorum* (Burm.f.) Andrews (Oleaceae), 148) *Fuchsia magellanica* Lam. (Onagraceae), 149) *Oxalis corniculata* L. (Oxalidaceae), and 150) *Passiflora tarminiana* Coppens & V.E.Barney (Passifloraceae).

151

152

153

154

155

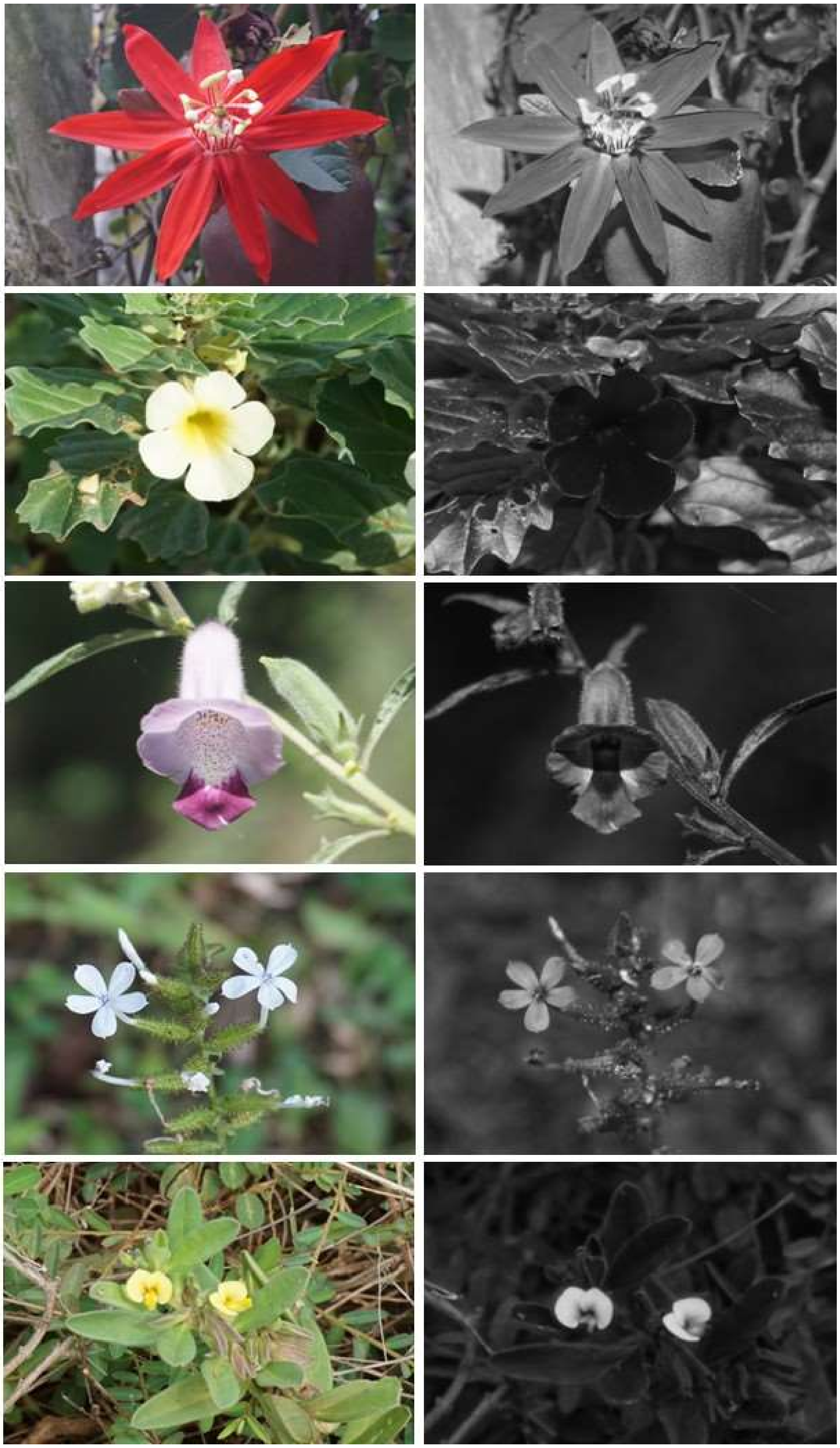

**Species list (cont.):** 151) *Passiflora vitifolia* Kunth (Passifloraceae), 152) *Pedalium murex* L. (Pedaliaceae), 153) *Sesamum radiatum* Thonn. ex Hornem. (Pedaliaceae), 154) *Plumbago zeylanica* L. (Plumbaginaceae), and 155) *Polygala arvensis* Willd. (Polygalaceae).

156

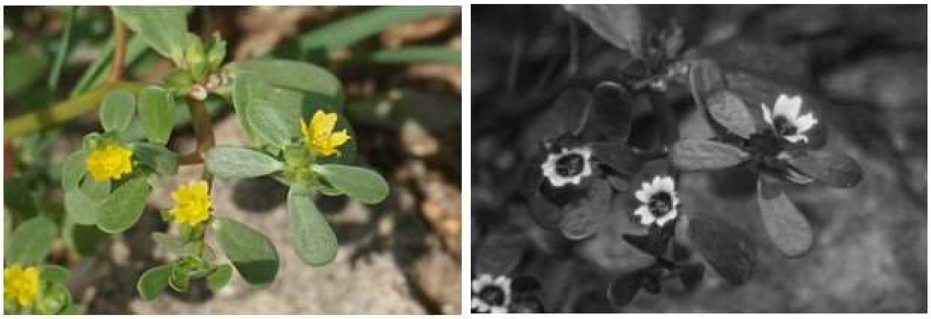

157

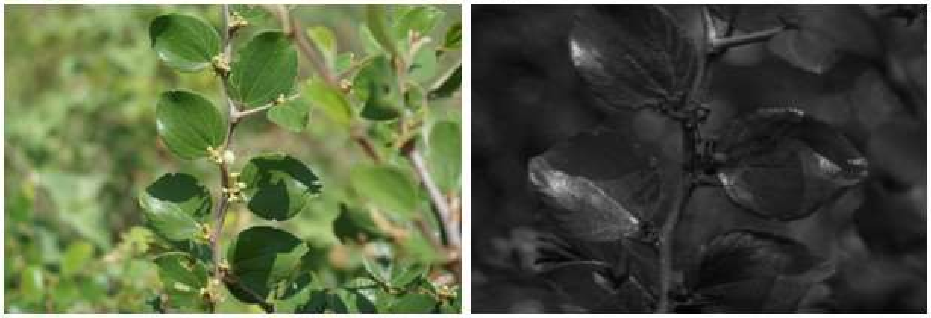

158

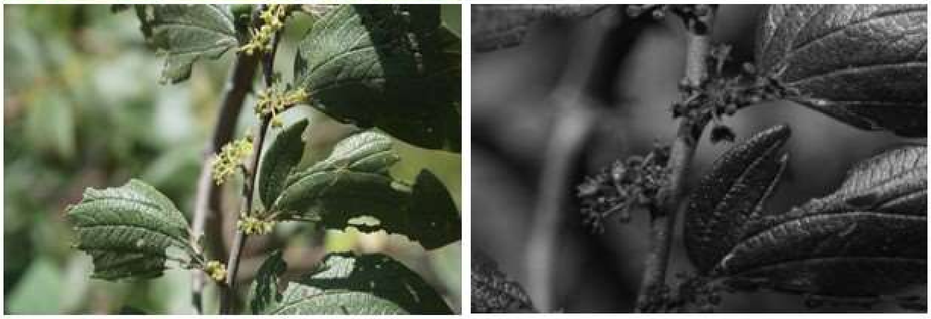

159

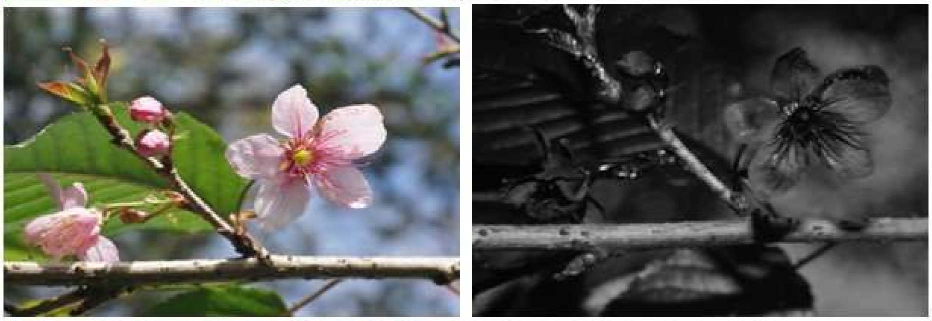

160

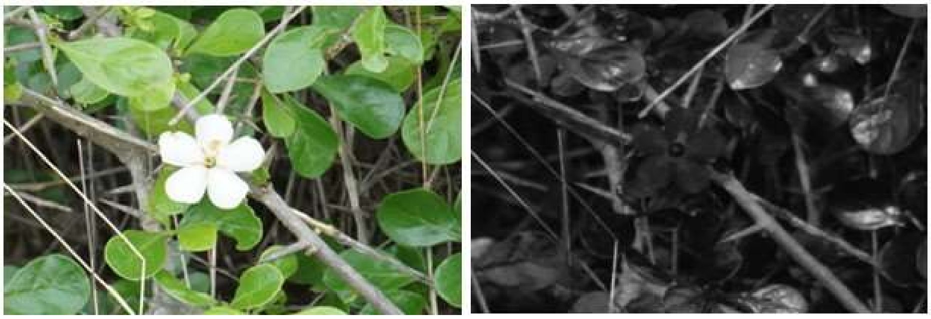

**Species list (cont.):** 156) *Portulaca oleracea* L. (Portulacaceae), 157) *Ziziphus nummularia* (Burm.f.) Wight & Arn. (Rhamnaceae), 158) *Ziziphus oenopolia* (L.) Mill. (Rhamnaceae), 159) *Prunus cerasoides* Koidz. (Rosaceae), and 160) *Catunaregam spinosa* (Thunb.) Tirveng. (Rubiaceae).

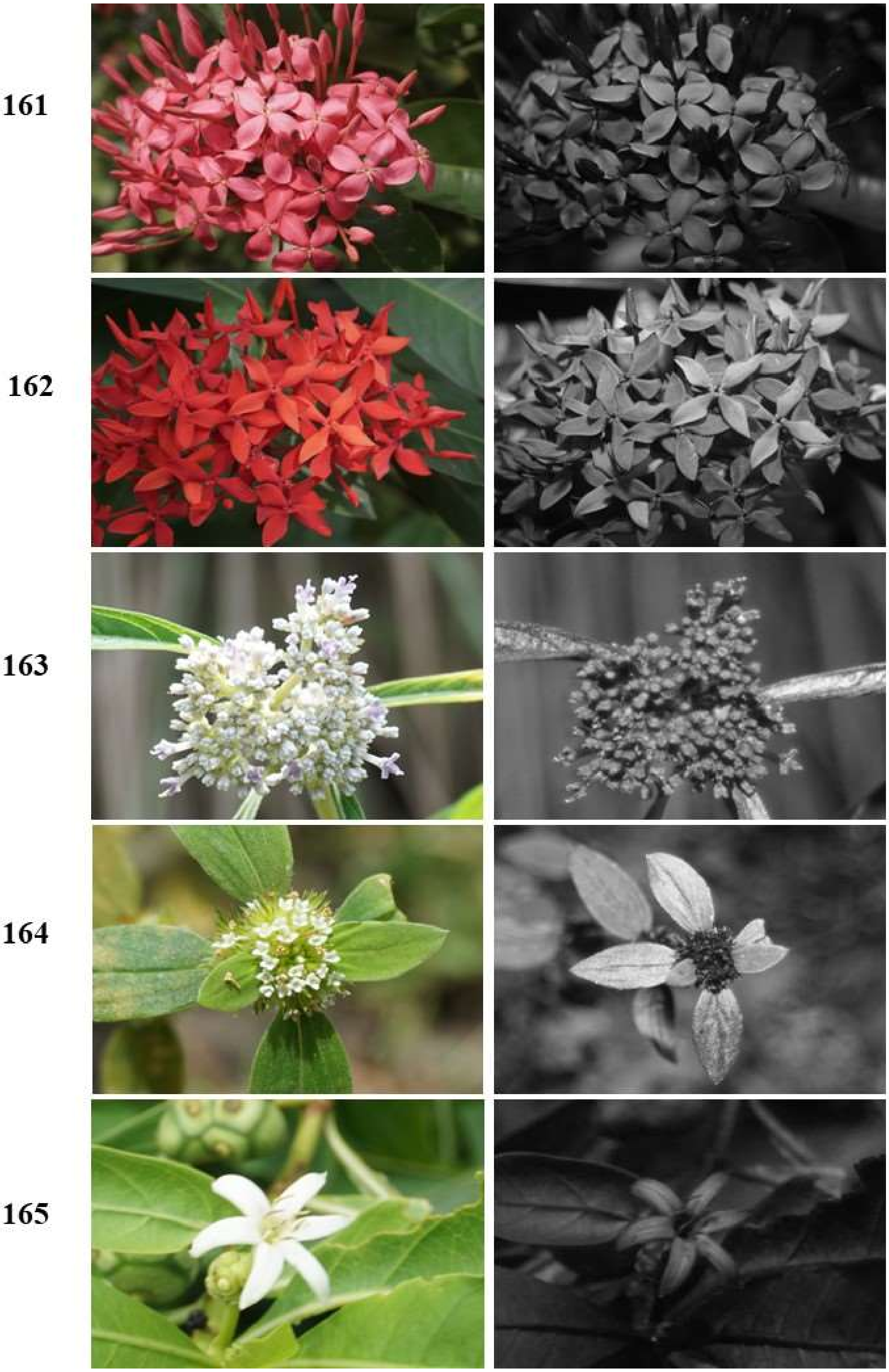


**Species list (cont.):** 161) *Ixora chinensis* Lam (Rubiaceae), 162) *Ixora coccinea* Curtis (Rubiaceae), 163) *Knoxia wightiana* Wall. ex Wight & Arn (Rubiaceae), 164) *Mitracarpus hirtus* (L.) DC (Rubiaceae), and 165) *Morinda coreia* Buch.-Ham. (Rubiaceae).

**166**

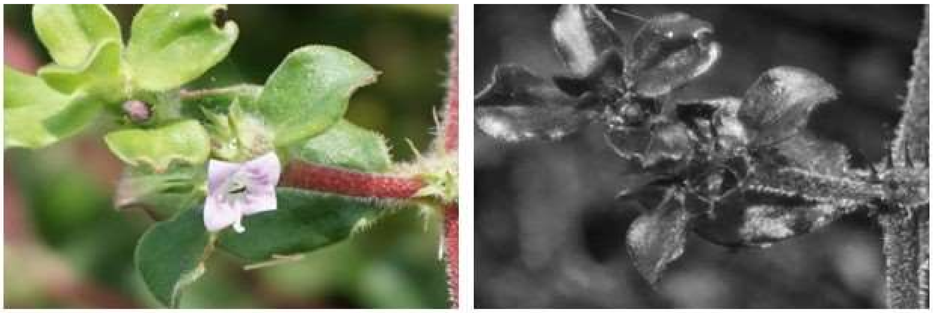

**167**

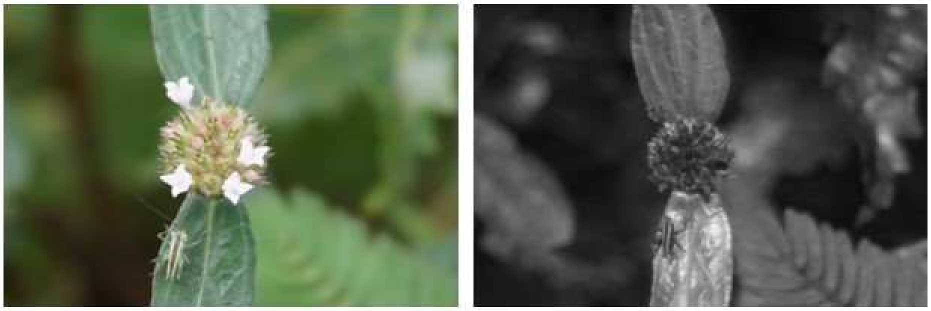

**168**

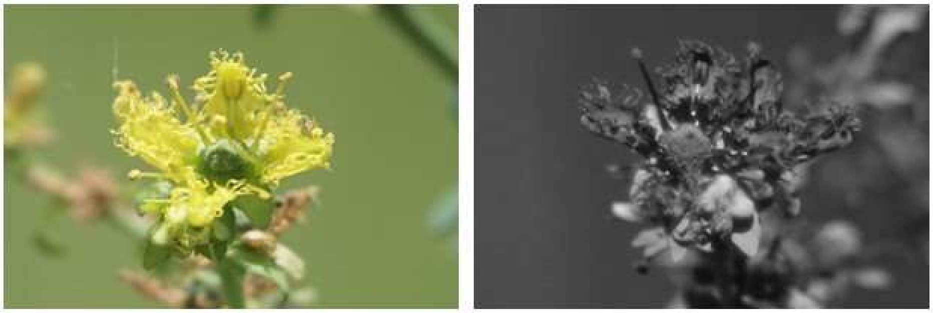

**169**

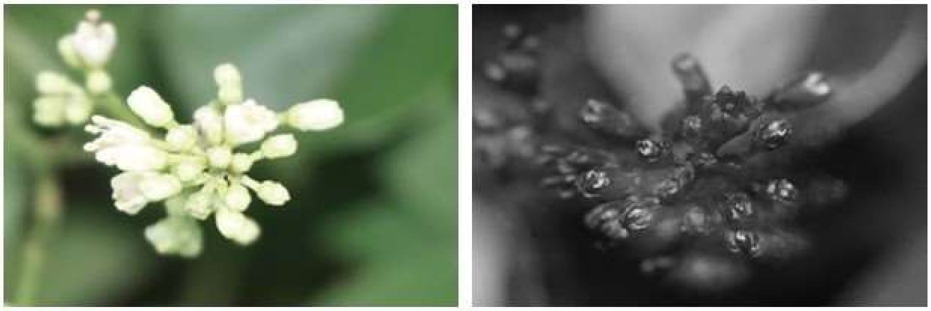

**170**

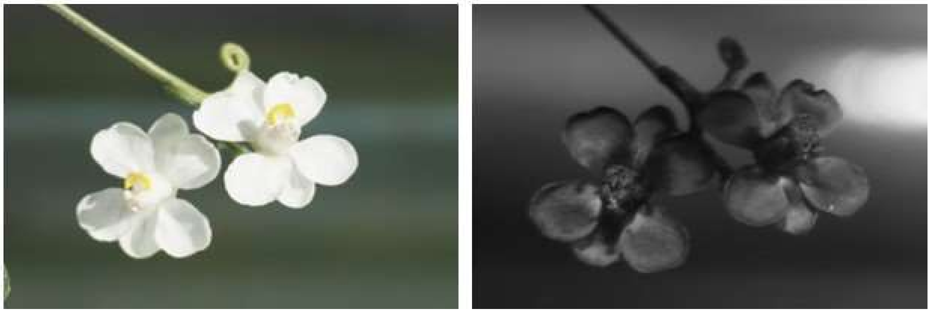

**Species list (cont.):** 166) *Spermacoce hispida* L. (Rubiaceae), 167) *Spermacoce remota* Lam. (Rubiaceae), 168) *Ruta graveolens* L. (Rutaceae), 169) *Toddalia asiatica* (L.) Lam. (Rutaceae), and 170) *Cardiospermum canescens* Wall. (Sapindaceae).

171

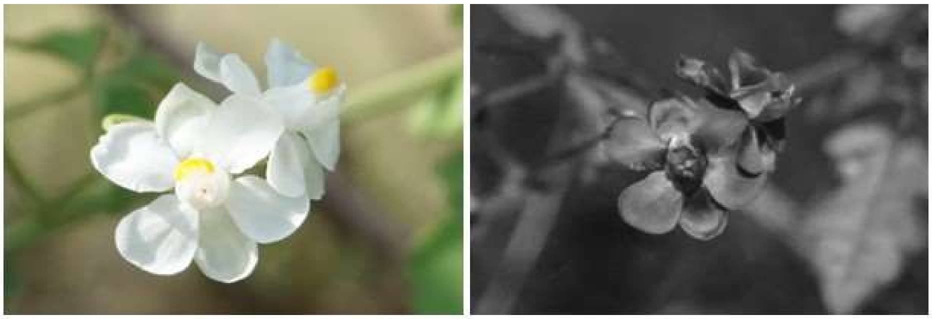

172

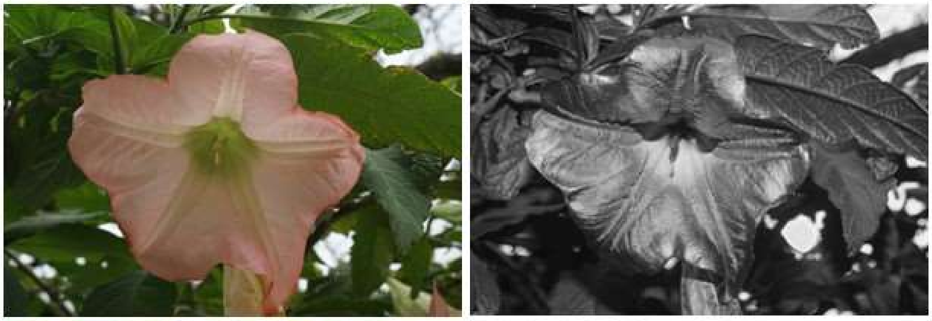

173

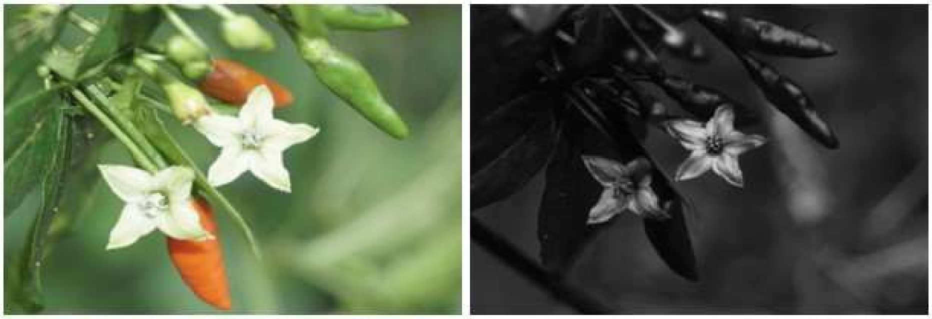

174

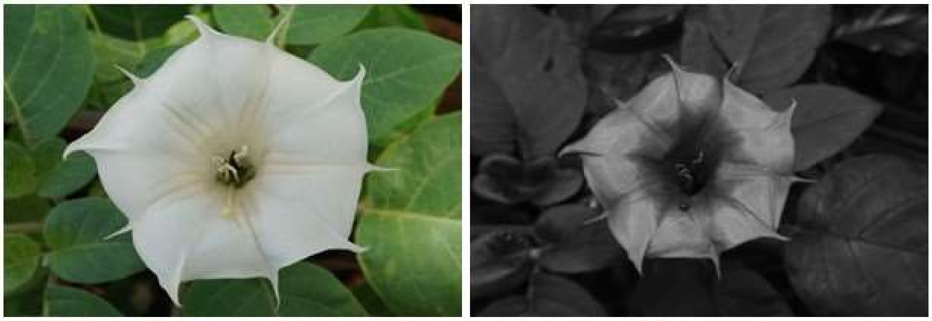

175

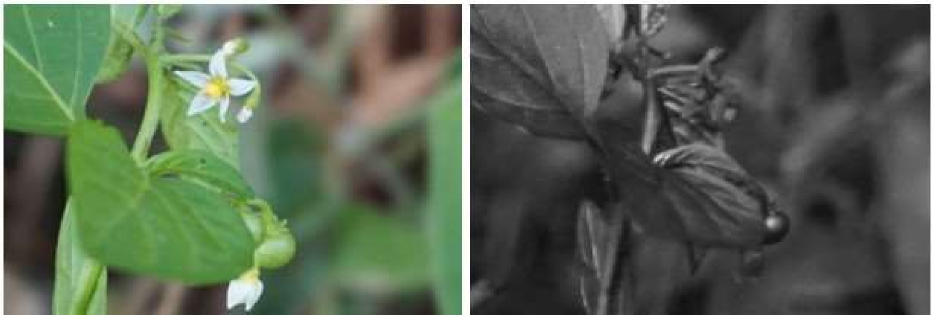

**Species list (cont.):** 171) *Cardiospermum corindum* L. (Sapindaceae), 172) *Brugmansia suaveolens* (Humb. & Bonpl. ex Willd.) Sweet (Solanaceae), 173) *Capsicum frutescens* L. (Solanaceae), 174) *Datura innoxia* Mill. (Solanaceae), and 175) *Solanum americanum* Mill. (Solanaceae).

176

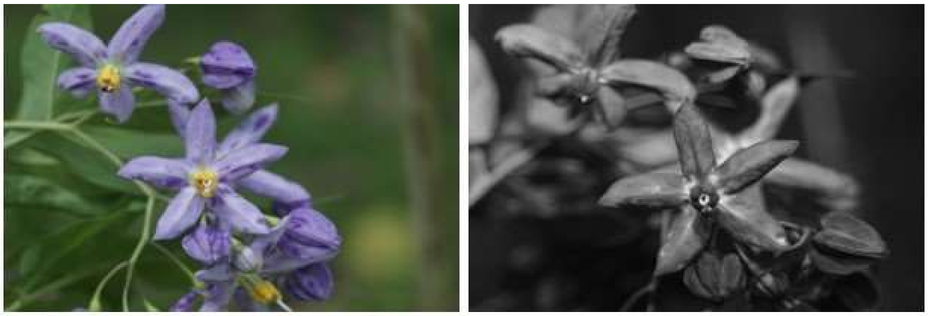

177

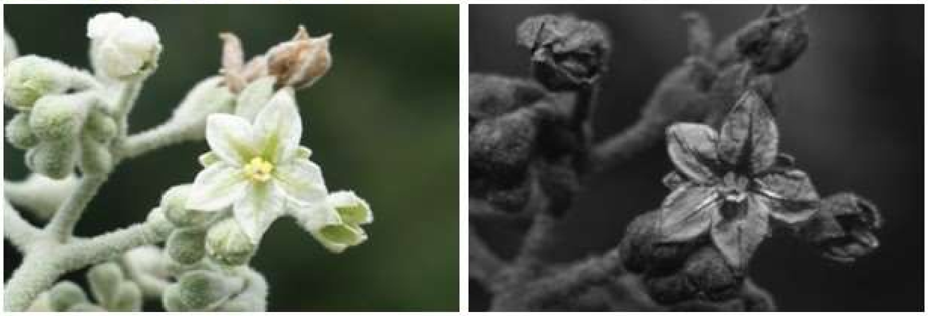

178

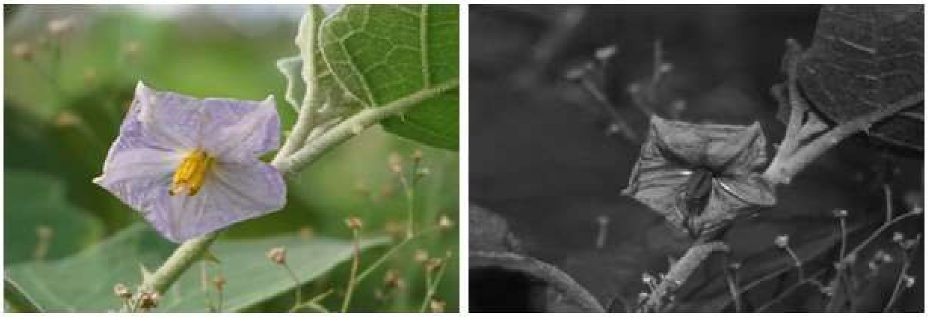

179

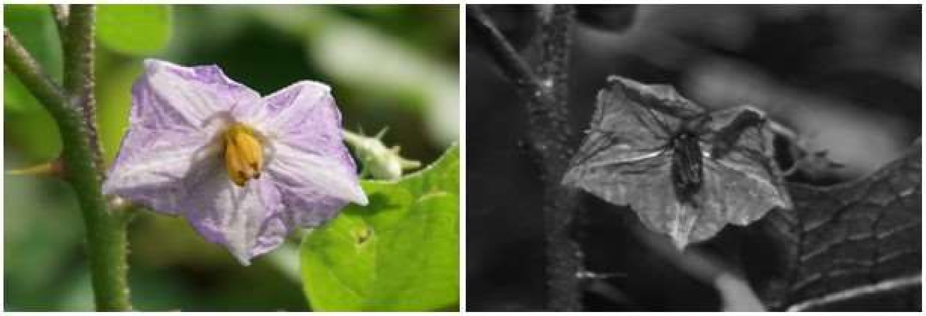

180

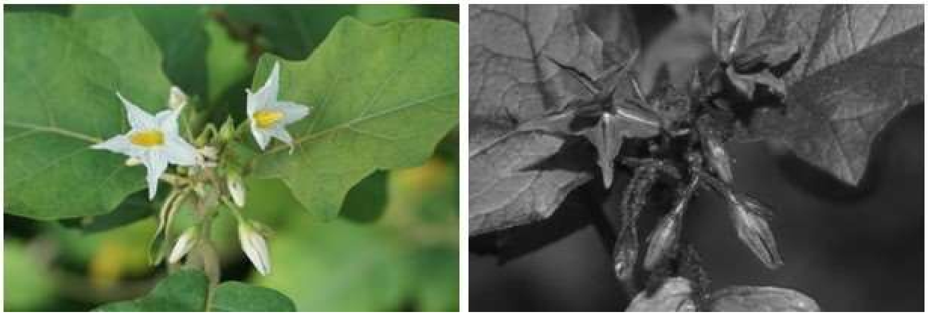

**Species list (cont.):** 176) *Solanum crispum* Ruiz & Pav. (Solanaceae), 177) *Solanum erianthum* D.Don (Solanaceae), 178) *Solanum melongena* L. (Solanaceae), 179) *Solanum melongena var. insanum* (L.) Filov (Solanaceae), and 180) *Solanum torvum* Buch.-Ham. ex Wall. (Solanaceae).

**181**

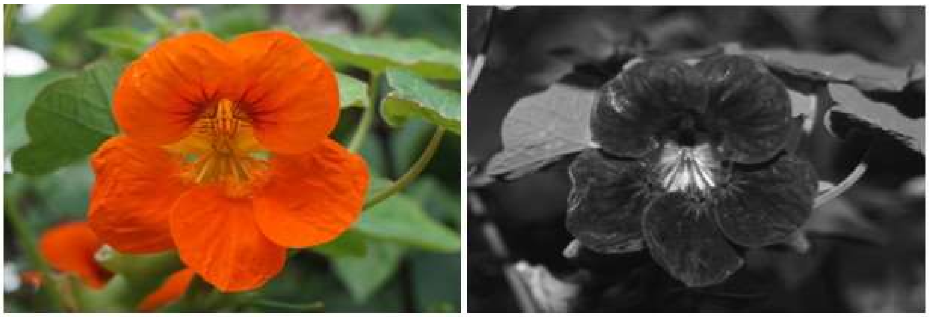

**182**

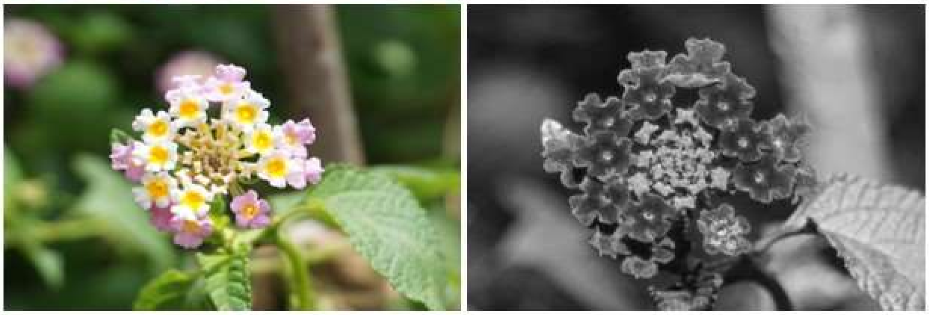

**183**

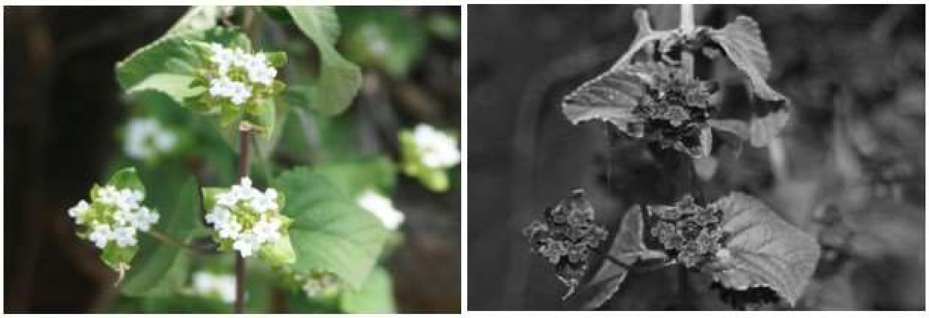

**184**

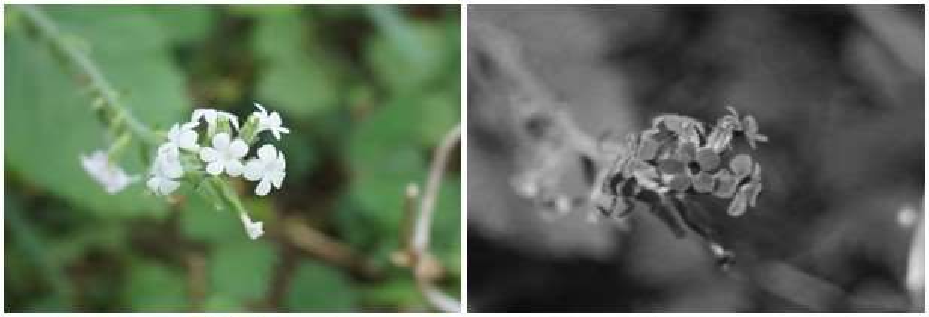

**185**

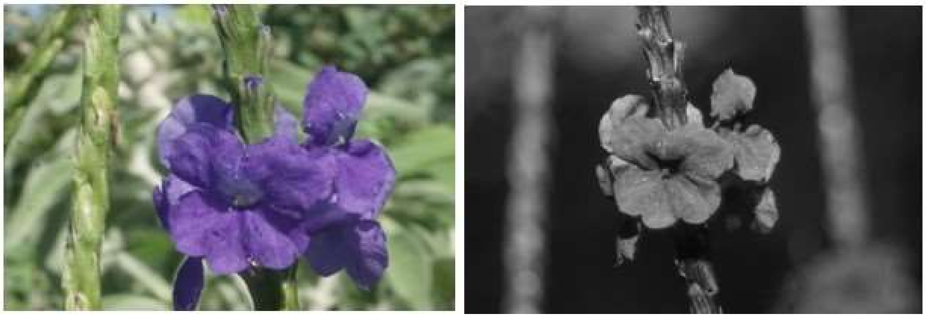

**Species list (cont.):** 181) *Tropaeolum majus* L. (Tropaeolaceae), 182) *Lantana camara* L. (Verbenaceae), 183) *Lantana veronicifolia* Hayek (Verbenaceae), 184) *Priva cordifolia* (L.f.) Druce (Verbenaceae), and 185) *Stachytarpheta cayennensis* (Rich.) Vahl (Verbenaceae).

**186**

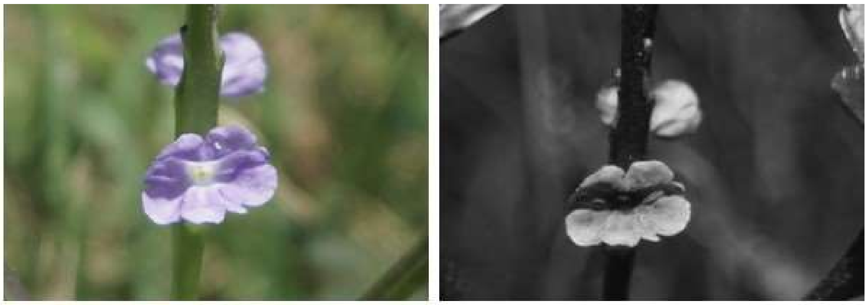

**187**

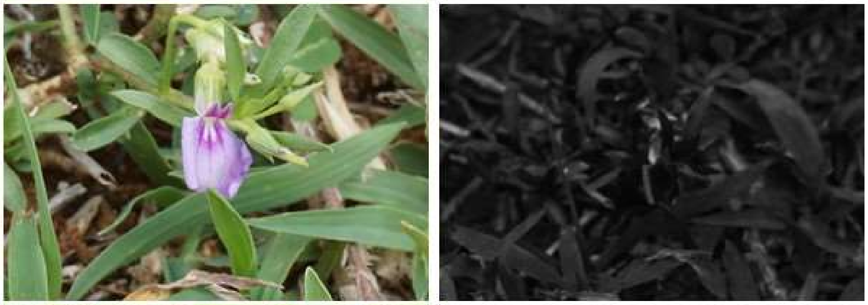

**188**

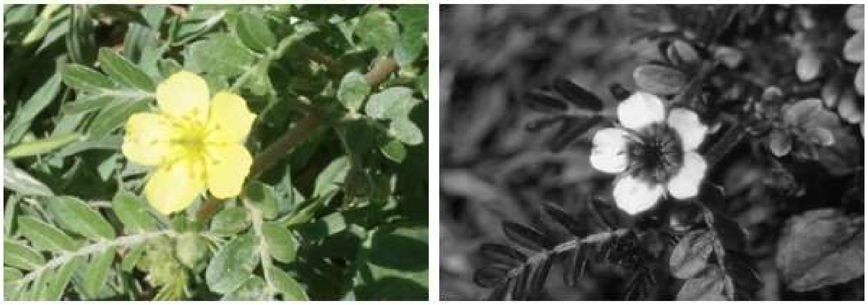

**Species list (cont.):** 186) *Stachytarpheta jamaicensis* (L.) Vahl (Verbenaceae), 187) *Hybanthus enneaspermus* (L.) F.Muell. (Violaceae), and 188) *Tribulus terrestris* L. (Zygophyllaceae).
